# Inferring the landscape of recombination using recurrent neural networks

**DOI:** 10.1101/662247

**Authors:** Jeffrey R. Adrion, Jared G. Galloway, Andrew D. Kern

**Author notes:** These authors contributed equally to this work (Jeffrey R. Adrion, Jared G. Galloway).

## Abstract

Accurately inferring the genome-wide landscape of recombination rates in natural populations is a central aim in genomics, as patterns of linkage influence everything from genetic mapping to understanding evolutionary history. Here we describe ReLERNN, a deep learning method for estimating a genome-wide recombination map that is accurate even with small numbers of pooled or individually sequenced genomes. Rather than use summaries of linkage disequilibrium as its input, ReLERNN takes columns from a genotype alignment, which are then modeled as a sequence across the genome using a recurrent neural network. We demonstrate that ReLERNN improves accuracy and reduces bias relative to existing methods and maintains high accuracy in the face of demographic model misspecification, missing genotype calls, and genome inaccessibility. We apply ReLERNN to natural populations of African *Drosophila melanogaster* and show that genome-wide recombination landscapes, while largely correlated among populations, exhibit important population-specific differences. Lastly, we connect the inferred patterns of recombination with the frequencies of major inversions segregating in natural *Drosophila* populations.

## Introduction

Recombination plays an essential role in the meiotic production of gametes in most sexual species, and is often required for proper segregation (***Nicklas, 1974***) and pairing of homologous chromosomes (reviewed in ***Zickler and Kleckner, 2015***). During prophase of meiosis, recombination is initiated by the formation of double-strand breaks (DSBs) across a wide array of organisms (***Lichten, 2001***). A subset of these DSBs will be repaired as crossover events, leading to reciprocal exchange between homologs. Those that are not resolved as crossovers are repaired through a number of mechanisms included noncrossover gene conversions and non-homologous end joining (***Do et al., 2014***). Recombination not only plays a central role in meiosis, but so too does it have wide ranging effects on both evolutionary and population genomics (***Lewontin and Kojima, 1960; Hill and Robertson, 1966; Ohta and Kimura, 1969, 1970; Smith and Haigh, 1974***).

Indeed, the population recombination rate *ρ* = 4*Nr* is a central parameter in population and statistical genetics (reviewed in ***Hahn, 2018***), as at equilibrium we expect *ρ* to be proportional to the scale of of linkage disequilibrium (LD) in a given region of the genome (***Ohta and Kimura, 1969***). In regions of the genome where *ρ* is relatively small we expect increased levels of LD, and conversely in genomic compartments with high *ρ* we expect little LD. Deviations from our expected levels of LD given the local recombination rate can be illustrative of the influence of other evolutionary forces such as selection or migration. For example, selective sweeps are expected to dramatically elevate LD near the target of selection (e.g. ***Kim and Nielsen, 2004; O’Reilly et al., 2008; Parsch et al., 2001***).

Structural variation itself is expected to modulate the landscape of recombination—herein the map of per-base recombination rates, *r*, to genomic positions along the chromosomes—as both crossovers and non-crossovers are predicated on the alignment of homologous sequences, and structural rearrangements may directly impact those alignments. Chromosomal inversions, long-known to suppress crossing over along a chromosome (e.g. ***Sturtevant, 1921***), are one of the best studied examples of such structural variation. Inversion polymorphisms have been implicated in diverse evolutionary phenomena including local adaptation (***Dobzhansky, 1937; Kirkpatrick and Barton, 2006; Ayala et al., 2013***), reproductive isolation (***White, 1977; Ayala et al., 2013; Noor et al., 2001; Rieseberg, 2001***), and the maintenance of meiotic drive complexes (reviewed in ***Jaenike, 2001***). As suppressors of recombination, we expect *a priori* that segregating inversions should show distinct histories of recombination in comparison to standard karyotype chromosomes.

While recombination plays a central role in meiosis and reproduction, the frequency and distribution of crossovers along the chromosomes are themselves phenotypes that can evolve. Not only is there a long tradition of work demonstrating the conditions under which rates of recombination might change (***Fisher, 1930; Muller, 1932; Charlesworth, 1976; Barton, 1995; Otto and Barton, 1997***), but increasingly there is good empirical evidence that such changes do indeed occur in nature (reviewed in ***Ritz et al., 2017***). Importantly, recombination rate variation exists between species, between populations, and between sexes of the same species (males generally having shorter maps than females) (***Hinch et al., 2011; Kong et al., 2010; Singh et al., 2013; Winckler et al., 2005***). Yet while there is abundant variation in the rate of recombination within and between taxa, methods for accurately measuring this variation have historically involved painstaking experiments or large pedigrees. Thus genetics, as a field, seeks ever-improving tools for directly estimating recombination rates from sequence data, without relying on pedigree genotyping or other ancillary information.

Accordingly, there is a rich history of estimating *ρ* in population genetics, including efforts to obtain minimum bounds on the number of recombination events (***Hudson and Kaplan, 1985; Myers and Griffiths, 2003; Wiuf, 2002***), methods of moments estimators (***Hudson, 1987; Wakeley, 1997***), composite likelihood estimators (***Chan et al., 2012; Hudson, 2002; McVean et al., 2002***), and summary likelihood estimators (***Li and Stephens, 2003; Wall, 2000***). Recently, supervised machine learning methods for estimating *ρ* have entered the fray (***Gao et al., 2016; Lin et al., 2013***), and have proven to be competitive in accuracy with state-of-the-art composite likelihood methods such as LDhat (***McVean et al., 2002***) or LDhelmet (***Chan et al., 2012***), often with far less computing effort. These methods, taken *en masse* have uncovered interesting biology, for instance the characterization of recombination hotspots (***Myers et al., 2005***), and are well suited for large samples of high quality genome or genotype data.

To this end, we sought to develop a novel method for inferring rates of recombination directly from a sequence alignment through the use of deep learning. In recent years deep artificial neural networks (ANNs) have produced remarkable performance gains in computer vision (***Krizhevsky et al., 2012; Szegedy et al., 2015***), speech recognition (***Hinton et al., 2012***), natural language processing (***Sutskever et al., 2014***), and data preprocessing tasks such as denoising (***Vincent et al., 2008***). Perhaps most illustrative of the potential of deep learning is the remarkable success of convolutional neural networks (CNNs; ***Lecun et al., 1998***) on problems in image analysis. For example, prior to the introduction of CNNs to the annual ImageNet Large Scale Visual Recognition Challenge (***Krizhevsky et al., 2012***), no method had achieved an error rate of less than 25% on the ImageNet data set. In the years that followed, CNNs succeeded in reducing this error rate below 5%, exceeding human accuracy on the same tasks (***Russakovsky et al., 2015***).

In this study we focus our efforts on recurrent neural networks (RNNs), a promising network architecture for population genomics, which has proven adept for analyzing sequential data of arbitrary lengths (***Graves et al., 2013***). Unlike other machine learning methods, deep learning approaches do not require a predefined feature vector. When fed labeled training data (e.g. a set of genotypes simulated under a known recombination rate), these methods algorithmically create their own set of informative statistics that prove most effective for solving the specified problem. By training deep learning networks directly on sequence alignments, we allow the neural network to automatically extract informative features from the data without human supervision. Learning directly from a sequence alignment for population genetic inference has recently been shown to be possible using CNNs (***Chan et al., 2018; Flagel et al., 2018***), and as we show below, is also true for RNNs. Moreover, supervised deep learning methods, such as RNNs, can be trained directly on the types of missing data that often beset researchers investigating non-model organisms using traditional tools.

Here we introduce **Re**combination **L**andscape **E**stimation using **R**ecurrent **N**eural **N**etworks, an RNN-based method for estimating the genomic map of recombination rates directly from a genotype alignment. We found that ReLERNN is both highly accurate and out-performs competing methods at small sample sizes. We also show that ReLERNN retains its high accuracy in the face of demographic model misspecification, missing genotypes, and genome inaccessibility. Further, we present an extension to ReLERNN that takes as input allele frequencies estimated by pooled sequencing (Pool-seq), making ReLERNN the first software package to directly infer rates of recombination in Pool-seq data. These results suggest that ReLERNN has the potential to fill a much-needed role in the analysis of low-quality or sparse genomic data. We then apply ReLERNN to population genomic data from African samples of *Drosophila melanogaster*. We demonstrate that the landscape of recombination is largely conserved in this species, yet individual regions of the genome show marked population-specific differences. Finally, we found that chromosomal inversion frequencies directly impact the inferred rate of recombination, and we demonstrate that the role of inversions in suppressing recombination extends far beyond the inversion breakpoints themselves.

## Results

### ReLERNN: an accurate method for estimating the genome-wide recombination landscape

We developed ReLERNN, a new deep learning method for accurately predicting genome-wide per-base recombination rates from as few as four chromosomes. Briefly, ReLERNN provides an end-to-end inferential pipeline for estimating a recombination map from a population sample: it takes as input either a Variant Call Format (VCF) file or, in the case of ReLERNN for Pool-seq data, a vector of allele frequencies and genomic coordinates. ReLERNN then uses the coalescent simulation program, msprime (***Kelleher et al., 2016***), to simulate training, validation, and test data sets under either constant population size or an inferred population size history. Importantly, these simulations are parameterized to match the distribution of Watterson’s estimator, *θ*_*W*_, calculated from the empirical samples. ReLERNN trains a specific type of RNN, known as a Gated Recurrent Unit (GRU; ***Cho et al., 2014***), to predict the per-base recombination rate for these simulations, using only the raw genotype matrix and a vector of genomic coordinates for each simulation example (***Figure 1, Figure S1, Figure S2***). It then uses this trained network to estimate genome-wide per-base recombination rates for empirical samples using a sliding-window approach. ReLERNN can optionally estimate 95% confidence intervals around each prediction using a parametric bootstrapping approach, and it uses the predictions generated while bootstrapping to correct for inherent biases in the training process (see Materials and Methods; ***Figure S3***).

**Figure 1.**
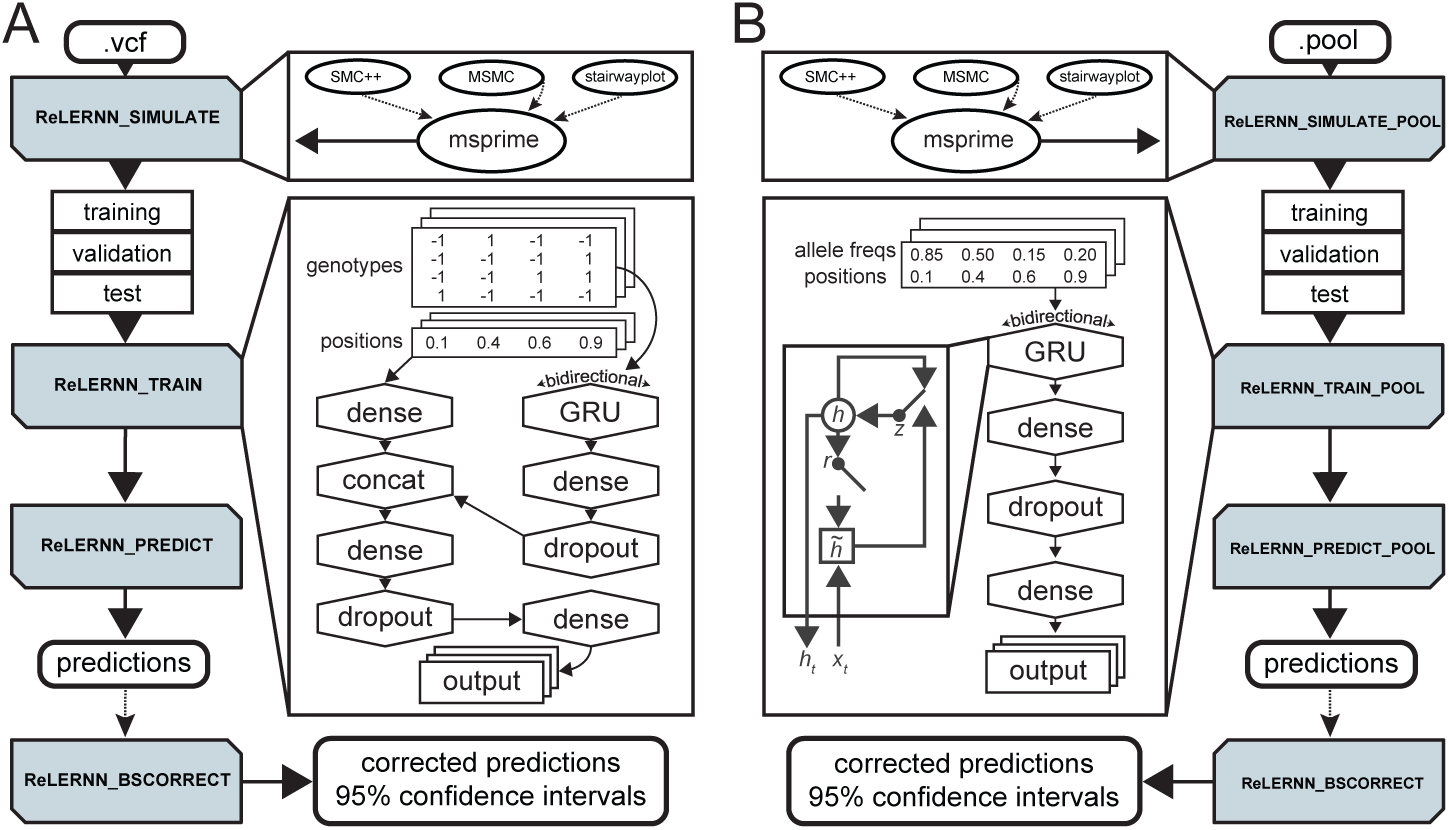
A cartoon depicting a typical workflow using ReLERNN’s four modules (shaded boxes) for **(A)** individually sequenced genomes or **(B)** pooled sequences. ReLERNN can optionally (dotted lines) utilize output from stairwayplot, SMC++, MSMC to simulate under a demographic history with msprime. Training inlays show the network architectures used, with the GRU inlay in **(B)** depicting the gated connections within each hidden unit. Here *r, z, h*_*t*_, and 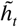 are the reset gate, update gate, activation, and candidate activation, respectively (***Cho et al., 2014***). The genotype matrix encodes alleles as reference (−1), alternative (1), or padded/missing data (0; *not shown*). Variant positions are encoded along the real number line (0-1).

A key feature of ReLERNN’s network architecture is the bidirectional GRU layer (***Figure 1, Figure S1***), which allows us to model genomic sequence alignments as a time series. While vanilla (feed-forward) networks use as input a full block of data for each example, recurrent layers break each genotype alignment into time steps corresponding to discrete genomic coordinates, and iterate over the time steps sequentially. At each time step, the gated recurrent units modulate the flow of information, using reset and update gates that control how the activation is updated (***Cho et al., 2014; Chung et al., 2014***). This process allows the gradient descent algorithm, known as backpropagation through time, to share parameters across time steps, as well as make inferences based on the ordering of SNPs—i.e. to have a spatial memory of allelic associations along the chromosome. The bidirectional attribute of the GRU layer simply means that each example is duplicated and reversed, so the sequence data are analyzed from both directions and then merged by concatenation. We present a generalized GRU for analyzing genomic sequence data, along with a more detailed look at the network architecture parameters used by ReLERNN in ***Figure S1***.

### Performance on simulated chromosomes

To assess our method we performed coalescent simulations using msprime (***Kelleher et al., 2016***), generating whole chromosome samples using a fine scale genetic map estimated from crosses of *D. melanogaster* (***Comeron et al., 2012***). We then used ReLERNN to estimate the landscape of recombination for these simulated examples. ReLERNN is able to predict the landscape of per-base recombination rates to a high degree of accuracy across a wide range of realistic parameter values, assumptions, and sample sizes (*R*^2^ ≥0.82; Mean absolute error (*MAE*) ≤ 1.28× 10^−8^). Importantly, the accuracy of ReLERNN is only modestly diminished when comparing predictions based on 20 samples (*R*^2^ = 0.93; *MAE* = 3.72 × 10^−9^; ***Figure 2*A****) to those based on four samples (*R*^2^ = 0.82; *MAE* = 6.66 × 10^−9^; ***Figure S4***). We also show that ReLERNN performs equally well on phased and unphased genotypes (*W* = 68.5; *P* = 0.17; Mann-Whitney *U* test; ***Figure S5***), suggesting that any effect of computational phasing error might be mitigated by treating the inputs as unphased variants.

**Figure 2.**
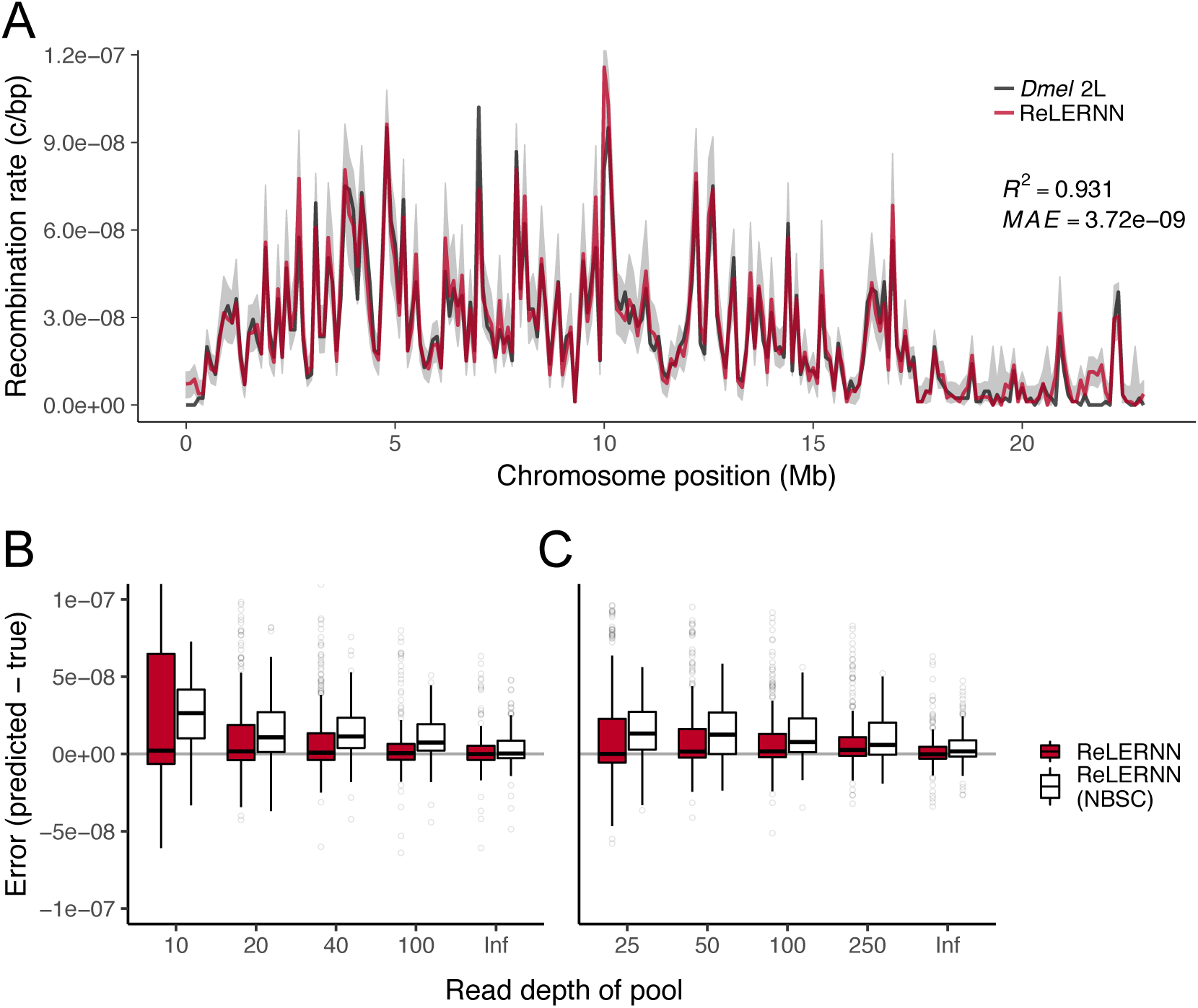
**(A)** Recombination rate predictions for a simulated *Drosophila* chromosome (black line) using ReLERNN for individually sequenced genomes (red line). The recombination landscape was simulated for *n* = 20 chromosomes under mutation-drift equilibrium using msprime (***Kelleher et al., 2016***), with per-base crossover rates taken from *D. melanogaster* chromosome 2L (***Comeron et al., 2012***). Gray ribbons represent 95% confidence intervals. *R*^2^ is reported for the general linear model of predicted rates on true rates and mean absolute error was calculated across all 100 kb windows. **(B)** Distribution of raw error (*r*_*predicted*_ − *r*_*true*_) using ReLERNN for Pool-seq data. Pools simulated from the same recombination landscape as above, with *n* = 20 and **(C)** *n* = 50 chromosomes across a range of simulated read depths (0.5*X* to 5*X*; *Inf* represents infinite simulated sequencing depth). Both the bootstrap-corrected predictions (red) and the non-bootstrap-corrected (NBSC; white) predictions are shown.

Because ReLERNN performed exceedingly well on unphased genotypes, we speculated that it might be able to glean crucial information about recombination rates from a vector of allele frequencies alone. Therefore we set out to extend ReLERNN to work with Pool-seq data, where the only inputs are a vector of allele frequencies and their corresponding genomic coordinates. Surprisingly, ReLERNN exhibits modest accuracy on simulated Pool-seq data, despite simulated sample and read depths as low as *n* = 50 and *coverage* = 50*X* (*R*^2^ = 0.54; *MAE* = 1.59×10^−8^; ***Figure S6***).

Increasing the read depth to a nominal 5*X* the sample depth (e.g. *n* = 50 and *coverage* = 250*X*) produced substantially greater accuracy (*R*^2^ = 0.69; *MAE* = 1.20 × 10^−8^; ***Figure S7***). As a general trend, we show that prediction error is reduced by increasing the number of chromosomes sampled in the pool (i.e. increasing allele frequency resolution) and by increasing the depth of sequencing (i.e. reducing sampling error) (***Figure 2*B****). While there currently exists software for estimating LD in Pool-seq data (***Feder et al., 2012***), to our knowledge ReLERNN is the first software to directly estimate rates of recombination using these data.

While ReLERNN retains accuracy at small sample sizes, it exhibits somewhat greater sensitivity to both the assumed genome-wide average mutation rate, 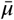, and the assumed maximum value for recombination, *ρ*_*max*_. To assess the degree of sensitivity to these assumptions, we ran ReLERNN on simulated chromosomes assuming 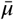 was both 50% greater and 50% less than the simulated mutation rate, *μ*_*true*_. In both scenarios, ReLERNN predicts crossover rates that are highly correlated with the true rates (*R*^2^ > 0.91). However, in both scenarios *MAE* is inflated but still modest, and the absolute rates of recombination are underpredicted (*R*^2^ = 0.91; *MAE* = 1.23 × 10^−8^; ***Figure S8***) and slightly overpredicted (*R*^2^ = 0.94; *MAE* = 1.28 × 10^−8^; ***Figure S9***) when assuming 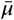 is less than or greater than *μ*_*true*_, respectively. Moreover, underestimating *ρ*_*max*_ causes ReLERNN to underpredict rates of recombination roughly proportional to the the magnitude of the underestimate (***Figure S10, Figure S11***), while overestimating *ρ*_*max*_ causes only a minor loss in accuracy (*R*^2^ = 0.90; *MAE* = 4.07 × 10^−9^; ***Figure S12***). Together these results suggest that ReLERNN is in fact learning information about the ratio of crossovers to mutations, and while ReLERNN is highly robust to errant assumptions when predicting relative recombination rates within a genome, caution must be taken when comparing absolute rates between organisms with large differences in per-base mutation rate estimates or for species. One additional limitation to ReLERNN it’s inability to fully resolve narrow recombination rate hot spots (herein defined as ≤ 10 kb genomic regions with *r* ≥ 50*X* the genome-wide average). We simulated hot spots of different lengths [*length* ∈ {2*kb*, 4*kb*, 6*kb*, 8*kb*, 10*kb*}, *r*_*background*_ = 2.5*e*^−9^, *r*_*hotspot*_ = 1.25*e*^−7^] and found that errors at hot spots were negatively correlated with hot spot length (***Figure S13***), suggesting that signal for crossovers at hot spots is being swamped by the background rate within the focal window, especially for very narrow hot spots relative to the focal window. This limitation could be of particular importance when attempting to resolve hot spots in human data, where lengths are often between 1-2 kb (***Jeffreys et al., 2001; Jeffreys and May, 2004***).

### ReLERNN compares favorably to competing methods, especially for small sample sizes and under model misspecification

To assess the accuracy of ReLERNN relative to existing methods, we took a comparative approach whereby we made predictions on the same set of simulated test chromosomes using methods that differ broadly in their approaches. Specifically, we chose to compare ReLERNN against two types of machine learning methods—a boosted regression method, FastEPRR (***Gao et al., 2016***), and a convolutional neural network (CNN) recently described in ***Flagel et al***. (***2018***)—and both LDhat (***McVean et al., 2002***) and LDhelmet (***Chan et al., 2012***), two widely cited approximate-likelihood methods. We independently simulated 10^5^ chromosomes using msprime (***Kelleher et al., 2016***) [parameters: *sample*_*size* ∈ {4, 8, 16, 32, 64}, *recombination*_*rate* = *U*(0.0, 6.25*e*^−8^), *mutation*_*rate* = *U*(1.875*e*^−8^, 3.125*e*^−8^), *length* = 3*e*^5^]. Half of these were simulated under demographic equilibrium and half were simulated under a realistic demographic model (based on the out-of-Africa expansion of European humans; see Materials and Methods). We show that ReLERNN outperforms all other methods, exhibiting significantly reduced absolute error (|*r*_*predicted*_ −*r*_*true*_|) under both the demographic model and under equilibrium assumptions (*T* ≤ −31; *P* < 10^−16^; *post hoc* Welch’s two sample *t*-tests for all comparisons; ***Figure S14, Figure S15***). ReLERNN also exhibited less bias than likelihood-based methods across a range of sample sizes (***Figure 3***), although all methods generally performed well at the largest sample size tested (*n* = 64).

**Figure 3.**
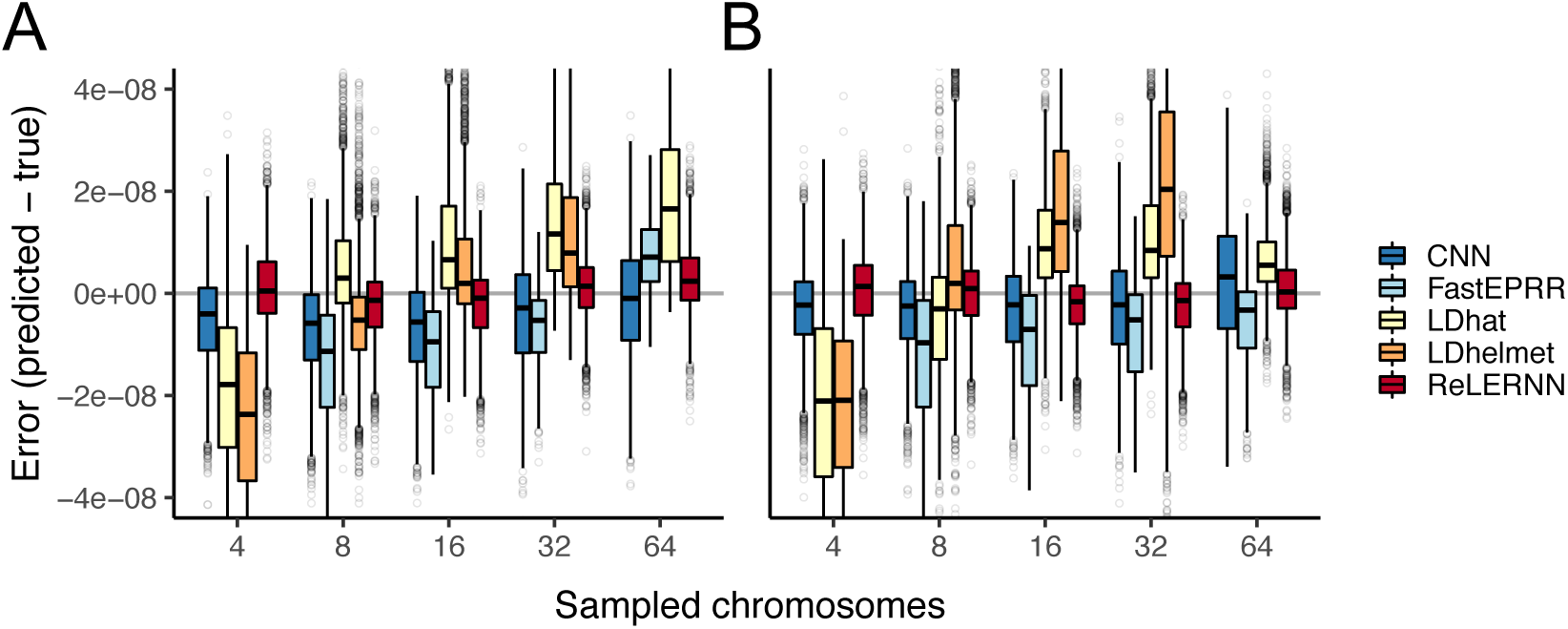
**(A)** Distribution of raw error (*r*_*predicted*_ − *r*_*true*_) for each method across 5000 simulated chromosomes (1000 for FastEPRR). Independent simulations were run under a model of population size expansion or **(B)** demographic equilibrium. Sampled chromosomes indicate the number of independent sequences that were sampled from each msprime (***Kelleher et al., 2016***) coalescent simulation. LDhelmet was not able be used with *n* = 64 chromosomes, and FastEPRR was not able to be used with *n* = 4.

We also sought to assess the robustness of ReLERNN to demographic model misspecification, where different generative models are used for simulating the training and test sets—e.g. training on assumptions of demographic equilibrium when the test data was generated by a population bottleneck. Methods robust to this type of misspecification are crucial, as the true demographic history of a sample is often unknown and methods used to infer population size histories can disagree or be unreliable (see ***Figure S21***). Moreover, population size changes alter the landscape of LD across the genome (e.g ***Slatkin, 1994; Rogers, 2014***), and thus have the potential to reduce accuracy or produce biased recombination rate estimates.

To this end, we trained ReLERNN on examples generated under equilibrium and made predictions on 5000 chromosomes generated by the human demographic model specified above (and also carried out the reciprocal experiment; ***Figure 4***). We compared ReLERNN to the CNN, LDhat, and LDhelmet, with all methods similarly misspecified (see Materials and Methods). We found that ReLERNN outperforms these methods under nearly all conditions, exhibiting significantly lower absolute error under both directions of demographic model misspecification (*T* ≤ −26; *P*_*WTT*_ < 10−16 for all comparisons, with the exception of the comparison to LDhelmet using 16 chromosomes; ***Figure S16, Figure S17***). We show that the error directly attributed to model misspecification (which we term marginal error; see Materials and Methods) is occasionally higher in ReLERNN relative to other methods, even though ReLERNN exhibited the lowest absolute error among methods. As a prime example of this, we found predictions from LDhelmet were not affected by our misspecification regime at all, but these predictions were still, on average, less accurate than those made by a misspecified ReLERNN. Interestingly, marginal error is significantly greater when ReLERNN was trained on equilibrium simulations and tested on demographic simulations than under the reciprocal misspecification (*T* = 26.3; *P*_*WTT*_ < 10^−16^; ***Figure S18***). While this is true, it is important to note that mean marginal error for ReLERNN, in both directions of misspecification and across all sample sizes, never exceeded 3.90 × 10^−9^, suggesting that the additional information gleaned from an informative demographic model is limited.

**Figure 4.**
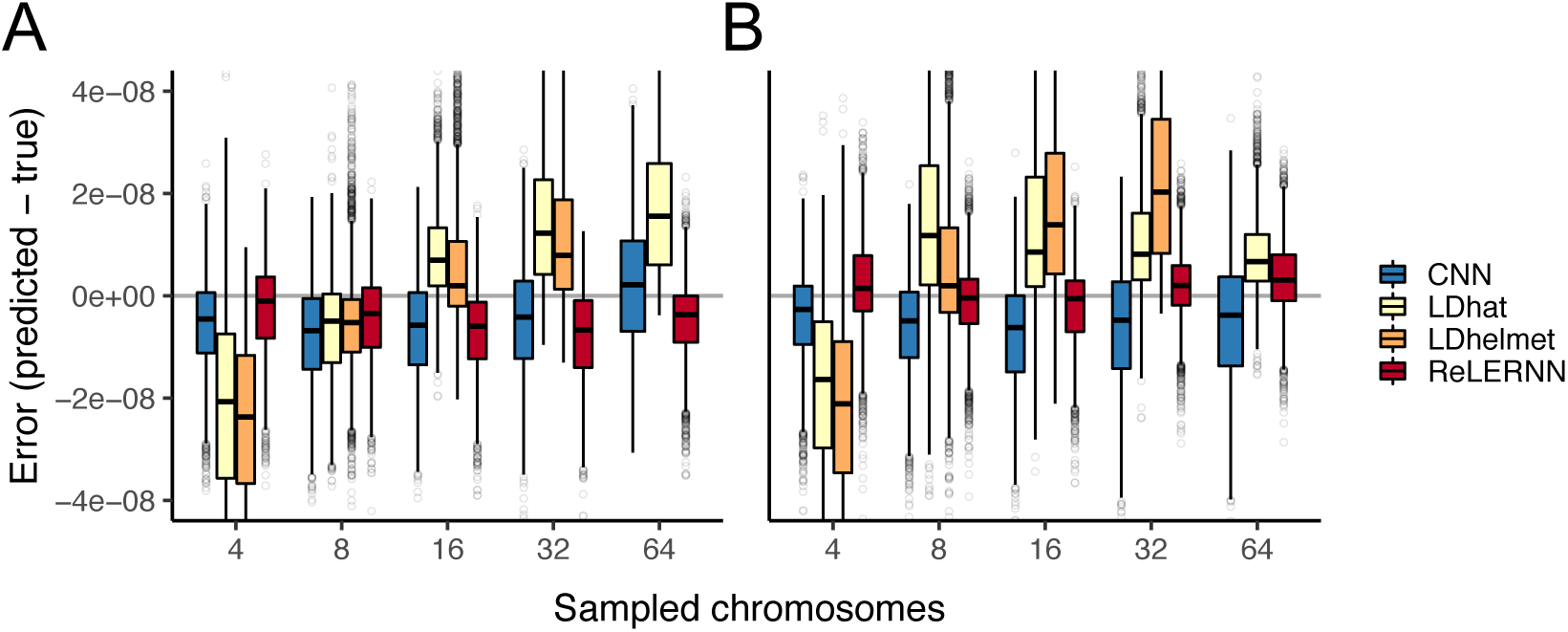
**(A)** Distribution of raw error (*r*_*predicted*_ − *r*_*true*_) for each method across 5000 simulated chromosomes after model misspecification. For the CNN and ReLERNN, predictions were made by training on equilibrium simulations while testing on sequences simulated under a model of population size expansion or **(B)** training on demographic simulations while testing on sequences simulated under equilibrium. For LDhat and LDhelmet, the lookup tables were generated using parameters values that were estimated from simulations where the model was misspecified in the same way as described for the CNN and ReLERNN above. Sampled chromosomes indicate the number of independent sequences that were sampled from each msprime (***Kelleher et al., 2016***) coalescent simulation. LDhelmet was not able be used with *n* = 64 chromosomes and the demographic model could not be intentionally misspecified using FastEPRR.

In addition to model misspecification, differences in the ratio of homologous gene conversion events to crossovers can also bias the inference of recombination rates, as conversion tracts break down LD within the prediction window (***Gay et al., 2007; Przeworski and Wall, 2001***). We treated the effect of gene conversion as another form of model misspecification, by training on examples that lacked gene conversion and testing on examples that included gene conversion. As ReLERNN uses msprime for all training simulations, and msprime cannot currently simulate gene conversion, we generated all test set simulations with ms (***Hudson, 2002***). We found that including gene conversion in our simulations biased our predictions, resulting in an overestimate of the true recombination rate (***Figure S19***). Moreover, the magnitude of this bias increased with the ratio of gene conversion events to crossovers, 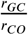. As expected, we also observed a similar pattern of bias for LDhelmet, although the magnitude of bias for LDhelmet was less than that exhibited by ReLERNN for 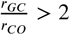 (*T* > 4.37; *P*_*WTT*_ < 1.32× 10^−5^; ***Figure S19***). As errors in genotype calls can mimic gene conversion—e.g. a heterozygous sample being called as a homozygote—filtering low-quality SNP calls, either by removing the individual genotype or by masking sites, has the potential to mitigate gene conversion-induced bias. However, missing genotypes and inaccessible sites have the potential to introduce their own biases, highlighting an area where deep learning methods may have a unique advantage over traditional tools.

### ReLERNN retains high accuracy on simulated low-quality genomic datasets

Deep learning tools have the potential to perform exceptionally well on poor-quality genomic datasets, such as those with low-quality or low-complexity reference genomes, under sampling regimes where individual samples are at a premium, or where base- and map-quality scores are suspect. This is in part because such attributes of genomic quality can be readily incorporated during training, and deep learning methods can generalize despite these limitations. To address the potential for ReLERNN to serve as an asset for researchers working with low-quality data—e.g. those studying non-model organisms—we simulated 1 Mb chromosomes under a randomized fine-scale recombination landscape, and then masked increasing fractions of both genotypes and sites. We then trained ReLERNN with both missing genotypes and genome inaccessibility, and generated predictions on the simulated chromosomes.

We show that ReLERNN exhibits high accuracy and low bias on datasets with missing genotypes, even as the faction of missing data increases to half of all genotypes (***Figure 5***). Moreover, we found that ReLERNN had reduced bias and significantly lower absolute error than LDhelmet at 50% missing genotypes for both *n* =4 and *n* = 20 (*T* ≤ −2.8; *P*_*WTT*_ < 0.007 for both comparisons). Here we define missing genotypes as any genotype call set to a. in the VCF, although in theory a simple quality threshold to identify missing genotypes could also be implemented. Additionally we tested ReLERNN across increasing levels of genome inaccessibility (up to 75% of all sites inaccessible), simulating a scenario where the vast majority of sites cannot be accurately mapped—e.g. in low-complexity genomic regions or for taxa without reference assemblies. Here genome inaccessibility refers to any site overlapping a window in the accessibility mask, where the entire genotype array at this site is discarded. Again, ReLERNN exhibited reduced bias in error across all levels of genome accessibility relative to LDhelmet (***Figure S20***). However, levels of absolute error were not significantly different between the methods after correcting for multiple tests (*T* ≤ −2.1; *P* _*WTT*_ ≥ 0.043 for all comparisons). Together these results suggest that ReLERNN may be of particular interest to researchers studying non-model organisms or for those without the access to high-quality reference assemblies.

**Figure 5.**
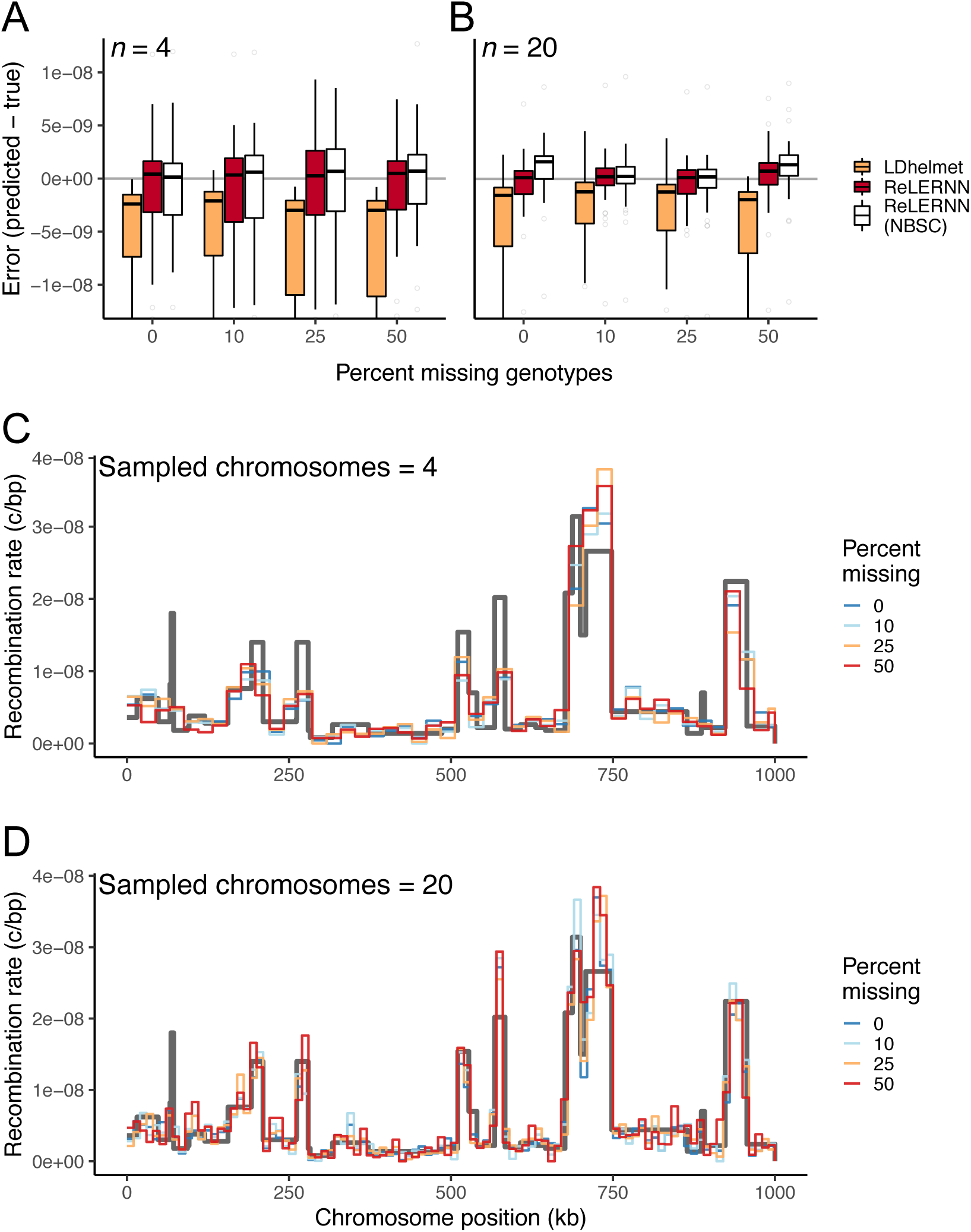
**(A)** Distribution of raw error (*r*_*predicted*_ − *r*_*true*_) for LDhelmet and ReLERNN when presented with varying levels of missing genotypes for simulations with *n* = 4 and **(B)** *n* = 20 chromosomes. **(C)** Fine-scale rate predictions generated by ReLERNN for a 1 Mb recombination landscape (grey line) simulated with varying levels of missing genotypes, for *n* =4 and **(D)** *n* = 20 chromosomes.

### Recombination landscapes are largely concordant among populations of African *D. melanogaster*

Using our method, we characterized the genome-wide recombination landscapes of three populations of African *D. melanogaster* (sampled from Cameroon, Rwanda, and Zambia). Each population was derived from the sequencing of 10 haploid embryos (detailed in ***Lack et al., 2015; Pool et al., 2012***), hence these data represent an excellent opportunity to exploit ReLERNN’s high accuracy on small sample sizes. The lengths of genomic windows selected by ReLERNN were roughly consistent among populations, and ranged from 38 kb for chromsomes 2R, 3L, and 3R in Zambia, to 51 kb for the X chromosome in Cameroon. We show that fine-scale recombination landscapes are highly correlated among all three populations of *D. melanogaster* (genome-wide mean pairwise Spearman’s *ρ* = 0.76; *P* < 10^−16^; 100 Kb windows; ***Figure 6***). The genome-wide mean pairwise coefficient of determination between populations was somewhat lower, *R*^2^ = 0.63 (*P* < 10^−16^; 100 Kb windows), suggesting there may be important population-specific differences in the fine-scale drivers of allelic association. These differences may also contribute to within-chromosome differences in recombination rate between populations. Indeed, we estimate that mean recombination rates are significantly different among populations for all chromosomes with the exception of chromosome 3L (*P* ≤ 3.78 × 10^−4^; one-way analysis of variance). Post-hoc pairwise comparisons suggest that this difference is largely driven by an elevated rate of recombination in Zambia, identified on all chromosomes (*P* ≤ 8.21× 10^−4^; Tukey’s HSD tests) except for 3L (*P*_*HSD*_ ≥ 0.15). ReLERNN predicts the recombination rate in simulated test sets to a high degree of accuracy for all three populations (*R*^2^ ≥ 0.93; *P* < 10^−16^; ***Figure S23***), suggesting that we have sufficient power to discern fine-scale differences in per-base recombination rates across the genome.

**Figure 6.**
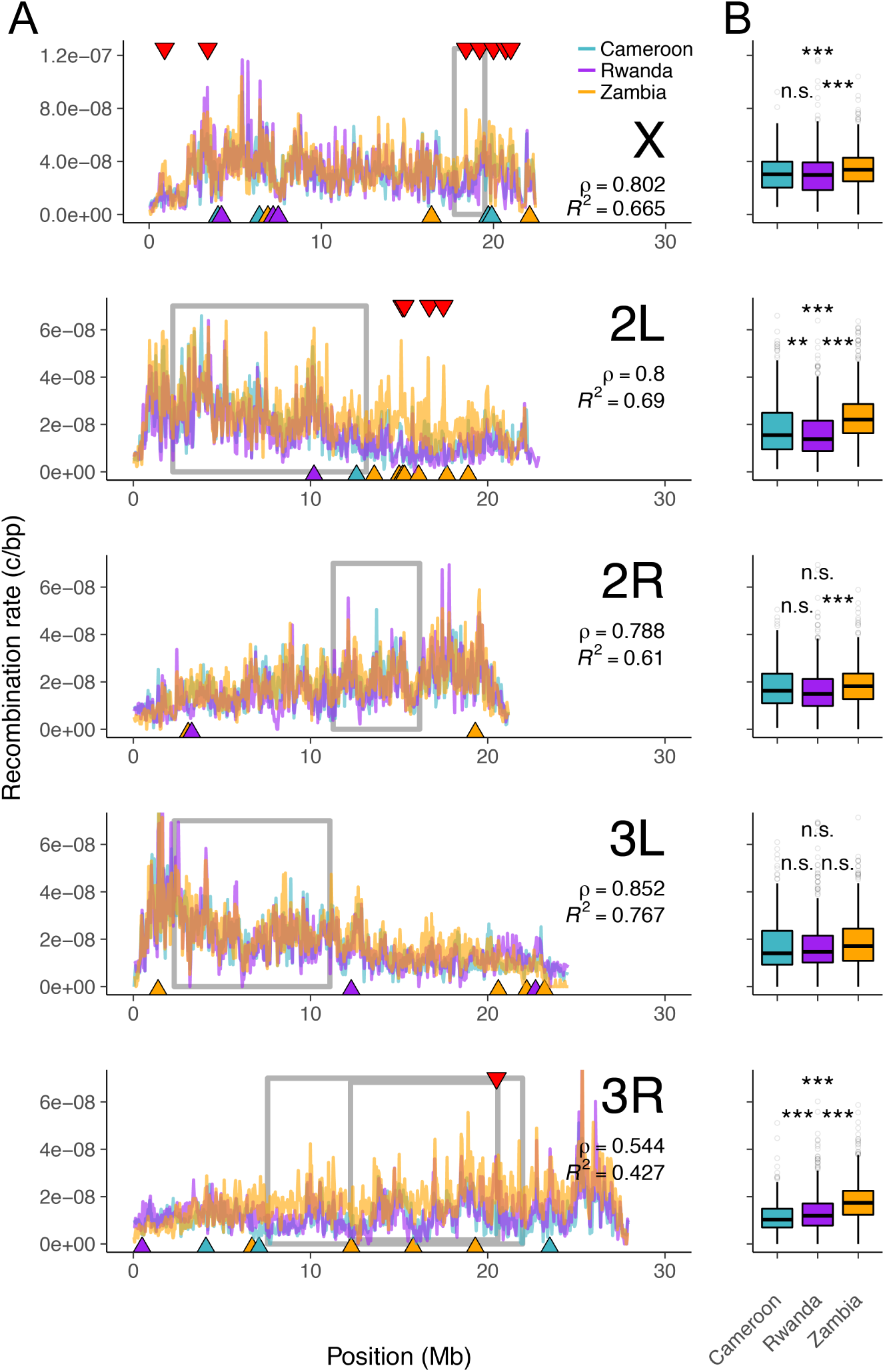
**(A)** Genome-wide recombination landscapes for *D. melanogaster* populations from Cameroon (teal lines), Rwanda (purple lines), and Zambia (orange lines). Grey boxes denote the inversion boundaries predicted to be segregating in these samples (***Pool et al., 2012; Corbett-Detig and Hartl, 2012***). Red triangles mark the top 1% of global outlier windows for recombination rate. Blue, purple, and orange triangles mark the top 1% of population-specific outlier windows for recombination rate, with triangle color indicating the outlier population (see Materials and Methods). **(B)** Per-chromosome recombination rates for each population. Spearman’s *ρ* and *R*^2^ are reported as the mean of pairwise estimates between populations for each chromosome. \*\**P* < 0.01 and \*\*\**P* < 0.001 are based on Tukey HSD tests for all pairwise comparisons.

When comparing our recombination rate estimates to those derived from experimental crosses of North American *D. melanogaster* (reported in ***Comeron et al., 2012***), we find that the coefficients of determination averaged over all three populations were *R*^2^ = 0.46, 0.70, 0.47, 0.08, 0.73 for chromosomes 2L, 2R, 3L, 3R, and X, respectively (***Figure S24***; 1 Mb windows). These results differ from those observed by ***Chan et al***. (***2012***), who compared 22 *D. melanogaster* sampled from the same Rwandan population to the FlyBase map and found *R*^2^ = 0.55, 0.63, 0.45, 0.42, 0.41 for the same chromosomes. The minor differences we observed between methods for chromosomes 2L, 2R, 3L, and the X chromosome can likely be attributed to the fact that we are comparing estimates from two different methods, using different African flies, to a different experimentally derived map. However, the larger differences found between methods for chromosome 3R seem less likely attributable to methodological differences. Importantly, African *D. melanogaster* are known to harbor large polymorphic inversions often at appreciable frequencies (***Lemeunier and Aulard, 1992; Aulard et al., 2002***). For example, the inversion *In(3R)K* segregates in our Cameroon population at *p* = 0.9. It is potentially these differences in inversion frequencies that contribute to the exceptionally weak correlation observed using our method for chromosome 3R.

An important cause of population-specific differences in recombination landscapes might be population-specific differences in the frequencies of chromosomal inversions, as recombination is expected to be strongly suppressed between standard and inversion arrangements. To test for an effect of inversion frequency inferences made by ReLERNN, we resampled haploid genomes from Zambia to create artificial population samples with the cosmopolitan inversion *In(2L)t* segregating at varying frequencies, *p* ∈ {0.0, 0.2, 0.6, 1.0}. In Zambia, *In(2L)t* arose recently (***Corbett-Detig and Hartl, 2012***) and segregates at *p* = 0.22 (***Lack et al., 2015***), suggesting that recombination within the inversion breakpoints may be strongly suppressed in individuals with the inverted arrangement relative to those with the standard arrangement. For these reasons, we predict that the inferred recombination rate should decrease as the low-frequency inverted arrangement is increasingly overrepresented in the set of sampled chromosomes (i.e. as more of the samples contain the high-LD inverted arrangements). As predicted, we found a strong effect of the sample frequency of *In(2L)t* on estimated rates of recombination for chromosome 2L in Zambia (***Figure S27***), demonstrating that ReLERNN is sensitive to the frequency of recent inversions.

To further explore population-specific differences in recombination landscapes we took a statistical outlier approach, whereby we define two types of recombination rate outliers—global outliers and population-specific outliers (see Materials and Methods). Global outliers are characterized by windows with exceptionally high variance in rates of recombination between all three populations (***Figure 6***; red triangles) while population-specific outliers are those windows where the rate of recombination in one population is strongly differentiated from the rates in the other two populations (***Figure 6***; population-colored triangles). We find that population-specific outliers, but not global outliers, are significantly enriched within inversions (*P* = 0.005; randomization test; ***Figure 6***; grey boxes). Moreover, this enrichment remains significant when extending the inversion boundaries by up to 250 Kb (*P*_*rand*_ ≤ 0.004). However, extending the inversion boundaries beyond 250 Kb, or restricting the overlap to windows surrounding only the breakpoints (250 Kb, 500Kb, 1 Mb, 2 Mb), erodes this pattern (*P*_*rand*_ ≥ 0.055 for all comparisons), suggesting that the role for inversions in generating population-specific differences in recombination rates is complex, at least for these populations.

Selection is another important factor that may confound the inference of recombination rates. For instance selective sweeps generate localized patterns of high LD on either side of the sweep site (***Kim and Nielsen, 2004; Schrider et al., 2015***), thus regions flanking selective sweeps may mimic regions of reduced recombination. Inasmuch population-specific selective sweeps are expected to contribute to population-specific differences in recombination rate estimates. We used diploS/HIC (***Kern and Schrider, 2018***) to identify hard and soft selective sweeps in our African *D. melanogaster* populations, and we tested for an excess of recombination rate outliers overlapping with windows classified as sweeps. In total, diploS/HIC classified 27.4%, 28.1%, and 26.8%, of all genomic widows as selective sweeps (either “hard” or “soft”) for Cameroon, Rwanda, and Zambia, respectively, when looking at 5kb, non-overlapping windows. The associated False Discovery Rates (FDR) for calling sweeps in these populations were appreciable: 33.9%, 33.1% and 34.7%, respectively (***Figure S26***). As expected, windows classified as sweeps had significantly lower rates of recombination relative to neutral windows in all three populations (*P*_*WTT*_ ≤ 10^−16^ for all comparisons; ***Figure S25***). However, we found that neither global nor population-specific outliers were enriched for selective sweeps (*P*_*rand*_ ≥ 0.246 for both comparisons), suggesting that, when treated as a class, recombination rate outliers are not likely driven by sweeps in these populations. When treated separately (i.e. independent permutation tests for each recombination rate outlier window), we identified 7 outliers enriched for sweeps at the *P* ≤ 0.05 threshold, corresponding to an expected FDR of 77%. However, given our FDR for calling sweeps in these populations, our measure of the enrichment in overlap with recombination rate outliers is likely to be conservative. Two of these outlier windows may represent potential true positives; an outlier in Cameroon contains 5 out of 6 non-overlapping 5 kb windows classified as “hard” sweeps, the second from Rwanda has 10 out of 12 windows classified as “hard” sweeps (*P*_*rand*_ = 0.0 for both comparisons). These two recombination rate outlier windows are potentially ripe for future studies on selective sweeps in these populations, and suggest that in at least some instances, selection contributes to observed differences in estimates of recombination rates between *Drosophila* populations.

## Discussion

We introduced a new method, ReLERNN, for predicting the genome-wide map of per-base recombination rates from polymorphism data, through the use of deep neural networks. Importantly, ReLERNN is particularly well-suited to take advantage of emerging small-scale sequencing experiments—e.g. those traditionally associated with the study of non-model organisms. Population genomics, as a field, relies on estimates of recombination rates to understand the effects of diverse phenomena ranging from the impacts of natural selection (***Elyashiv et al., 2016***), to patterns of admixture and introgression (***Price et al., 2009; Brandvain et al., 2014; Schumer et al., 2018***), to polygenic associations in genome-wide association studies (***Bulik-Sullivan et al., 2015***). As befits this need, there has been a long tradition of development of statistical methods for estimating the population recombination parameter, *ρ* = 4*Nr* (***Chan et al., 2012; Gao et al., 2016; Hudson and Kaplan, 1985; Hudson, 1987, 2002; Li and Stephens, 2003; Lin et al., 2013; McVean et al., 2002; Myers and Griffiths, 2003; Wakeley, 1997; Wall, 2000; Wiuf, 2002***).

We sought to harness the power of deep learning, specifically deep recurrent neural networks, to address the problem of estimating recombination rates, and in so doing, we developed a workflow that reconstructs the genome-wide recombination landscape to a high degree of accuracy from very small sample sizes—e.g. four haploid chromosomes or directly from allele frequencies obtained through Pool-seq. The use of deep learning has recently revolutionized the fields of computer vision (***Krizhevsky et al., 2012; Szegedy et al., 2015***), speech recognition (***Hinton et al., 2012***), and natural language processing (***Sutskever et al., 2014***), and while its use in population genomics has only recently begun, it is anticipated to be similarly fruitful (***Schrider and Kern, 2018***). The natural extension of deep learning to population genomic analyses comes as a result of the ways in which ANNs learn abstract representations of their inputs. In the case of population genomic analyses, the inputs can be naturally represented as DNA sequence alignments, eliminating the need for human oversight (and potentially constraint) in the form of statistical summaries (i.e. compression) of the raw data. ANNs can then learn high-dimensional statistical associations directly from the sequence alignments, and use these to return highly accurate predictions.

ReLERNN utilizes a variant of an ANN, known as a Gated Recurrent Unit (GRU), as its primary technology. GRU networks excel at identifying temporal associations (***Jozefowicz et al., 2015***), and therefore we modeled our sequence alignment as a bidirectional time series, where each ordered SNP represented a new time step along the chromosome. We also modeled the distance between SNPs using a separate input tensor, and these two inputs were concatenated after passing through the initial layers of the network (see ***Figure 1*** inlay). We demonstrated that ReLERNN can predict a simulated recombination landscape with a high degree of accuracy (*R*^2^ = 0.93; ***Figure 2***), and that these predictions remain high, even when using small sample sizes (*R*^2^ = 0.82; ***Figure S4***). These predictions compared favorably to those made by a leading composite likelihood methods (LDhat and LDhelmet; ***McVean et al., 2002; Chan et al., 2012***), as well as other machine learning methods (the CNN and FastEPRR; ***Figure 3***).

We also showed that ReLERNN can achieve modest accuracy when presented solely with allele frequencies derived from simulated Pool-seq data, especially when sequenced at the relatively modest depth of 5*X* the pool size (***Figure S7***). Moreover, ReLERNN performed well at estimating recombination rates in the face of missing genotype calls—exhibiting reduced bias when compared to LDhelmet, even with 50% of genotypes missing (***Figure 5***) or 75% of the genome inaccessible to SNP calls (***Figure S20***). Together, these results suggest that ReLERNN will be well suited to the increasing amount of population genomic data from non-model organisms. While the abstract nature of the data represented in its internal layers constrains our ability to interpret the exact information ReLERNN relies on to inform its predictions, our experiments using incorrect assumed mutation rates (***Figure S9, Figure S8***) suggests ReLERNN is potentially learning the relative ratio of recombination rates to mutation rates. Because the assumed rate of mutation governs the inherent potential for ReLERNN to resolve recombination events—i.e. recombination events cannot be detected without informative SNPs—and because simulation results suggest ReLERNN is more accurate when overestimating 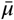 relative to underestimating it, we suggest erring on the side of overestimating 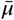. For these reasons however, an extra caveat is warranted—use caution when interpreting the results from ReLERNN as absolute measures of the per-base recombination rate unless precise mutation rate estimates are also known. This actually presents an opportunity—we suspect that ReLERNN (or a related network) has the potential to infer the joint landscape of recombination and mutation, though this task likely poses an additional set of unknown challenges.

Demographic model misspecification is another potential source of error that should affect not only deep learning methods targeted at estimating *ρ*, but also likelihood-based methods. Historical demographic events (e.g. population bottlenecks, rapid expansions, etc.), because they may alter the structure of LD genome-wide, can bias inference of recombination based on genetic variation data. Our simulations demonstrated that while all the methods we tested had elevated error in the context of demographic model misspecification, ReLERNN remained the most accurate across all misspecification scenarios (***Figure 4***). While we caution against generalizing too much from this experiment, the model misspecification tested here was extreme: we are replacing a human-like demography of a bottleneck followed by exponential growth with a model of constant population size. We suspect that ReLERNN, by using an RNN, is able to encode higher-order allelic associations across the genome, for instance three-locus or four-locus linkage disequilibrium, and in so doing capture more of the information available than traditional methods that use composite likelihoods of two-locus LD summaries. Additionally, there are clear opportunities for future improvements to ReLERNN. For instance, our simulation studies demonstrated that the GRU used by ReLERNN is also sensitive to gene conversion events (***Figure S19***), thus the joint estimation of rates of recombination and gene conversion may be quite feasible. Ultimately, it remains far from clear what network architectures will be best suited for population genetic inference, though we remain optimistic that ANNs will prove useful for a variety of applications in the field.

A natural application of ReLERNN, due in part to its high accuracy with small sample sizes, was to characterize and compare the recombination landscapes for multiple populations of African *D. melanogaster*, for which few populations with large samples sizes are currently available. Previous estimates of genome-wide fine-scale recombination maps in flies have focused on characterizing recombination in experimental crosses (***Comeron et al., 2012***), or by running LDhat (or the related LDhelmet) on populations with relatively moderate sample sizes (i.e. ≥ 22 samples) (***Chan et al., 2012; Langley et al., 2012***). Here, we applied ReLERNN to three populations for which at least ten haploid embryos were sequenced: Cameroon, Rwanda, and Zambia (***Lack et al., 2015; Pool et al., 2012***). Generally, recombination landscapes were well correlated among populations. Mean pairwise coefficients of determination among all three populations were *R*^2^ = 0.69, 0.61, 0.77, 0.43, 0.66 for chromosomes 2L, 2R, 3L, 3R, and X, respectively. These correlations are notably lower than those observed in humans (***Myers et al., 2005***) and mice (***Wang et al., 2017***), and one potential biological cause for this large difference could be the cosmopolitan chromosomal inversions that segregate in African *D. melanogaster* (***Corbett-Detig and Hartl, 2012; Lack et al., 2015***).

Our results suggest that recombination suppression extends well beyond the predicted breakpoints of the inversion (at least 5 Mb beyond in the case of *In(2L)t*; ***Figure S27, Figure S28***). This large-scale suppression of recombination due to inversions in *Drosophila* has been observed both directly in experimental crosses (***Dobzhansky and Epling, 1948; Novitski and Braver, 1954; Kulathinal et al., 2009; Miller et al., 2016; Fuller et al., 2018***), and indirectly from patterns of variation surrounding known inversion breakpoints (***Corbett-Detig and Hartl, 2012; Langley et al., 2012***). While it is true that the negative relationship between inversion frequency and recombination should only exist for inversions segregating at low frequencies (e.g. crossover suppression is not expected in inversion homozygotes), we predict a negative relationship to dominate in these populations, as the majority of polymorphic inversions are young, segregate at low frequencies, and show elevated LD along their lengths perhaps due to the actions of natural selection (***Corbett-Detig and Hartl, 2012; Lack et al., 2015***).

While polymorphic inversions exert strong effects on recombination landscapes, support for their role in explaining the most diverged regions among populations was mixed—we found that population-specific recombination rate outliers, but not global outliers, were significantly enriched within the inversions known to segregate in these populations (***Figure 6***). Moreover, our predictions for the relative rates of recombination among populations, based on inversion frequencies per chromosome, were largely not met—the inversions *In(2L)t, In(2R)NS*, and *In(3L)Ok* segregate at the highest frequencies in Zambia, yet this population also has the highest average recombination rate for these three chromosomes. One might speculate that such a result could be due to the reapportioning of crossovers that occurs due to the interchromosomal effect (***Schultz and Redfield, 1951***), although we have no firm evidence for this. Chromosome 3R, however, did match these predictions, having inversions segregating at the highest frequencies of any chromosome (e.g. *p*_*In(3R)K*_ = 0.9 in Cameroon) and also both the lowest coefficient of determination (*R*^2^ = 0.43) and population-specific recombination rates ranked in accordance with inversion frequencies (***Figure 6***).

Interestingly, while we identified two individual outlier regions characterized by numerous selective sweeps, we did not observe a significant enrichment of sweeps overlapping either global or population-specific outliers when these outliers were treated as a class of genomic elements. This is perhaps surprising, given that selective sweeps are known to create characteristic elevations of LD (***Kim and Nielsen, 2004***), and perhaps could mimic regions with very divergent levels of recombination in a population-specific way. A number of other evolutionary forces might explain the existence of our outlier regions as well. For example, mutation rate heterogeneity along the chromosomes could, in principle, generate spurious peaks or troughs in our estimates of recombination rate, as ReLERNN in effect scales its per-base recombination rate estimates by a mutation rate that is assumed to be constant along the chromosome (***Figure S9, Figure S8***). Moreover, introgression from diverged populations might affect patterns of allelic association in a a local way along the genome (***Schrider et al., 2018; Schumer et al., 2018***). Taken together, our results suggest that while both inversions and selection can influence population-specific differences in the landscape of recombination, the preponderance of these differences likely have complex causes.

While ReLERNN currently stands as a functional end-to-end pipeline for measuring recombination rates, the modular design herein presents a number of important opportunities for extension, with the potential to address myriad questions in population genomics. For example, the RNN structure we exploit here could be used for inferring the joint distribution of gene conversion and crossover events, or for inferring the distribution of selection coefficients and/or migration rates from natural populations. In addition, ReLERNN presents an excellent opportunity for the implementation of transfer learning, whereby ReLERNN could be trained in-house on an otherwise prohibitively extensive parameter space, allowing end-users to make accurate predictions by generating only a small fraction of the current number of simulations and training epochs presently required. The application of machine learning, and deep learning in particular, to questions in population genomics is ripe with opportunity. The software tools that we provide with ReLERNN support a simple foundation on which the population genetics community might begin this exploration.

## Materials and Methods

### The ReLERNN workflow

The ReLERNN workflow proceeds by the use of four python modules—`ReLERNN_SIMULATE`, `ReLERNN_TRAIN`, `ReLERNN_PREDICT`, and `ReLERNN_BSCORRECT` (or alternatively `ReLERNN_SIMULATE_POOL`, `ReLERNN_TRAIN_POOL`, and `ReLERNN_PREDICT_POOL` when analysing Pool-seq data). The first three modules are mandatory, and include functions for estimating parameters such as *θ*_*W*_ and *N*_*e*_ from the inputs, functions for masking genotypes and inaccessible regions of the genome, functions for simulating the training, validation, and test set, functions for training the neural network, and functions for predicting rates of recombination along the chromosomes. The fourth module, `ReLERNN_BSCORRECT`, can be used both with individually sequenced data and Pool-seq data. This module is optional (though recommended) and includes functions for estimating 95% confidence intervals and implementing a correction function to reduce bias. The output from ReLERNN is a list of genomic windows and their corresponding recombination rate predictions (reported as per-base crossover events), along with 95% confidence intervals and corrected predictions through the use of `ReLERNN_BSCORRECT`.

#### Estimation of simulation parameters and coalescent simulations

ReLERNN takes as input a VCF file of phased or unphased biallelic variants. A minimum of four sample chromosomes must be included, and users should ensure proper quality control of the input file beforehand—e.g. filtering low-coverage, low-quality, and non-biallelic sites. ReLERNN for Pool-seq takes a single file of genomic coordinates and their corresponding pooled allele frequency estimates (example files can be found at https://github.com/kern-lab/ReLERNN/tree/master/examples). ReLERNN then steps along the chromosome in non-overlapping windows of length *l*, where *l* is the minimum window size for which the number of segregating sites, *S*, in all windows is ≤ 1750. By default, we require that *S* ≤ 1750, as extensive experimentation during development showed that *S* >> 1750 has the potential to cause the so-called exploding gradient problem to arise during training (see ***Pascanu et al., 2013***). However, *S* is a user-configurable parameter (− − *maxW inSize*), and can be increased at the expense of potential training failures. The minimum number of sites in a window is another user-configurable parameter (− − *minSites* in `ReLERNN_PREDICT`) and is set to 50 by default. As a result of independently estimating *l* for each chromosome, the output predictions file may return different window sizes for different chromosomes, depending on SNP densities.

Once *l* has been estimated, `ReLERNN_SIMULATE` uses the coalescent simulation software, msprime (***Kelleher et al., 2016***), to independently generate 10^5^ training examples and 10^3^ validation and test examples. By default, these simulations are generated under assumptions of demographic equilibrium using the following parameters in msprime: [*sample*_*size* = *n*, where *n* is the number of chromosomes in the VCF; *Ne* = *N*_*e*_, where 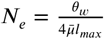 and 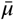 is the assumed genome-wide perbase mutation rate, *l*_*max*_ is the maximum value for *l* across all chromosomes, and 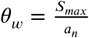 where *S*_*max*_ is the genome-wide maximum number of segregating sites for all windows and 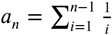; *mutation*_*rate* = *U*(*μ*_*low*_, *μ*_*high*_), where 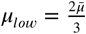 and 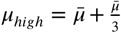; *recombination*_*rate* = *U*(0.0, *r*_*max*_), where 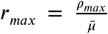, and *length* = *l*_*max*_]. In addition to simulating under equilibrium, ReLERNN can also simulate under a population size history inferred by one of three programs: stairwayplot (***Liu and Fu, 2015***), SMC++ (***Terhorst et al., 2016***), or MSMC (***Schiffels and Durbin, 2014***). This is handled by proving the raw final output file to `ReLERNN_SIMULATE` using the −− *demographicHistory* option. When a demographic history is supplied to ReLERNN, the *Ne* parameter in msprime is substituted with a history of population size changes through time, but the *mutation*_*rate, recombination*_*rate*, and *length* parameters are the same as when simulating under equilibrium. After each simulation is completed, ReLERNN writes both the genotype matrix and a vector of SNP coordinates to temporary.npy files, which are later used during batch generation.

#### Sequence batch generation and network architectures

To reduce the large memory utilization common to the analysis of genomic sequence data, we took a batch generation approach using the `fit_generator` function in `Keras`—i.e. only small batches (defaulty *batch*_*size* = 64) of simulation examples are called into memory at any one time. Moreover, both the order of examples within each batch, and the order of individuals within a single training example are randomly shuffled (i.e. sample 1 is not always at the top of the genotype matrix). Data normalization and padding occurs when a training batch is called, and the genotype and position arrays are read into memory. The zeroth axis of the genotype and positions arrays is then padded with 0s (*pad*_*size* = 5) to *max*(*S*_*max*_, *S*_*sim*_), where *S*_*max*_ is the genome-wide maximum number of segregating sites for all windows in the samples and *S*_*sim*_ is the maximum number of segregating sites generated across all training, validation, and test simulations.

The targets for each training batch are the per-base recombination rates used by msprime to simulate each example. These targets are z-score normalized across all training examples. Genotypes and positions are not normalized, per se. Rather, the genotype matrix encodes alleles as reference (−1), alternative (1), or padded/missing data (0), and variant positions are encoded along the real number line (0−1). In the case of ReLERNN for Pool-seq, we convert the simulated genotypes into allele frequencies by sampling with replacement the vector of alternative and reference alleles for all sites to the assumed mean read depth of the pool (a user supplied parameter). We then exclude any site where the sampled variant is fixed or where *p*< 0.05, and stack this newly created allele frequency vector with the vector of positions. Here, allele frequencies (but not positions) are z-score normalized. The normalized and padded genotype, position, and allele frequency arrays form the input tensors to our neural networks, and take the shapes defined in ***Figure S1***.

ReLERNN trains a recurrent neural network with `Keras` (***Chollet et al., 2015***) using a `Tensorflow` backend (***Abadi et al., 2015***). The complete details of our neural architecture can be found in the python module https://github.com/kern-lab/ReLERNN/blob/master/ReLERNN/networks.py, and a detailed flow diagram showing the connectivity between layers as well as network parameters can be found in ***Figure S1***. Briefly, the ReLERNN neural network utilizes distinct input layers for the genotype and position tensors, which are later merged using a concatenation layer in Keras. The genotype tensor is first fed to a GRU layer, as implemented with the bidirectional wrapper in Keras, and the output of this layer is passed to a dense layer followed by a dropout layer. On the positions side of the network, the input positions tensor is fed directly to a dense layer and then to a dropout layer. Dropout (***Srivastava et al., 2014***) was used extensively in our network, and accuracy was significantly improved when employing dropout relative to networks without dropout. Once concatenated, output from the dropout layer is passed to a final round of dense and dropout layers, and the final dense layer returns a single z-score normalized prediction for each example, which is unnormalized back to units of crossovers per-base. ReLERNN implements early stopping to terminate training (*min*_*delta* = 0.01, *patience* = 100) and uses the “Adam” optimizer (***Kingma and Ba, 2014***) and a Mean Squared Error (*MSE*) loss function. Our hyper-tuning trials were completed via a grid search over the set of parameters: recurrent layer output dimensions (64, 82, 128), loss function (*MSE, MAE*), input merge strategy (concatenate, average), and dense layer output dimensions (64, 128), optimizing for *MSE*.

Total runtime estimates are highly dependent on 1) the number of epochs needed to train before the early stopping threshold is met (which can vary extensively) and 2) the coalescent simulation parameters (most notably recombination rate and population size). As an example, the total runtime for `ReLERNN_SIMULATE`, `ReLERNN_TRAIN`, and `ReLERNN_PREDICT` on a 1 Mb chromosome with 90290 segregating sites [parameters: *n* = 20, 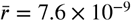, and 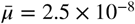], which trained for 348 epochs before terminating, was 8527 seconds (40 cores Intel Xeon, 1 NVIDIA 2070 GPU). Total runtimes are not strongly influenced by genome size—e.g. the time needed for ReLERNN to make predictions on the 90290 SNPs in the example above was less than 8.2 seconds.

#### Parametric bootstrap analysis and prediction corrections

ReLERNN includes the option to generate confidence intervals around each predicted recombination rate and correct for potential biases generated during training. To accomplish this we used parametric bootstrapping, as implemented by `ReLERNN_BSCORRECT` in the following way: after the network has been trained and predictions have been generated, `ReLERNN_BSCORRECT` simulates 10^3^ test examples for each of 100 recombination rate bins drawn from the distribution of recombination rates used to train the network. The parameters for each new simulation example are drawn from the same distribution of parameters used to simulate the original training set, with the exception of *recombination*_*rate*, which is held constant for each rate bin. Predictions are then generated for these 10^5^ simulated test examples using the previously trained network, generating a distribution of predictions for each respective recombination rate bin. 95% confidence intervals are calculated for each bin by taking the upper and lower 2.5% predictions from this distribution of rates.

The distribution of test predictions can potentially be biased in systematic ways—e.g. predictably underestimating rates of recombination for those examples with the highest simulated crossover events, possibly due to the limited ability to resolve high recombination rates with a finite number of SNPs. From our inferred confidence intervals we can correct for inferred bias in the following way. The bias correction function takes each empirical prediction, *r*_*predicted*_, and identifies the nearest median value, 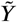, from the distribution of 10^5^ bootstrap rate predictions (***Figure S3***). Because each 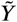 was generated from a rate bin corresponding to the true recombination rate, *Y*, we can apply the correction function, 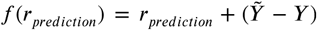, to all predictions. This method has the effect of increasing *r*_*predicted*_ in areas of parameter space where we are reasonably confident that we are underestimating rates and reducing *r*_*predicted*_ in areas where we are likely to be overestimating rates. `ReLERNN_BSCORRECT` is provided as an optional module for this task, as the resimulation of 10^5^ test examples has the potential to be computationally expensive, and may not be warranted in all circumstances. However, as stated above, the extent of the computational expense is highly dependent on the parameters used in the coalescent simulation, and may not always contribute substantially to total runtimes. For example, `ReLERNN_BSCORRECT` increased the total runtime in the example mentioned above by 8.6 percent (9266 seconds compared to 8527 seconds).

### Testing the accuracy of ReLERNN on simulated recombination landscapes

To test the accuracy of ReLERNN at recapitulating a dynamic recombination landscape, we ran our complete ReLERNN workflow on simulation data replicating chromosome 2L of *D. melanogaster*. Using crossover rates estimated by ***Comeron et al***. (***2012***), we simulated varying numbers of samples of *D. melanogaster* chromosome 2L with msprime using the RecombinationMap class [parameters: *n* ∈ {4, 20, 50}, 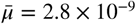, *N*_*e*_ = 2.5× 10^5^]. Simulated samples were exported to a VCF file using ploidy = 1, and all simulations were generated under demographic equilibrium. We used these simulated VCF files as the input to our ReLERNN pipeline, where we varied the assumed 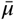 and the assumed ratio of *ρ*_*max*_ to *θ* given to ReLERNN. The assumed 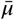 was varied from 50% less than the rate used in simulations (2.8 × 10^−9^) to 50% greater than the true rate. Likewise, the ratio of *ρ*_*max*_ to *θ* was either held constant, resulting in the training set containing on average higher or lower per-base recombination rates than the true rate, or was adjusted to correctly reflect the true maximum per-base recombination rate used—i.e. approximately 1.2× 10^−7^ crossovers per base. To run ReLERNN on simulated Pool-seq data we used the same VCFs generated above, but converted all variants to allele frequencies in the following way: for all sites in the VCF, we resampled the variant haplotypes with replacement to a simulated read depth of 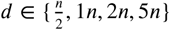 and then excluded all sites where the resampled variant was fixed or where *p*< 0.05.

### Comparative methods

We chose to compare ReLERNN to three published methods for estimating recombination rates— FastEPRR (***Gao et al., 2016***), a 1-dimensional CNN recently described in ***Flagel et al***. (***2018***) and both LDhat (***McVean et al., 2002***) and LDhelmet (***Chan et al., 2012***). We generated a training set (used by ReLERNN and the CNN) with 10^5^ examples and tested all of the methods on an identical set of 5× 10^3^ simulation examples. We generated two classes of simulations, one simulated under demographic equilibrium and one using a demographic history derived from European humans (CEU model; detailed in “ReLERNN_demographic_models.py”; ***Tennessen et al., 2012; Gravel et al., 2011***). Both classes of simulations were generated for *n* ∈ {4, 8, 16, 32, 64}, where *n* is the number of chromosomes sampled from the population. All simulations were generated in msprime with the common set of parameters [*recombination*_*rate* = *U*(0.0, 6.25*e*^−8^), *mutation*_*rate* = *U*(1.875*e*^−8^, 3.125*e*^−8^), *length* = 3*e*^5^].

For both ReLERNN and the CNN, the same training set consisting of 10^5^ examples was used to train each neural network, and the same test examples were used to compare the predictions produced by each method. Comparisons with LDhat and LDhelmet where made using the above training examples to parameterize the generation of independent coalescent likelihood lookup tables. For each set of examples of sample size *n*, we used the known value of *ρ*_*max*_ from the simulated training examples, and we then calculated the average per-base values for *θ* from the simulated test examples using Watterson’s estimator. These parameter values were passed to the functions for lookup table generation in LDhat and LDhelmet [LDhat options: −*n*, −*rhomax*, −*theta* and −*n*_*pts*101; LDhelmet options: −*r*0.00.110.01.0100.0]. For LDhelmet we also ran the *pade* function using the options [−*x*12 and − −*def ect*_*threshold*40]. The resulting tables were used to make predictions on our 5× 10^3^ test examples using the *pairwise* function for LDhat and *max*_*lk* function for LDhelmet [options: − − *max*_*lk*_*start*0.0 and − − *max*_*lk*_*resolution*0.000001]. Comparisons with FastEPRR were made by transforming the genotype matrices resulting from our test simulations into fasta-formated input files, and running the FastEPRR_ALN function [using format = 1] in R. As LDhat, LDhelmet, and FastEPRR all predict *ρ*, the resulting predictions were transformed to per-base recombination rates for direct comparison with ReLERNN using the function 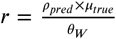, where *ρ*_*pred*_ is the prediction output by each method, and *θ*_*W*_ and *μ*_*true*_ are Watterson’s estimator and the true per-base mutation rate used in the simulation example, respectively. To compare accuracy among methods we directly compared the distribution of absolute errors (|*r* _*predicted*_ − *r*_*true*_|) for each method for each set of examples of sample size *n*.

To test the effects of model misspecification on predictions, we simply directed ReLERNN and the CNN to use a training set generated under demographic equilibrium for making predictions on a test set generated under the CEU model, and vice versa. To test for the effects of model misspecification in LDhat and LDhelmet, we generated a lookup table using parameter values estimated from the misspecified training set (e.g. the lookup table used for predicting the CEU model test set was generated by using parameter values directly inferred from training simulations under equilibrium. We did not directly test the effect of model misspecification using FastEPRR, as this method takes as input only a fasta sequence file, and therefore the internal training of the model was not able to be separated from the input sequences. To address the effects of model misspecification, we also directly compared the distribution of absolute errors (|*r*_*predicted*_ − *r*_*true*_ |).Additionally, we compared the marginal error directly attributable to model misspecification among methods. We defined marginal error as *ϵ*_*m*_ − *ϵ*_*c*_, where *ϵ*_*m*_ and *ϵ*_*c*_ are equal to |*r*_*predicted*_ − *r*_*true*_| when the model is misspecified and correctly specified, respectively. We simulated gene conversion test sets using ms (***Hudson, 2002***), with a mean conversion tract length of 352 bp (corresponding to the mean empirically derived tract length in *D. melanogaster* (***Hilliker et al., 1994***)) and simulated a range of gene conversion to crossover ratios, 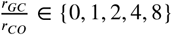.

### Training on missing genotypes and inaccessible regions of the genome

Deep neural networks, through their aptitude for pattern recognition, can be trained to infer information from missing data. To harness this ability, we took two different approaches: 1) we infer patterns of recombination when some fraction of individual genotype calls are absent (missing genotypes), and 2) we infer these patterns when some fraction of all sites cannot be sequenced (genome inaccessibility). To simulate levels of missing genotypes similar to those found in real data, we first sample the distribution of all missing genotypes from the input VCF. We then generate a missing genotype mask for all windows in the genome and write this mask as a temporary file to the disk. Simulation proceeds as if all genotypes are present, however during batch generation, one random mask is drawn from the genomic distribution of masks and applied to the generated genotype matrix, setting some fraction of genotype calls to 0 (the same element used to pad). This has the effect of training the network to infer recombination, even where genotype calls are missing in real data. To infer recombination in the face of genome inaccessibility, we take a similar approach. Here, ReLERNN accepts an empirical accessibility mask similar to that provided by the 1000 Genomes project (***Consortium et al., 2015***). This is provided in BED format, which is then fragmented into smaller arrays corresponding to the window size used by `ReLERNN_SIMULATE`. After simulation proceeds with all sites present, we randomly draw a mask from the distribution of empirical accessibility masks, and apply it during batch generation, removing all sites marked inaccessible from the array. We then remove the corresponding sites from the positions array, and train as usual.

To test ReLERNN’s ability to learn recombination rates in the face of missing genotypes and genome inaccessibility, we simulated a 1 Mb randomize dynamic recombination landscape in msprime. Here we randomly selected 39 sites along the chromosome to serve as recombination rate breakpoints, generating 40 windows of different rates. For each rate multiplier, *m* ∈ {3, 3, 3, 3, 3, 5, 5, 5, 5, 5, 7, 7, 7, 10, 10, 10}, we randomly selected a window to have the recombination rate 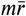, were 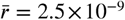 is the simulated background recombination rate. To simulate missing genotypes, we randomly set genotype calls in the simulated VCF to a., corresponding to a fraction of total genotypes ∈ {0.0, 0.10, 0.25, 0.50}. To simulate an empirical accessibility mask we simply sampled directly from the phase 3 1000 Genomes accessibility masks (***Consortium et al., 2015***) and removed sites in the VCF corresponding to a fraction of total genomic sites ∈ {0.0, 0.25, 0.50, 0.75}. To directly compare between the predictions made by ReLERNN and LDhelmet, we then broke the VCF into windows of the same length (e.g. 22 kb for *n* =4 and 10 kb for *n* = 20 for the simulations with missing genotypes). We then ran both ReLERNN and LDhelmet as described above, and compared the distribution of absolute errors (|*r*_*predicted*_ − *r*_*true*_|) for each method for each set of examples of sample size *n* ∈ {4, 20}.

### Recombination rate variation in *D. melanogaster*

We obtained *D. melanogaster* population sequence data from the *Drosphila* Genome Nexus (DGN; https://www.johnpool.net/genomes.html; ***Lack et al., 2015; Pool et al., 2012***). We converted DGN “consensus sequence files” to a simulated VCF format, excluding all non-biallelic sites and those containing missing data. We chose to analyze populations from Cameroon, Rwanda, and Zambia, as these populations contained at least 10 haploid embryo sequences per population and each population included multiple segregating chromosomal inversions (supplemental table 1). To ensure roughly equivalent power to compare rates among populations, we downsampled both Rwanda and Zambia to 10 chromosomes. We selected individual haploid genomes for each population by requiring that our sampled inversion frequencies for each of the six segregating inversions—*In(1)Be, In(2L)t, In(2R)NS, In(3L)Ok, In(3R)K*, and *In(3R)P*—closely approximate their population frequencies as measured in the complete set of haploid genomes for that population. All sample accessions and their corresponding inversion frequencies are located in the supporting materials.

Before running ReLERNN, we first set out to model the demographic history for each population using each of three methods: stairwayplot (***Liu and Fu, 2015***), SMC++ (***Terhorst et al., 2016***), and MSMC (***Schiffels and Durbin, 2014***). With the exception of MSMC, all methods were run using default parameters. For MSMC, the use of default parameters generated predictions that were unusable (***Figure S22***). For these reasons, and after direct communication with MSMC’s authors, we determined that running MSMC with a sample size of two chromosomes would be the most appropriate. Using all three methods, we show that inferred historical population sizes are unreliable for these populations—no two methods recapitulate the same history, and the histories generated by MSMC vary dramatically depending on the number of samples used (***Figure S21, Figure S22***). For these reasons, and because results from our simulations suggest that marginal error due to demographic misspecification is quite low for our method (***Figure S18***), we decided to simulate our training data under the assumptions of demographic equilibrium [options: − − *estimateDemographyFalse* − − *assumedMu*3.27*e* −9 − − *upperRhoT hetaRatio*35].

We measured the correlation in recombination rates between each African *D. melanogaster* populations by recalculating the raw rate for 100 kb sliding windows, as ReLERNN will predict the rates of recombination in slightly different window sizes, depending on *θ* for each chromosome. The recombination rate for each 100 kb window was calculated by taking the average of all raw rate windows predicted by ReLERNN, weighted by the fraction that each window overlapped the larger 100 kb sliding window. Recombination rate outliers were identified in two ways: as global outliers and population-specific outliers. Global outliers were identified by first calculating the mean and standard deviation in recombination rates for all three populations in each 100 kb sliding window. We then used the top 1% of outliers from the distribution of residuals, after fitting a linear model to the standard deviation on the mean. Population-specific outliers were identified by using a modification of the population branch statistic (herein PBS*; ***Yi et al., 2010***), whereby we replaced pairwise *F*_*ST*_ with the pairwise differences in recombination rates. We then used the top 1% of all PBS* scores as our population-specific outliers, with each outlier corresponding to a PBS* score for a single population.

To test the effect of inversion frequency on predicted recombination rates, we resampled 10 haploid chromosomes from the available set of haploid genomes from Zambia to generate sampled populations containing *In(2L)t* at varying frequencies, *p* ∈ {0.0, 0.2, 0.6, 1.0}. We then ran ReLERNN on chromosome 2L for each of these resampled Zambian populations. We classified recombination windows by their overlap with the coordinates of In(2L)t (as defined in ***Corbett-Detig and Hartl, 2012***), defining windows within the breakpoints (inside), windows up to 3 Mb outside the breakpoints (flanking), and windows > 3 Mb outside the breakpoints (outside). Recombination rates were negatively correlated with inversion frequency in our sample, not only within the inversion, but also in regions 3 Mb outside the inversion (flanking regions) (*ρ*_*Spearman*′*s*_ = −1; *P* = 0.04 for both comparisons). We also saw a similar negative correlation outside the flanking regions, although this association was weakened relative to that within or flanking the inversion (***Figure S27***). Importantly, varying the size of the flanking regions (from 1-5 Mb) produced patterns that were qualitatively identical, suggesting that the effect of inversions on recombination suppression extends far beyond the inversion breakpoints themselves (***Figure S28***).

We also expect that rates of recombination should be correlated with distance to the inversion breakpoint on smaller spatial scales. Likewise, recombination rates in the inversion interior (> 2 Mb from the breakpoints) are expected to be higher than in those regions immediately surrounding the breakpoints. To test this we looked at the recombination rates in our African *D. melanogaster* populations, binned by distance to the nearest inversion breakpoints segregating in these populations. We classified windows by their overlap with inversion interiors (> 2 Mb inside the inversion breakpoints) and their overlap with windows within 200 Kb, 500 Kb, 1 Mb, and 2 Mb of inversion breakpoints. We found that recombination rates in the flanking regions are positively correlated with distance to inversion breakpoints in both Rwanda and Zambia (*ρ*_*Spearman*′*s*_ = 1; *P* = 0.04 for both comparisons) but not in Cameroon (*ρ*_*Spearman*′*s*_ = 0.8; *P* = 0.17; ***Figure S25***). However, with the exception of Cameroon (Inversion interior compared to < 250 Kb from breakpoint; *P*_*WTT*_ = 0.035), we did not observe this pattern (*P*_*WTT*_ ≥ 0.057; ***Figure S25***).

We tested for an enrichment of both global and population-specific outliers within inversions by randomization tests, permuting the labels for outliers 10^4^ times and counting the overlap with inversions for each permutation to calculate the empirical p-values. We also tested for an effect of selection on recombination rates in these populations, by running diploS/HIC (***Kern and Schrider, 2018***) to detect selective sweeps. We ran diploS/HIC on each population, training on simulations generated under demographic equilibrium. For each population we simulated 2000 training examples from each of the five classes of regions required by diploS/HIC using the coalescent simulation software discoal (***Kern and Schrider, 2016***). For simulations which included sweeps we drew the selection coefficient from a uniform distribution such that *s* ∼ *U* (0.0001, 0.005), the time of completion of the sweep from *τ* ∼ *U* (0, 0.05), and the frequency at which a soft sweep first comes under selection as *f* ∼ *U* (0, 0.1). We drew *θ* from *U* (65, 654) and we drew *ρ* from an exponential distribution with mean 1799 and the upper bound truncated at triple the mean. For the discoal simulations we simulated 605 kb of data with the goal of classification of the central most 55 kb window. We looked at the overlap with “sweep” windows (those classified as either “hard” or “soft”) and those windows classified as “neutral” by diploS/HIC. Our complete diploS/HIC pipeline for these samples is available in the supporting materials online. All statistical tests were completed in R (***R Core Team, 2018***), with the exception of empirical randomization tests, which were completed using Python.

## Data availability

ReLERNN is currently available at https://github.com/kern-lab/ReLERNN. Supporting information, tables, and figures will be deposited online at the publication journal.

## Acknowledgments

The authors would like to gratefully acknowledge Matthew Hahn, Dan Schrider, and Peter Ralph for their helpful comments and suggestions. This work benefited from access to the University of Oregon high performance computer, Talapas. JRA, JGG, and ADK were supported by NIH award R01GM117241 to ADK.

---

**Figure S1.**
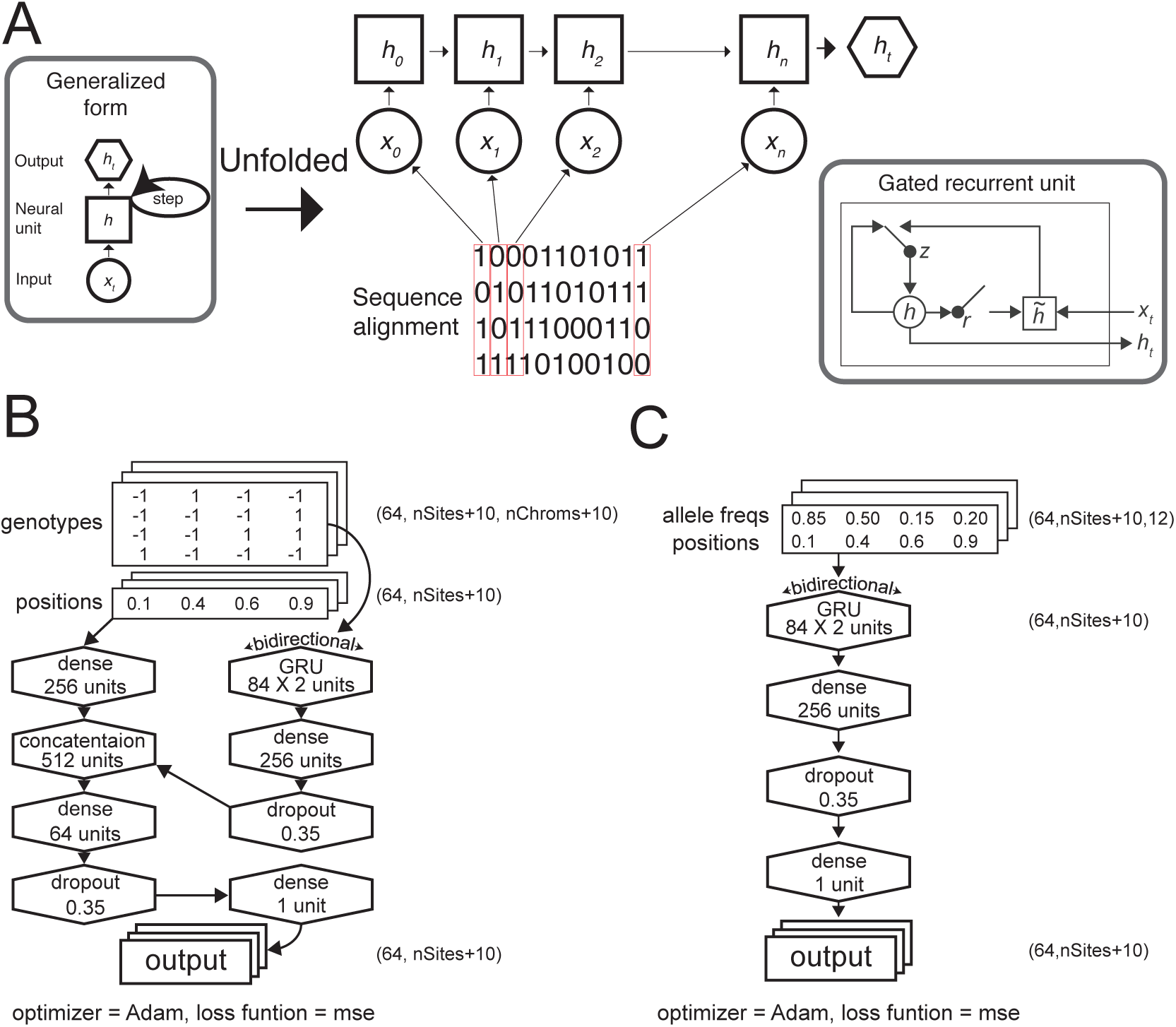
**(A, left)** Generalized form for a recurrent neural network trained on a genomic sequence alignment. **(A, right)** Generalized form of each gated recurrent unit, where *r, z, h*_*t*_, and 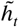 correspond to the reset gate, update gate, activation, and candidate activation, respectively (***Cho et al., 2014***). **(B)** Cartoon depicting the neural network architectures used in ReLERNN for individually sequenced genomes or **(C)** pooled sequences. Tensor shapes are shown for the default parameters [*batchsize* = 64, *padsize* = 5].

**Figure S2.**
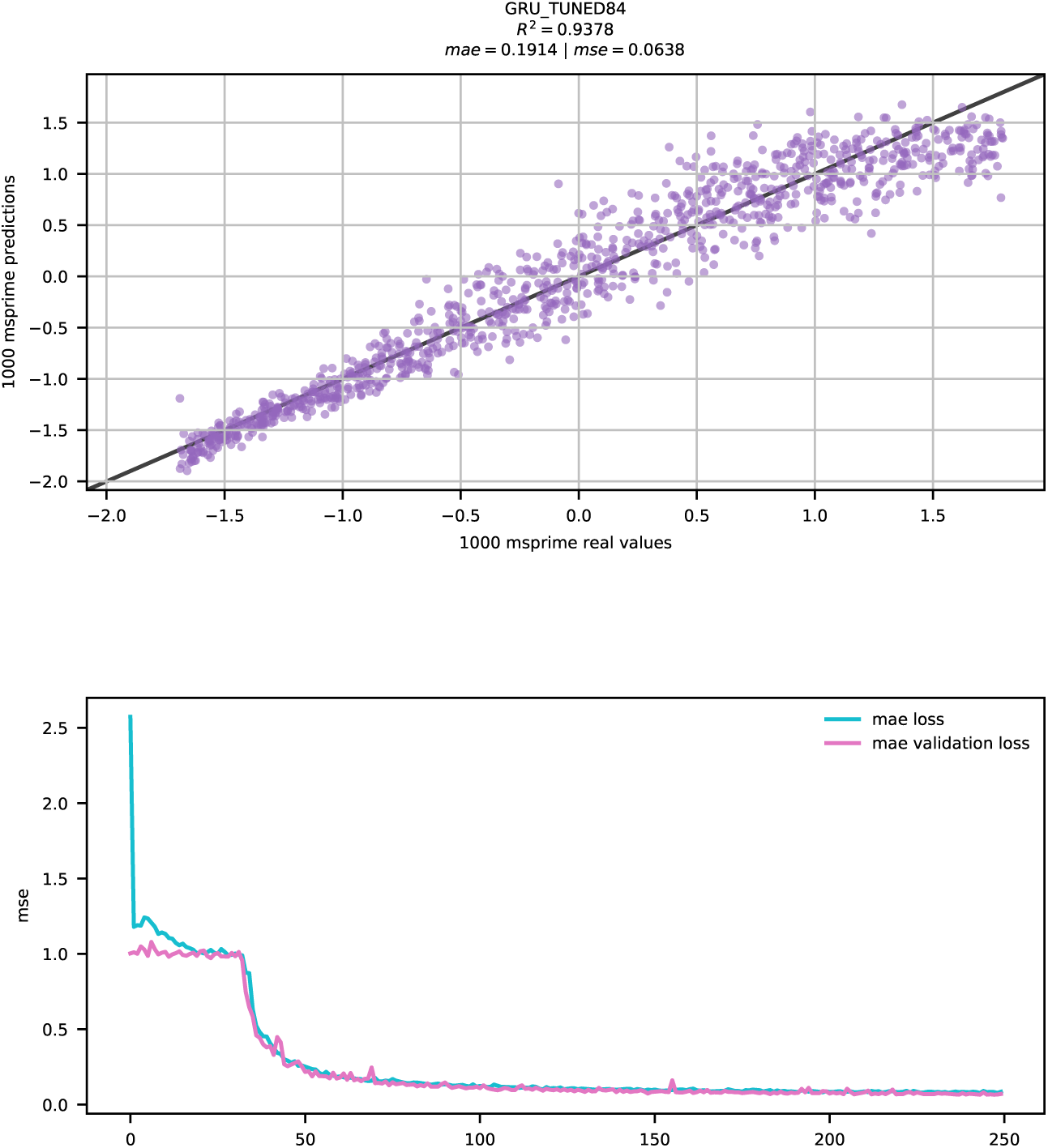
ReLERNN training and test results. **(Top)** Scatter plot of raw (unnormalized) predictions for 1000 test examples using ReLERNN with the same parameters used in ***Figure 2***. Mean absolute error and mean squared error are shown. **(Bottom)** Line graph showing the convergence of loss (measured by mean squared error) over time (epochs) during training on the same data as above, for both the training set (blue lines) and the validation set (purple lines).

**Figure S3.**
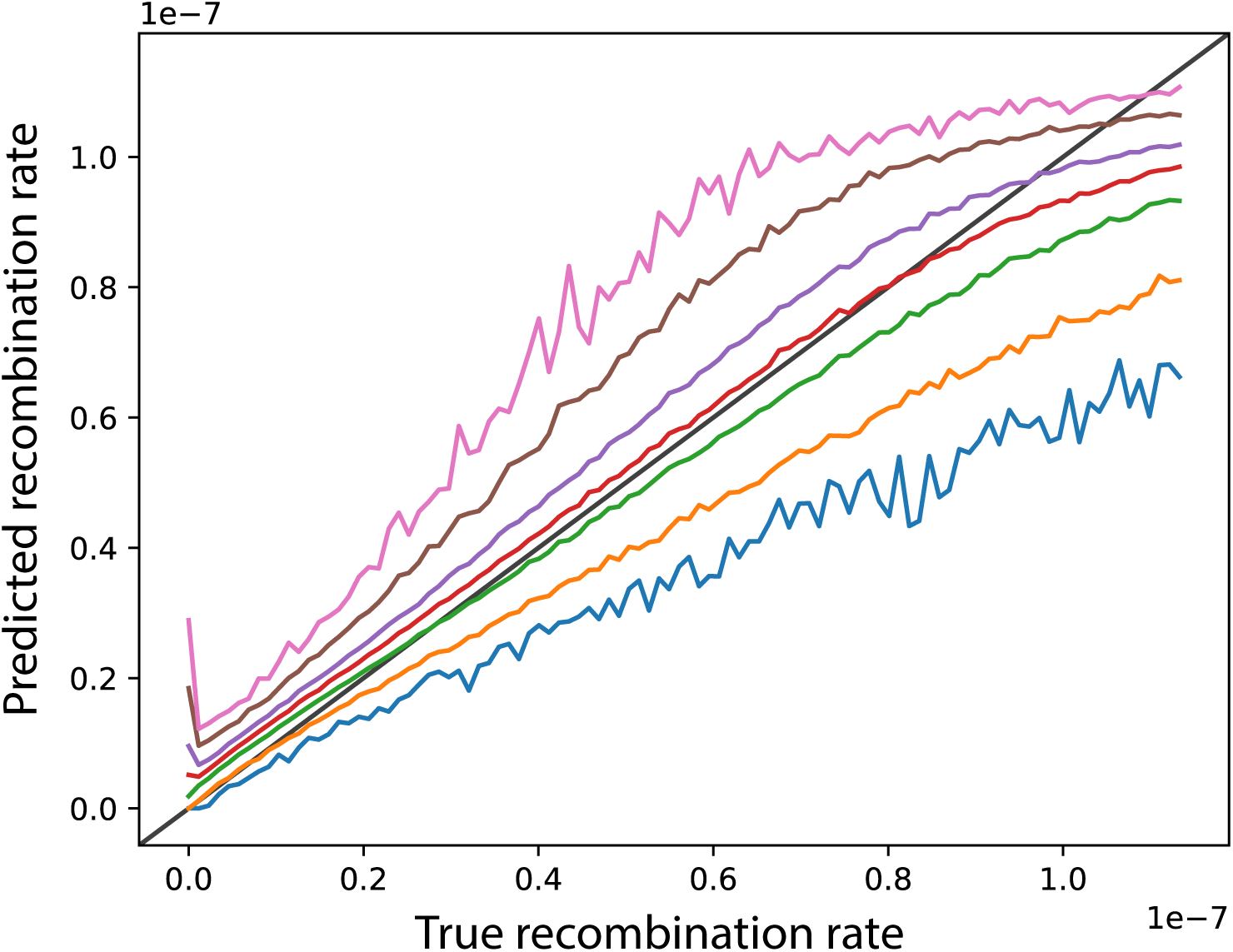
A characteristic example distribution of parametric bootstraping predictions, as implemented by `ReLERNN_BSCORRECT`. Lines represent the minimum (blue), lower 5% (orange), lower 25% (green), median (red), upper 25% (purple), upper 95% (brown), and maximum (pink) bounds for each of 1000 replicate simulations and predictions (y-axis) across 100 recombination rate bins (x-axis)

**Figure S4.**
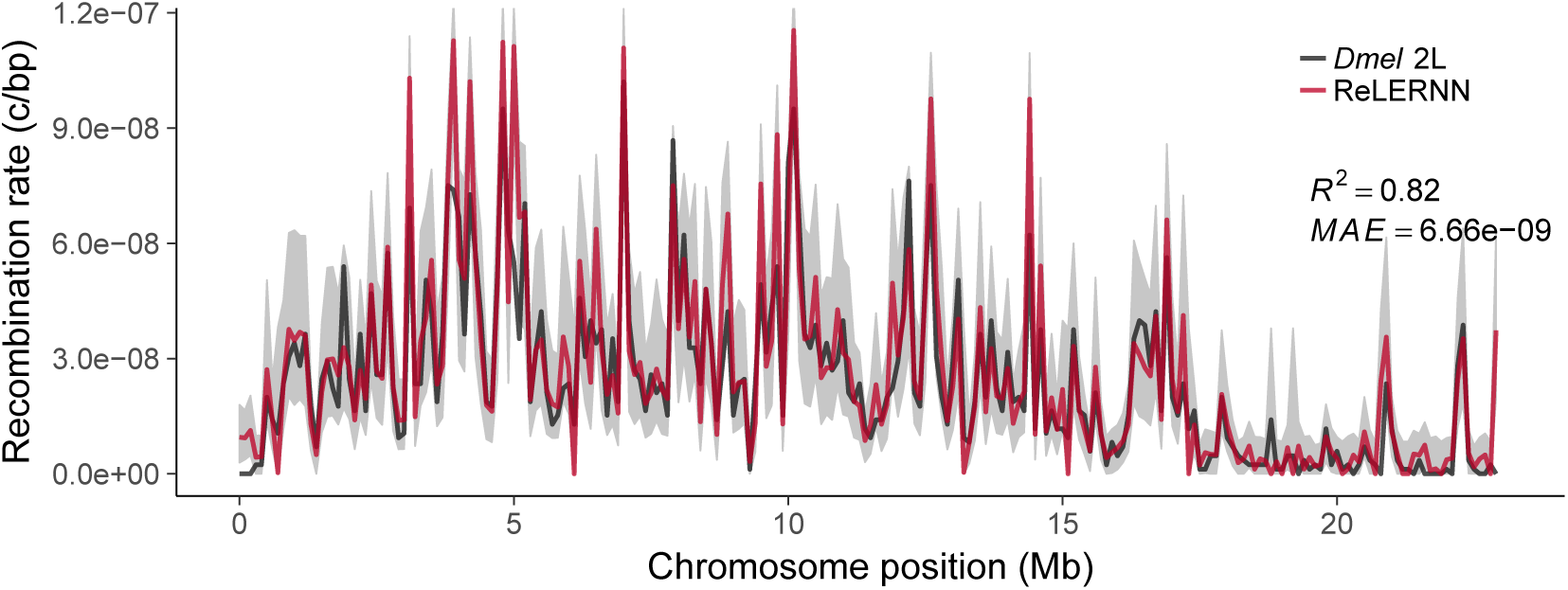
Recombination rate predictions for a simulated *Drosophila* chromosome (black line) using ReLERNN (red line). The recombination landscape was simulated for *n* = 4 chromosomes under mutation-drift equilibrium using msprime (***Kelleher et al., 2016***), with per-base crossover rates derived from *D. melanogaster* chromosome 2L (***Comeron et al., 2012***). Gray ribbons represent 95% confidence intervals. *R*^2^ is reported for the general linear model of predicted rates on true rates and mean absolute error was calculated across all 100 kb windows.

**Figure S5.**
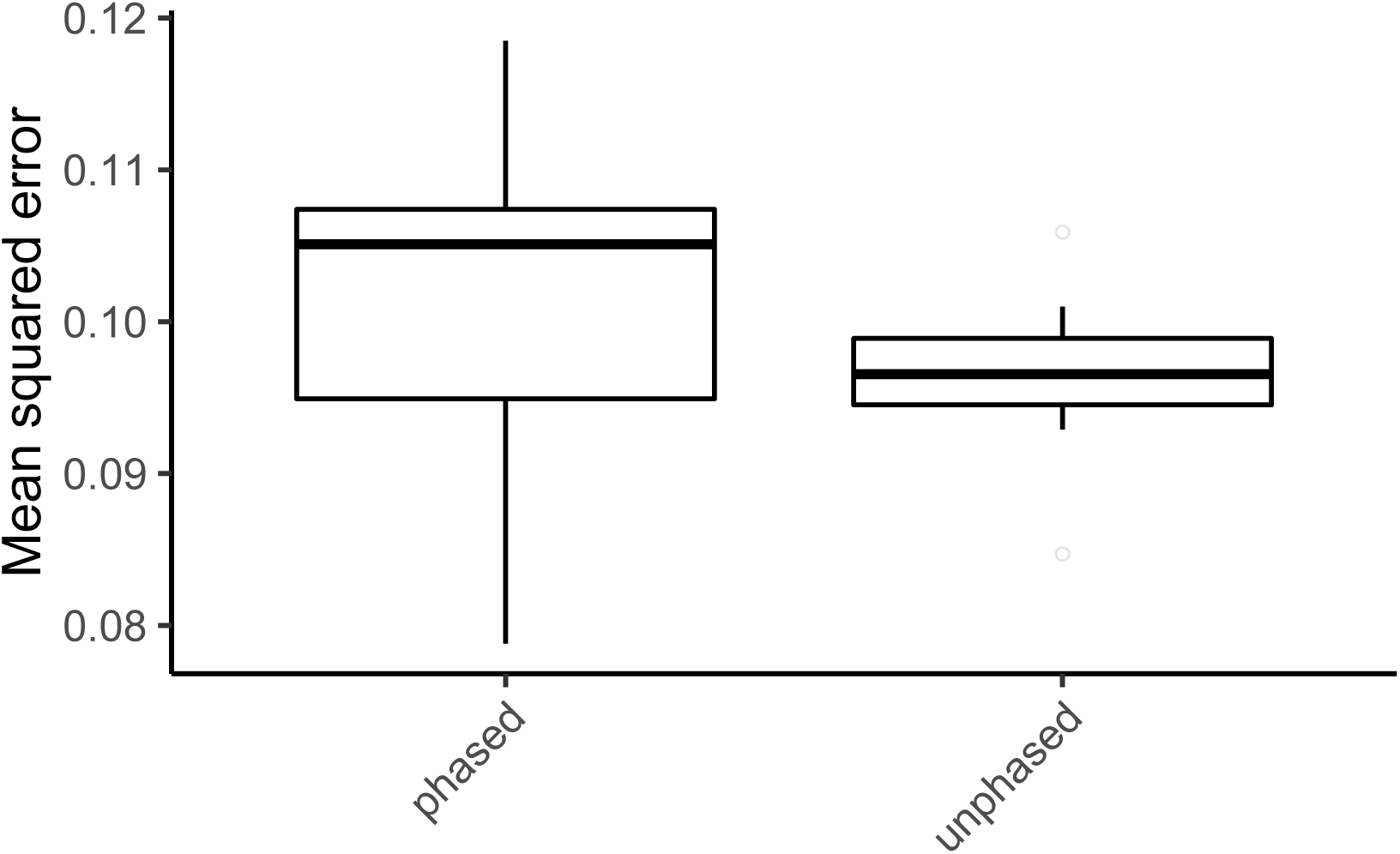
Mean squared error for ReLERNN predictions on 10 replicates of 1000 test simulations using 100% correctly phased input genotypes and completely unphased genotypes. All simulations used the recombination map derived from *D. melanogaster* chromosome 2L (***Comeron et al., 2012***).

**Figure S6.**
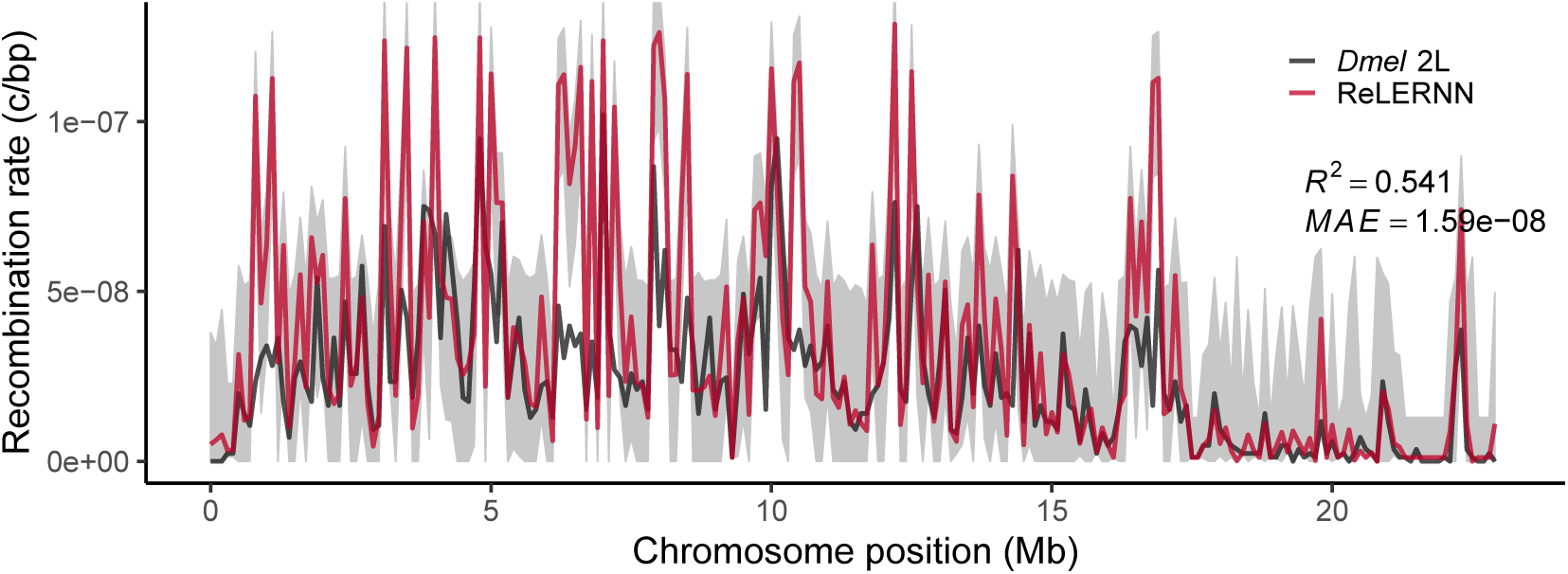
Recombination rate predictions from Pool-seq data for a simulated *Drosophila* chromosome (black line) using ReLERNN (red line). The recombination landscape was simulated for *n* = 50 chromosomes and a read depth of 50*X*, under mutation-drift equilibrium using msprime (***Kelleher et al., 2016***), with per-base crossover rates derived from *D. melanogaster* chromosome 2L (***Comeron et al., 2012***). Gray ribbons represent 95% confidence intervals. *R*^2^ is reported for the general linear model of predicted rates on true rates and mean absolute error was calculated across all 100 kb windows.

**Figure S7.**
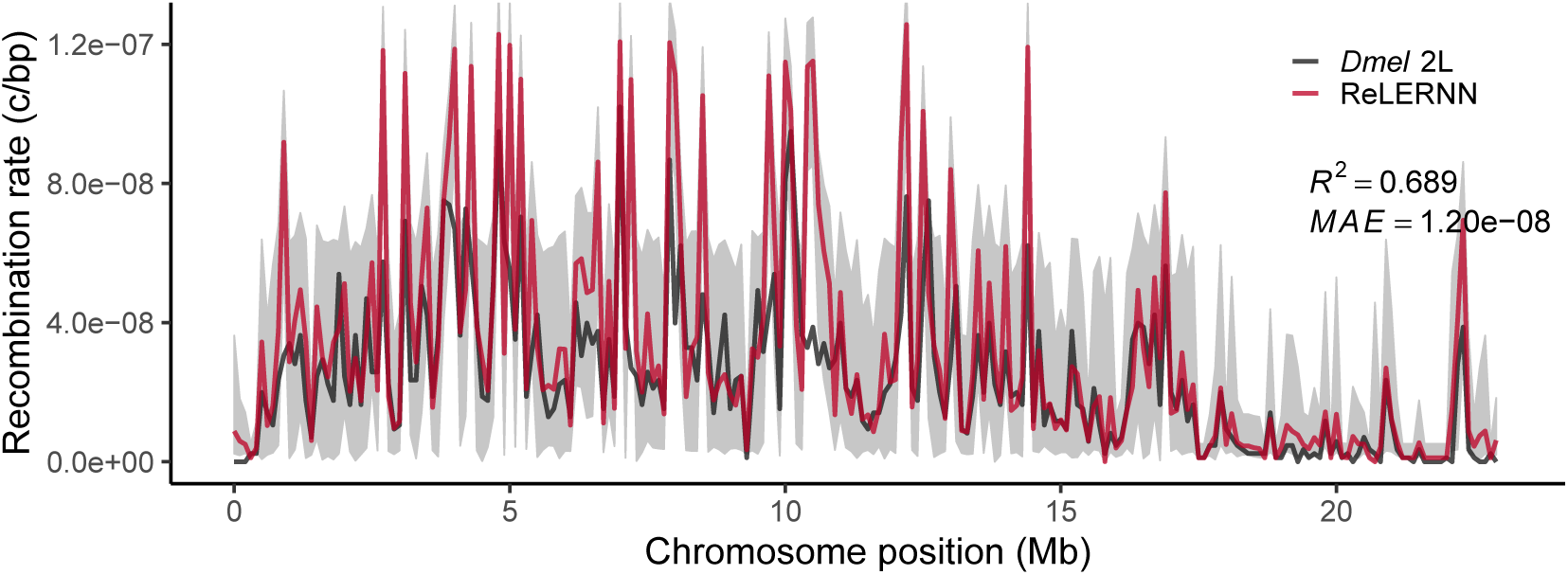
Recombination rate predictions from Pool-seq data for a simulated *Drosophila* chromosome (black line) using ReLERNN (red line). The recombination landscape was simulated for *n* = 50 chromosomes and a read depth of 250*X*, under mutation-drift equilibrium using msprime (***Kelleher et al., 2016***), with per-base crossover rates derived from *D. melanogaster* chromosome 2L (***Comeron et al., 2012***). Gray ribbons represent 95% confidence intervals. *R*^2^ is reported for the general linear model of predicted rates on true rates and mean absolute error was calculated across all 100 kb windows.

**Figure S8.**
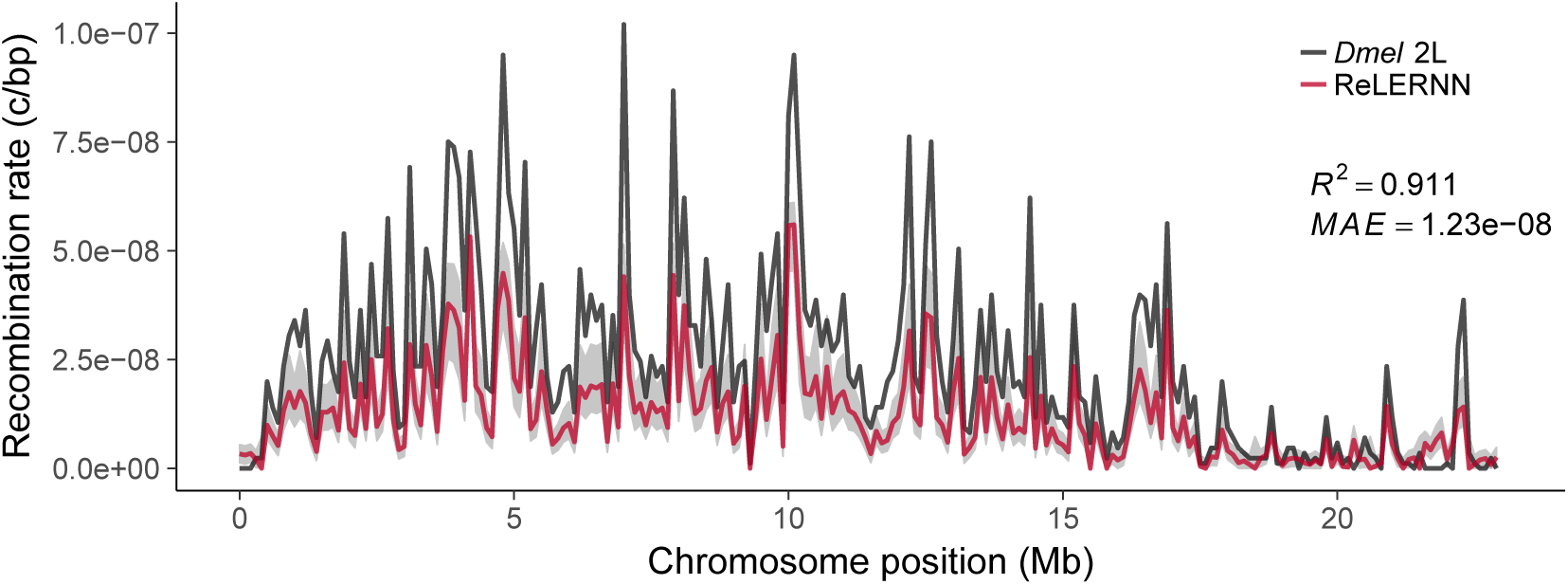
Recombination rate predictions for a simulated *Drosophila* chromosome (black line) using ReLERNN (red line). The recombination landscape was simulated for *n* = 20 chromosomes under mutation-drift equilibrium using msprime (***Kelleher et al., 2016***), with per-base crossover rates derived from *D. melanogaster* chromosome 2L (***Comeron et al., 2012***). Here the per-base mutation rate was assumed to be 50% less than the rate used for simulation. Gray ribbons represent 95% confidence intervals. *R*^2^ is reported for the general linear model of predicted rates on true rates and mean absolute error was calculated across all 100 kb windows.

**Figure S9.**
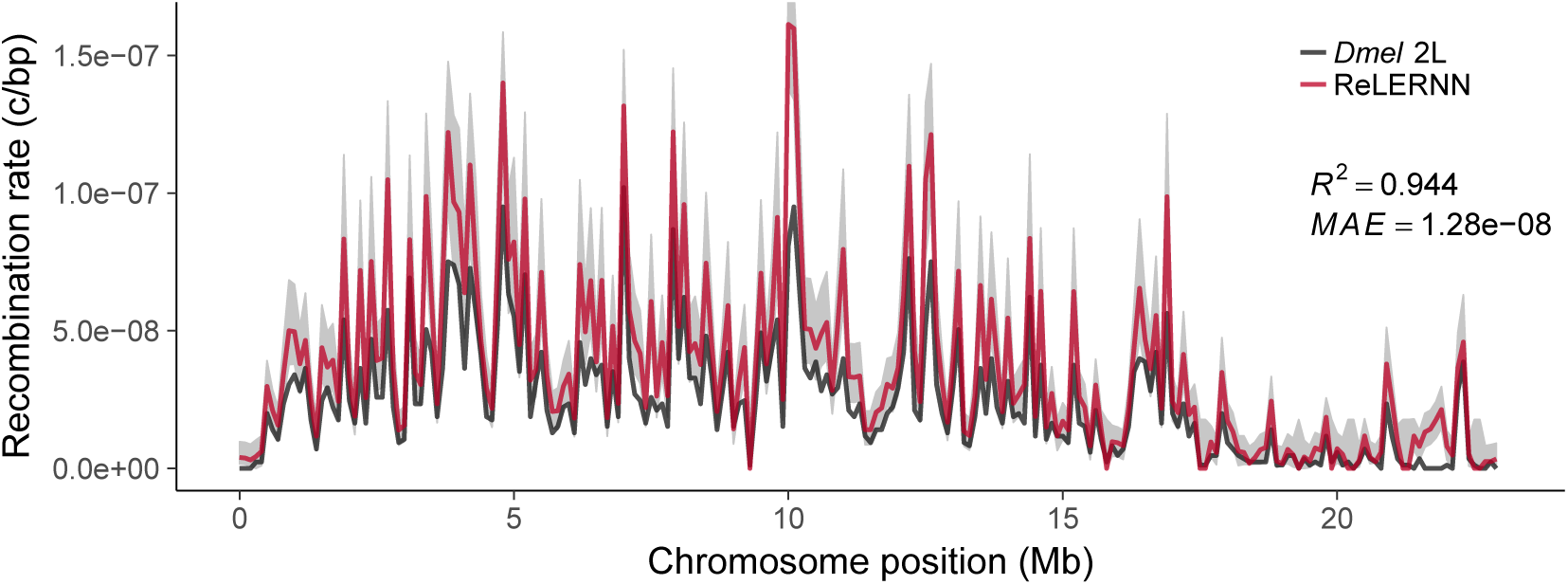
Recombination rate predictions for a simulated *Drosophila* chromosome (black line) using ReLERNN (red line). The recombination landscape was simulated for *n* = 20 chromosomes under mutation-drift equilibrium using msprime (***Kelleher et al., 2016***), with per-base crossover rates derived from *D. melanogaster* chromosome 2L (***Comeron et al., 2012***). Here the per-base mutation rate was assumed to be 50% greater than the rate used for simulation. Gray ribbons represent 95% confidence intervals. *R*^2^ is reported for the general linear model of predicted rates on true rates and mean absolute error was calculated across all 100 kb windows.

**Figure S10.**
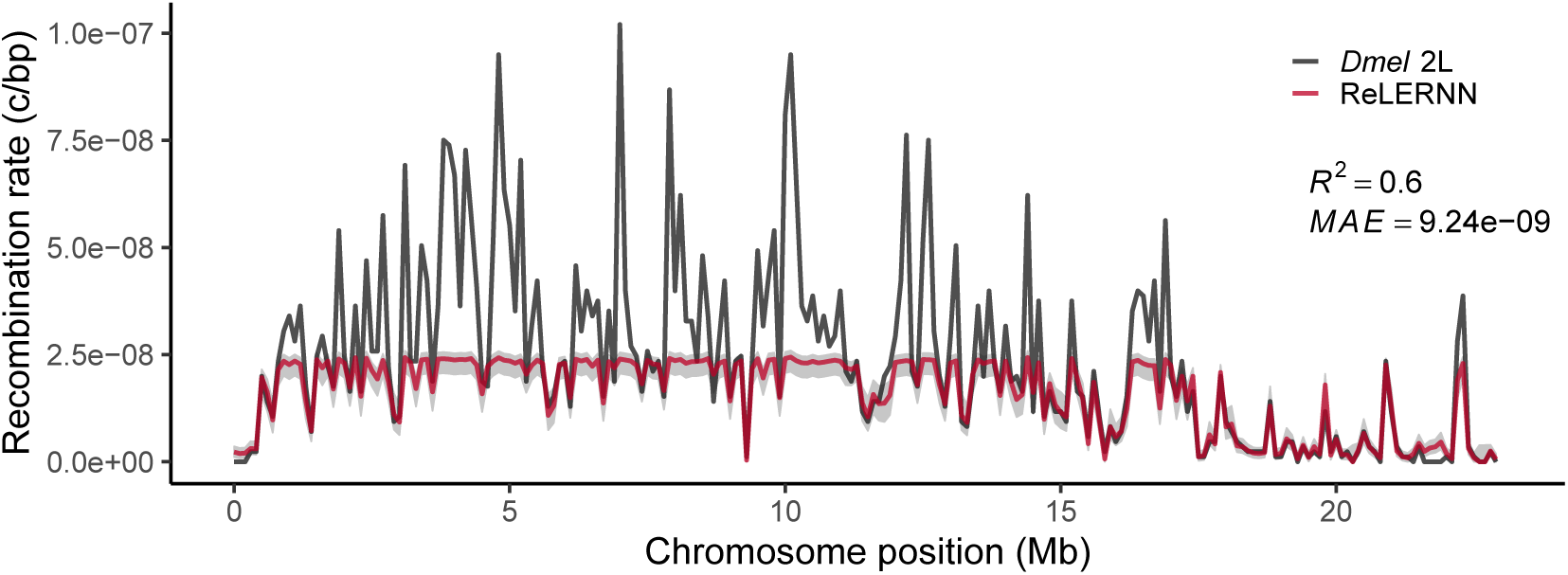
Recombination rate predictions for a simulated *Drosophila* chromosome (black line) using ReLERNN (red line). The recombination landscape was simulated for *n* = 20 chromosomes under mutation-drift equilibrium using msprime (***Kelleher et al., 2016***), with per-base crossover rates derived from *D. melanogaster* chromosome 2L (***Comeron et al., 2012***). Here the per-base mutation rate was assumed to be equal to the true rate, but *ρ*_*max*_ was assumed to be 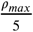. Gray ribbons represent 95% confidence intervals. *R*^2^ is reported for the general linear model of predicted rates on true rates and mean absolute error was calculated across all 100 kb windows.

**Figure S11.**
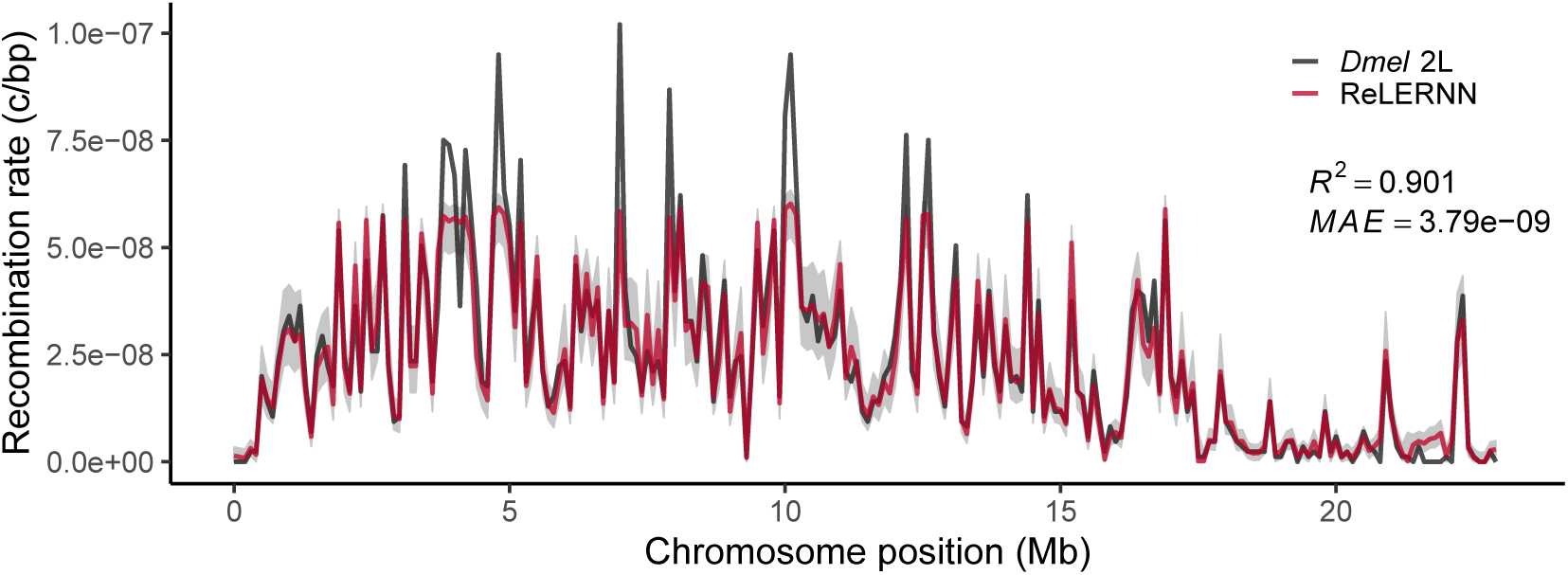
Recombination rate predictions for a simulated *Drosophila* chromosome (black line) using ReLERNN (red line). The recombination landscape was simulated for *n* = 20 chromosomes under mutation-drift equilibrium using msprime (***Kelleher et al., 2016***), with per-base crossover rates derived from *D. melanogaster* chromosome 2L (***Comeron et al., 2012***). Here the per-base mutation rate was assumed to be equal to the true rate, but *ρ*_*max*_ was assumed to be 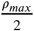. Gray ribbons represent 95% confidence intervals. *R*^2^ is reported for the general linear model of predicted rates on true rates and mean absolute error was calculated across all 100 kb windows.

**Figure S12.**
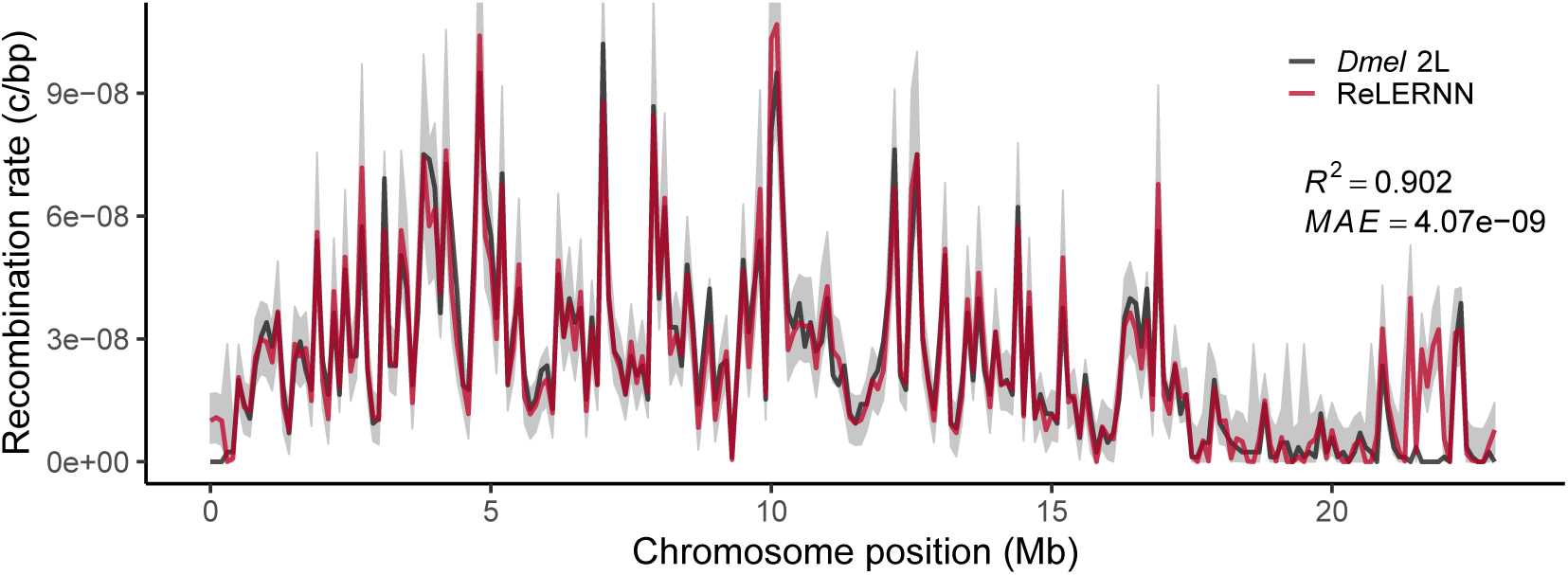
Recombination rate predictions for a simulated *Drosophila* chromosome (black line) using ReLERNN (red line). The recombination landscape was simulated for *n* = 20 chromosomes under mutation-drift equilibrium using msprime (***Kelleher et al., 2016***), with per-base crossover rates derived from *D. melanogaster* chromosome 2L (***Comeron et al., 2012***). Here the per-base mutation rate was assumed to be equal to the true rate, but *ρ*_*max*_ was assumed to be 2*ρ*_*max*_. Gray ribbons represent 95% confidence intervals. *R*^2^ is reported for the general linear model of predicted rates on true rates and mean absolute error was calculated across all 100 kb windows.

**Figure S13.**
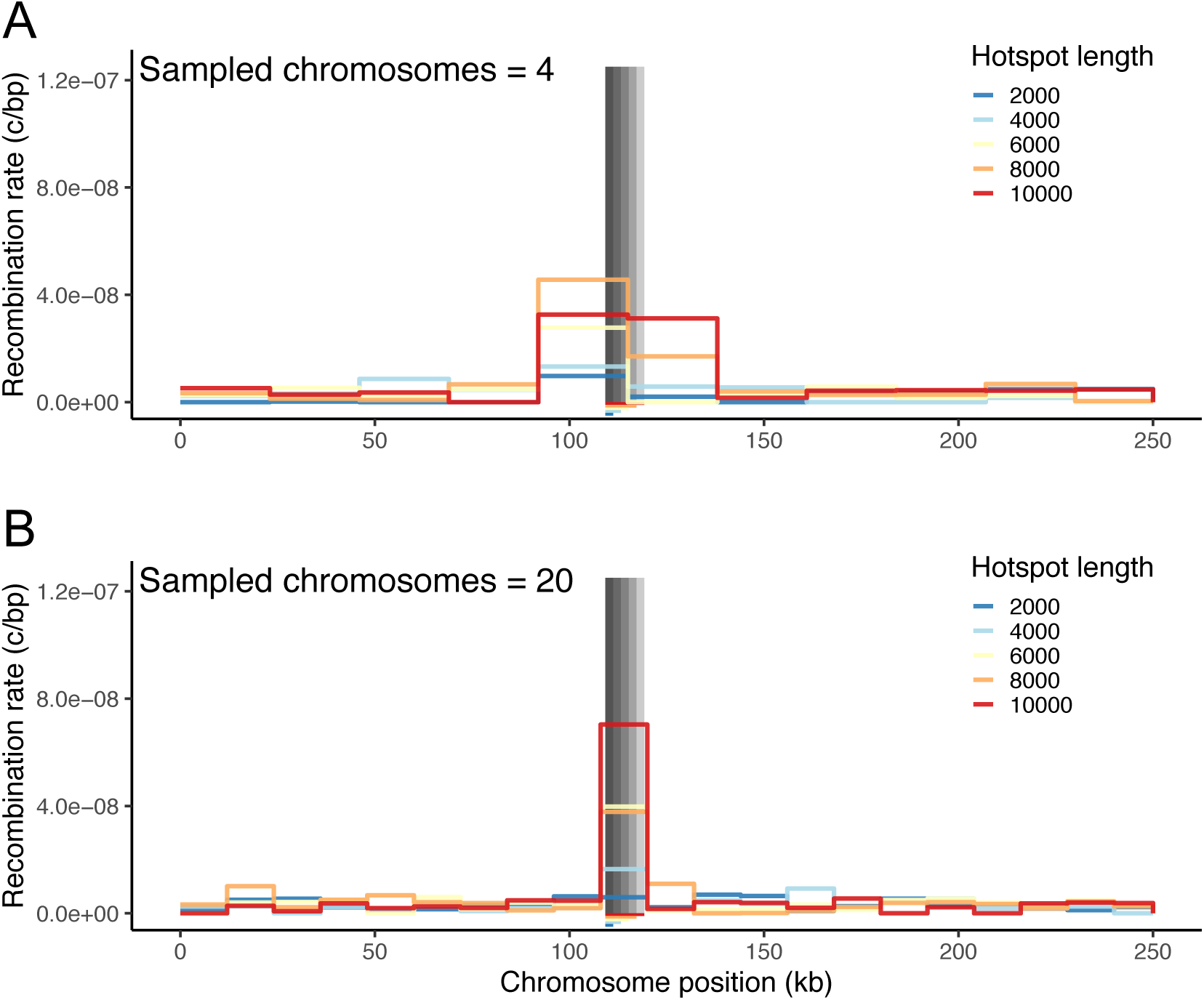
**(A)** Fine-scale rate predictions generated by ReLERNN for simulated recombination hot spots of varying lengths (*length* ∈ {2*kb*, 4*kb*, 6*kb*, 8*kb*, 10*kb*}, *r*_*background*_ = 2.5*e*^−9^, *r*_*hotspot*_ = 1.25*e*^−7^) for *n* =4 and **(B)** *n* = 20 chromosomes.

**Figure S14.**
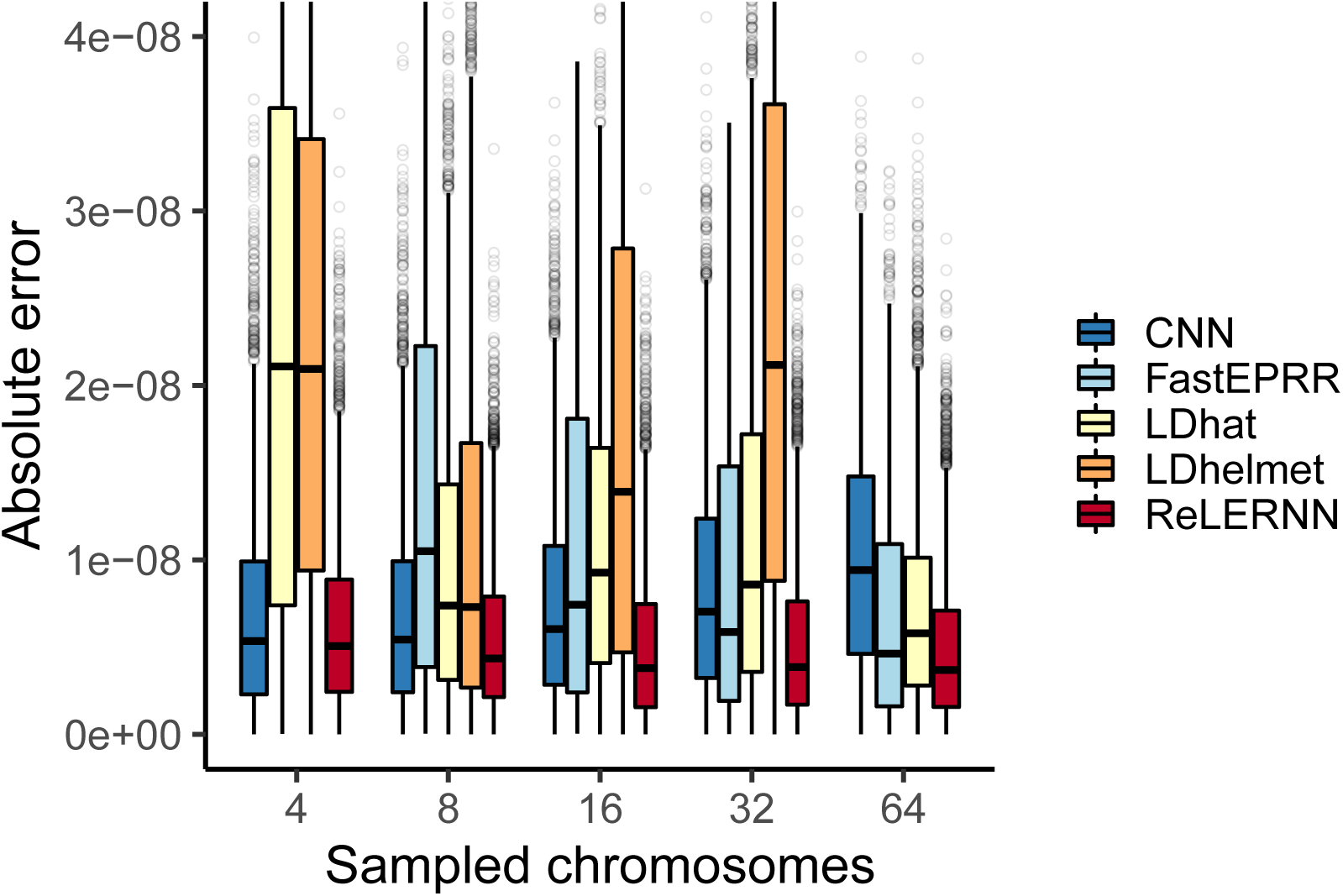
Distribution of absolute error (| *r* _*predicted*_ − *r* _*true*_|) for each method across 5000 simulated chromosomes (1000 for FastEPRR). Independent simulations were run under a model of demographic equilibrium. Sampled chromosomes indicate the number of independent sequences that were sampled from each msprime (***Kelleher et al., 2016***) coalescent simulation. LDhelmet was not able be used with *n* = 64 chromosomes, and FastEPRR was not able to be used with *n* = 4.

**Figure S15.**
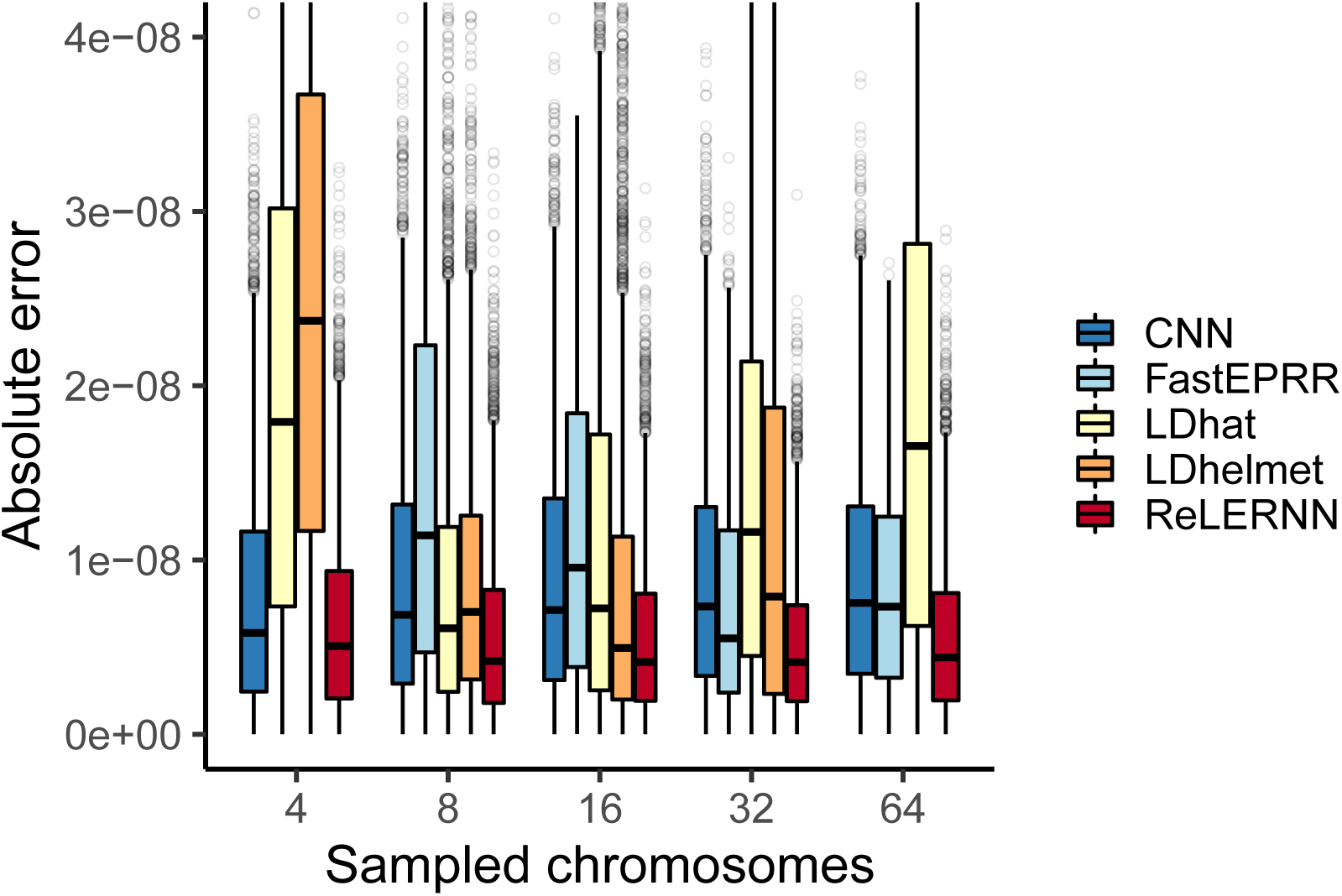
Distribution of absolute error (| *r* _*predicted*_ − *r* _*true*_|) for each method across 5000 simulated chromosomes (1000 for FastEPRR). Independent simulations were run under a model of population size expansion (see methods). Sampled chromosomes indicate the number of independent sequences that were sampled from each msprime (***Kelleher et al., 2016***) coalescent simulation. LDhelmet was not able be used with *n* = 64 chromosomes, and FastEPRR was not able to be used with *n* = 4.

**Figure S16.**
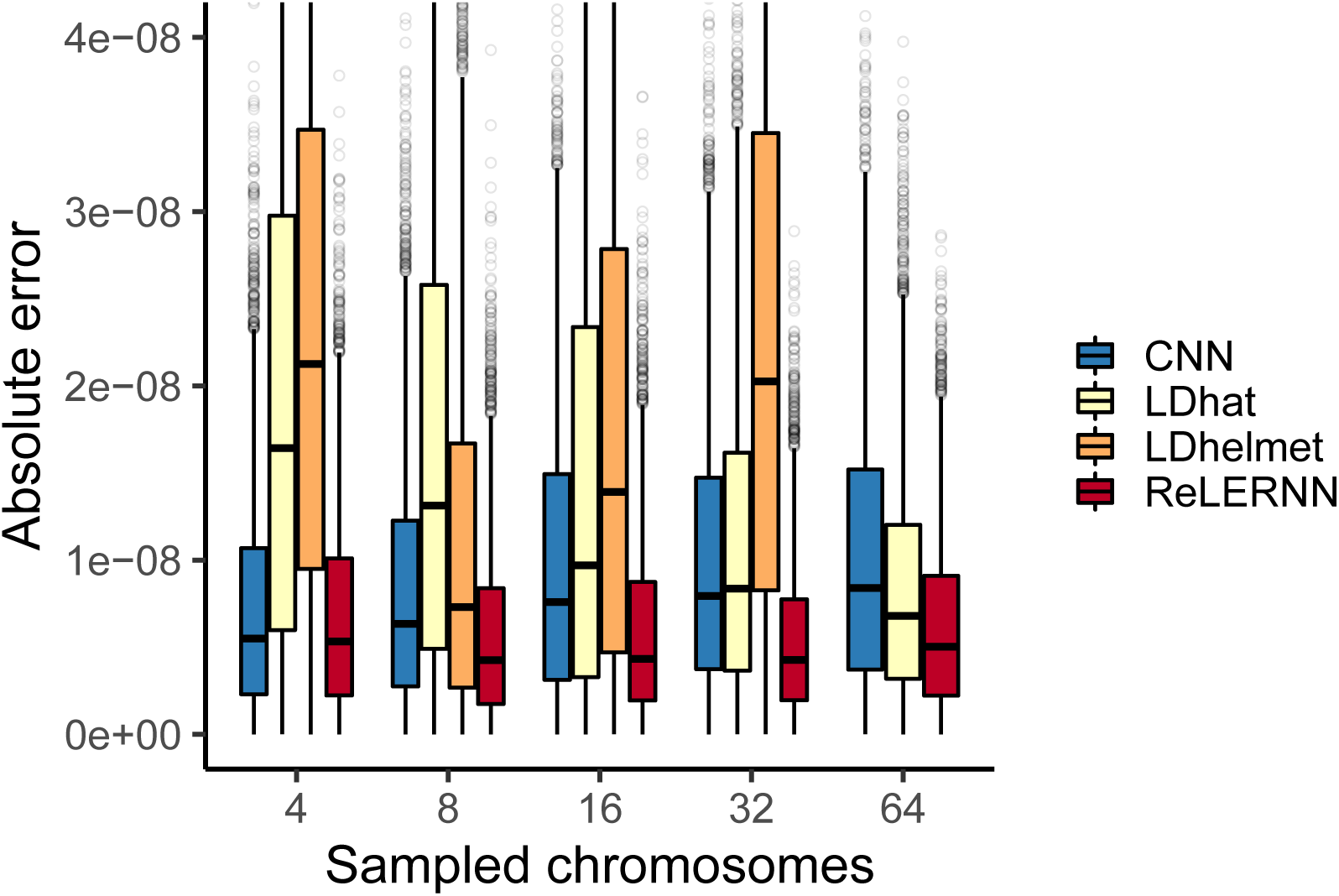
Distribution of absolute error (| *r* _*predicted*_ − *r* _*true*_|) for each method across 5000 simulated chromosomes after model misspecification. For the CNN and ReLERNN, predictions were made by training on demographic simulations while testing on sequences simulated under equilibrium. For LDhat and LDhelmet, the lookup tables were generated using parameters values that were estimated from simulations where the model was misspecified in the same way as described for the CNN and ReLERNN above. Sampled chromosomes indicate the number of independent sequences that were sampled from each msprime (***Kelleher et al., 2016***) coalescent simulation. LDhelmet was not able be used with *n* = 64 chromosomes and the demographic model could not be intentionally misspecified using FastEPRR.

**Figure S17.**
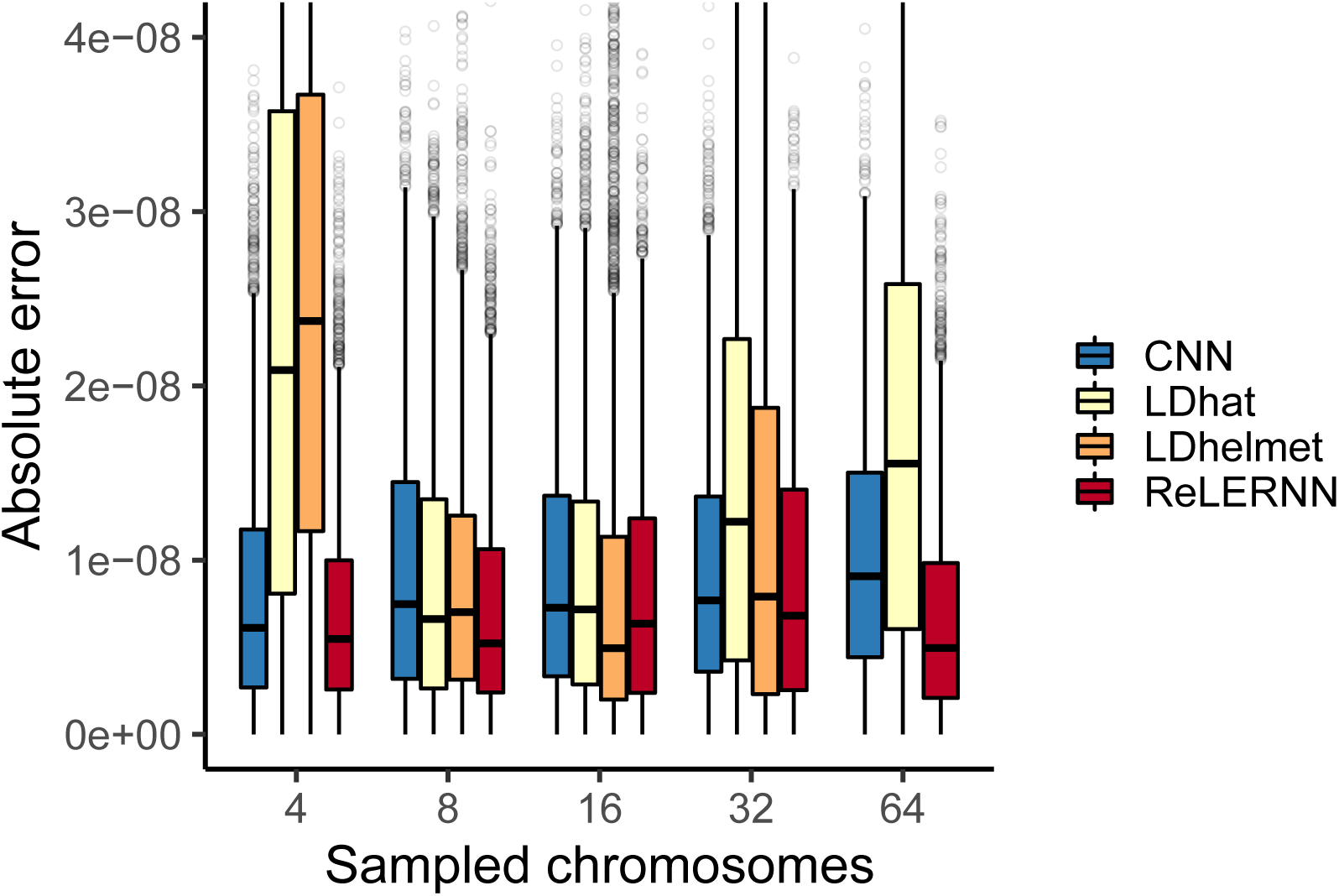
Distribution of absolute error (| *r* _*predicted*_ − *r* _*true*_|) for each method across 5000 simulated chromosomes after model misspecification. For the CNN and ReLERNN, predictions were made by training on equilibrium simulations while testing on sequences simulated under a model of population size expansion. For LDhat and LDhelmet, the lookup tables were generated using parameters values that were estimated from simulations where the model was misspecified in the same way as described for the CNN and ReLERNN above. Sampled chromosomes indicate the number of independent sequences that were sampled from each msprime (***Kelleher et al., 2016***) coalescent simulation. LDhelmet was not able be used with *n* = 64 chromosomes and the demographic model could not be intentionally misspecified using FastEPRR.

**Figure S18.**
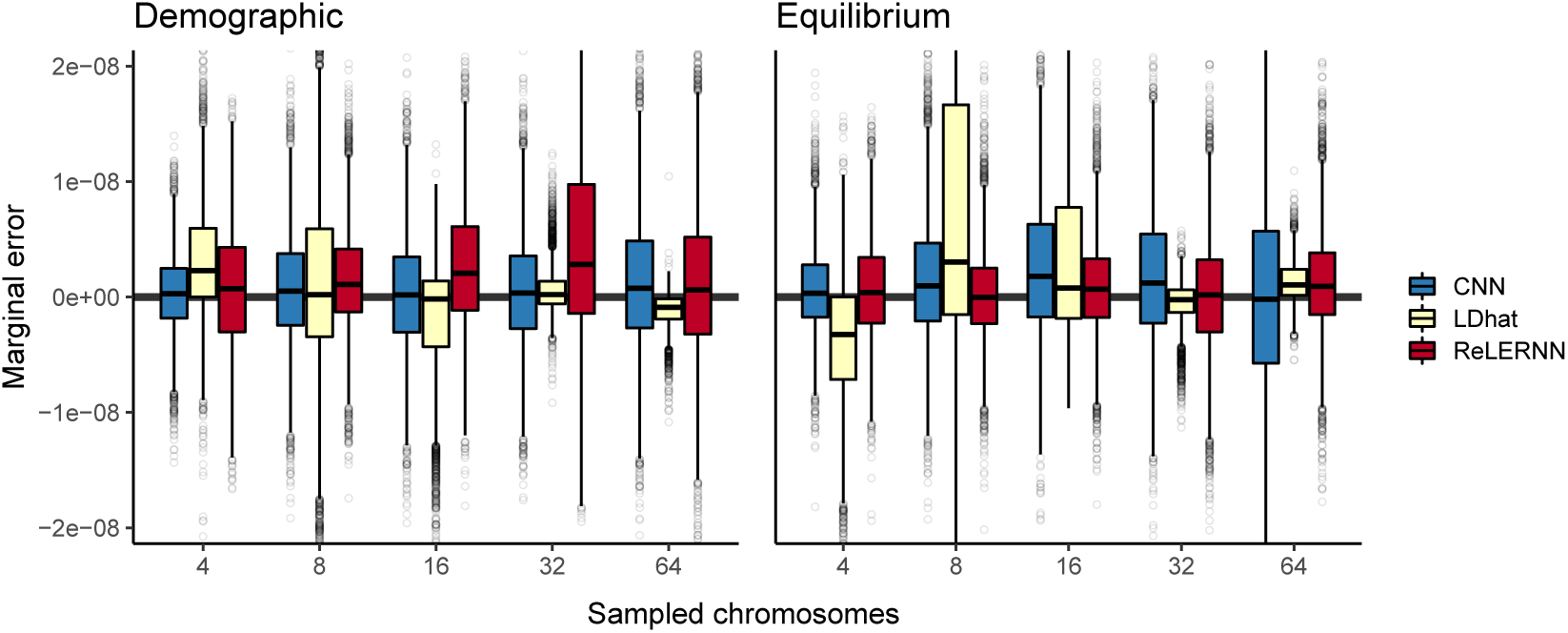
Distribution of marginal error attributed to model misspecification across 5000 simulated chromosomes. Predictions were made by training on equilibrium simulations and testing on sequences simulated under a demographic model **(left)** or training on demographic simulations and testing on sequences simulated under equilibrium **(right)**. Here, marginal error is represented as *ϵ*_*m*_ − *ϵ*_*c*_, where *ϵ*_*m*_ and *ϵ*_*c*_ are equal to (| *r* _*predicted*_ − *r* _*true*_|) when the model is misspecified and correctly specified, respectively. Sampled chromosomes indicate the number of independent sequences that were sampled from each msprime (***Kelleher et al., 2016***) coalescent simulation.

**Figure S19.**
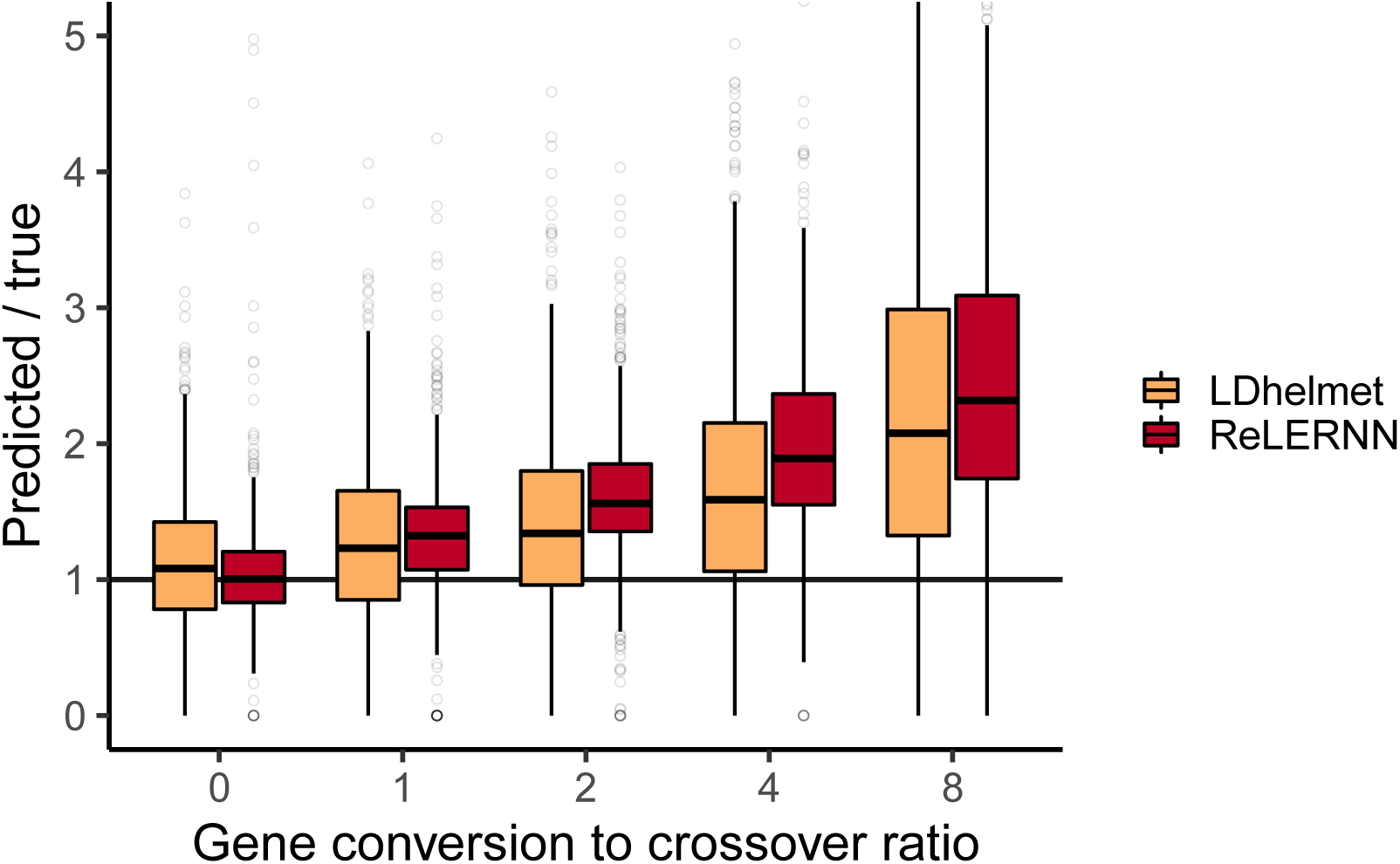
Distribution of predicted rates of recombination over true rates for 5000 examples simulated with gene conversion and *n* = 8. The ratio of gene conversion to crossovers was drawn from 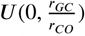, with 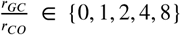. Gene conversion tract lengths were fixed at 352 bp. All simulations were completed in ms (***Hudson, 2002***).

**Figure S20.**
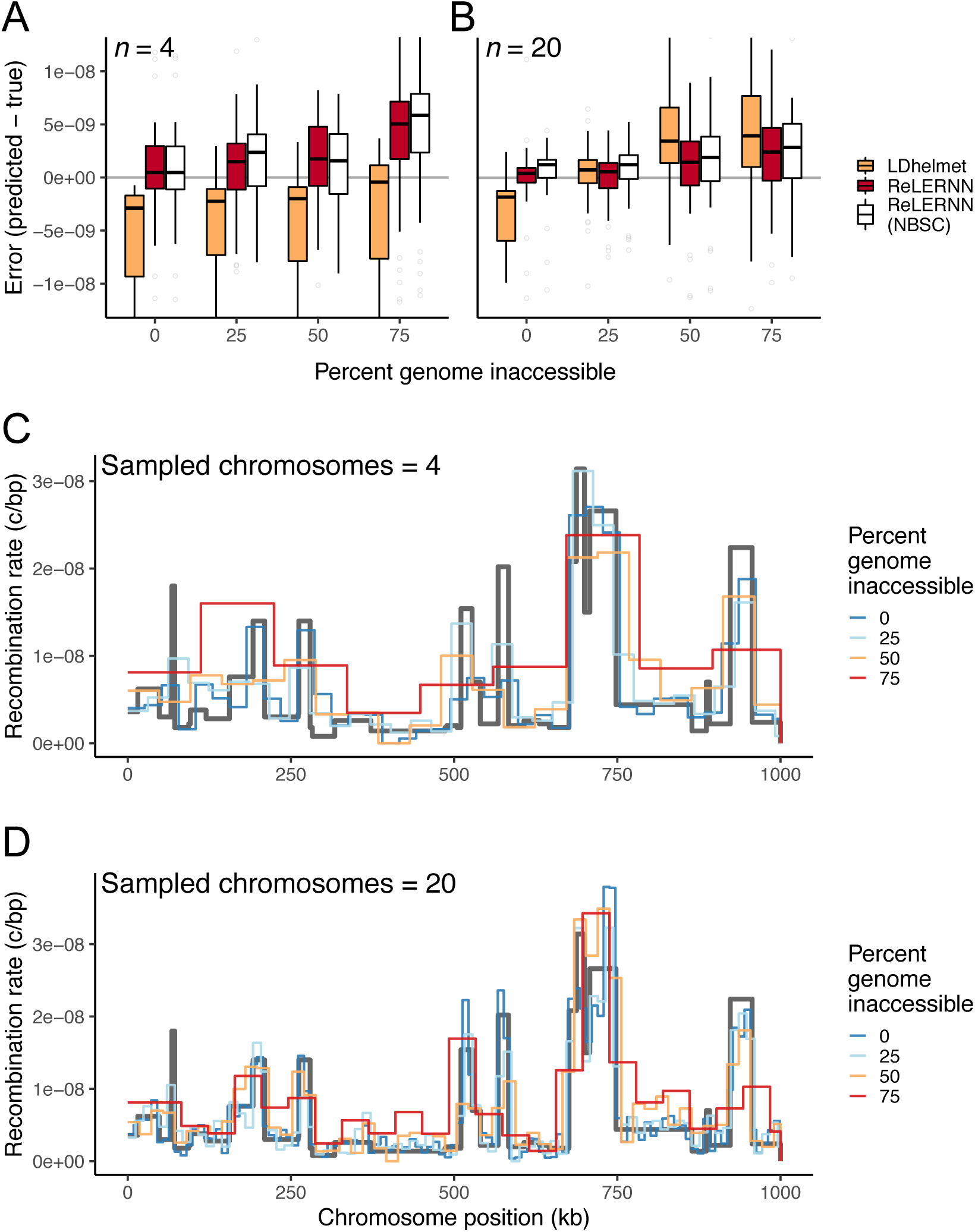
**(A)** Distribution of raw error (*r*_*predicted*_ − *r*_*true*_) for LDhelmet (***Chan et al., 2012***) and ReL-ERNN when presented with varying levels of genome inaccessibility for simulations with *n* =4 and **(B)** *n* = 20 chromosomes. **(C)** Fine-scale rate predictions generated by ReLERNN for a recombination landscape (grey line) simulated with varying levels of genome inaccessibility, for *n* = 4 and **(D)** *n* = 20 chromosomes.

**Figure S21.**
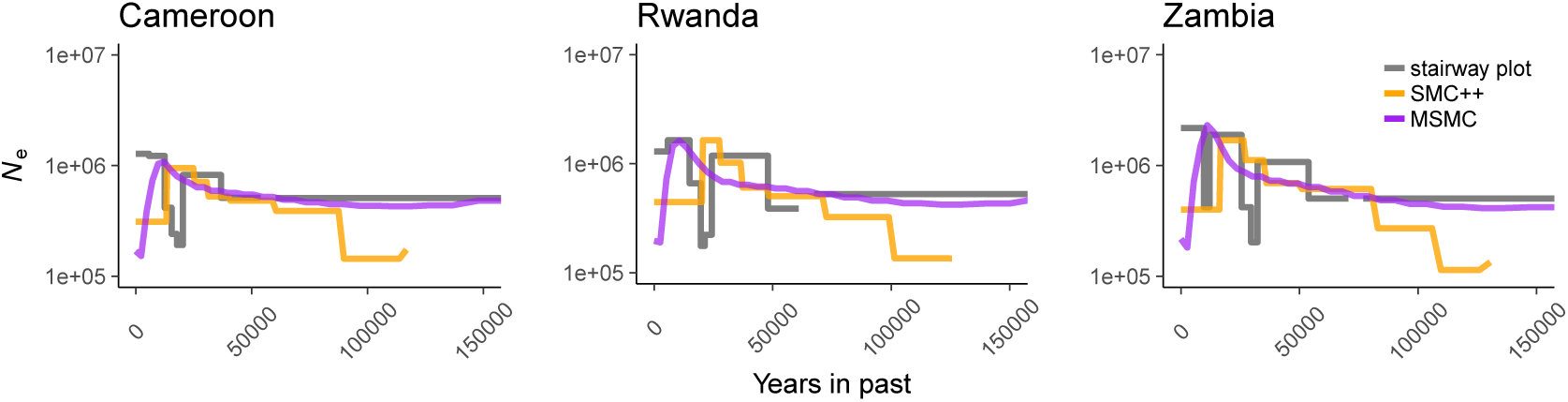
Historical population size estimates were inferred for Cameroon, Rwanda, and Zambia using three separate methods, all of which disagree with one another. Inferences are based on 10 samples for both stairwayplot (grey line) and SMC++ (orange line), and 2 samples for MSMC (purple line).

**Figure S22.**
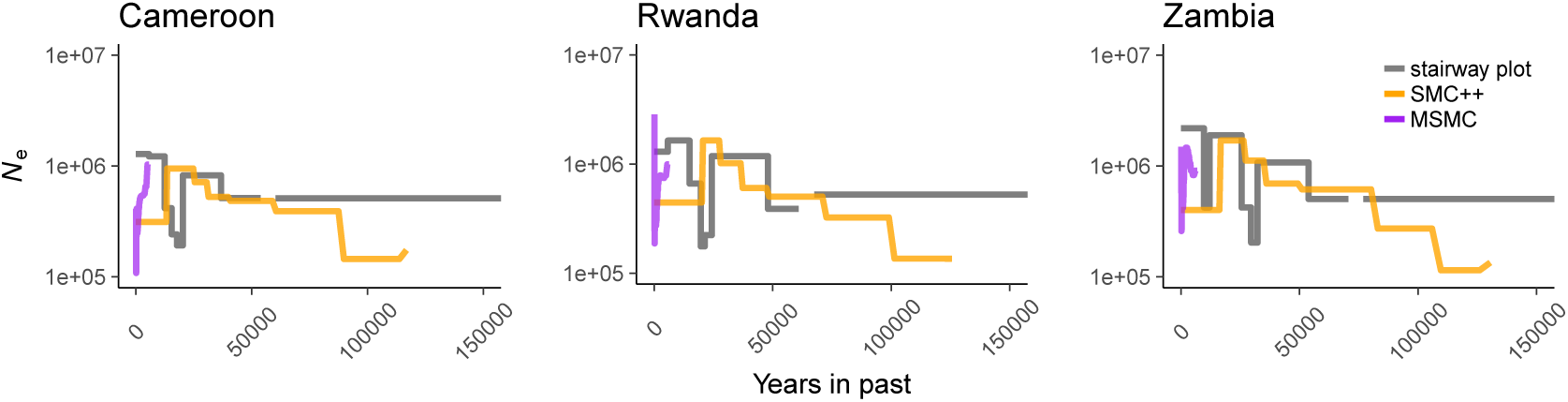
Historical population size estimates were inferred for Cameroon, Rwanda, and Zambia using three separate methods. Here, inferences are based on 10 samples for both stairway-plot (grey line) and SMC++ (orange line), and 10 samples for MSMC (purple line).

**Figure S23.**
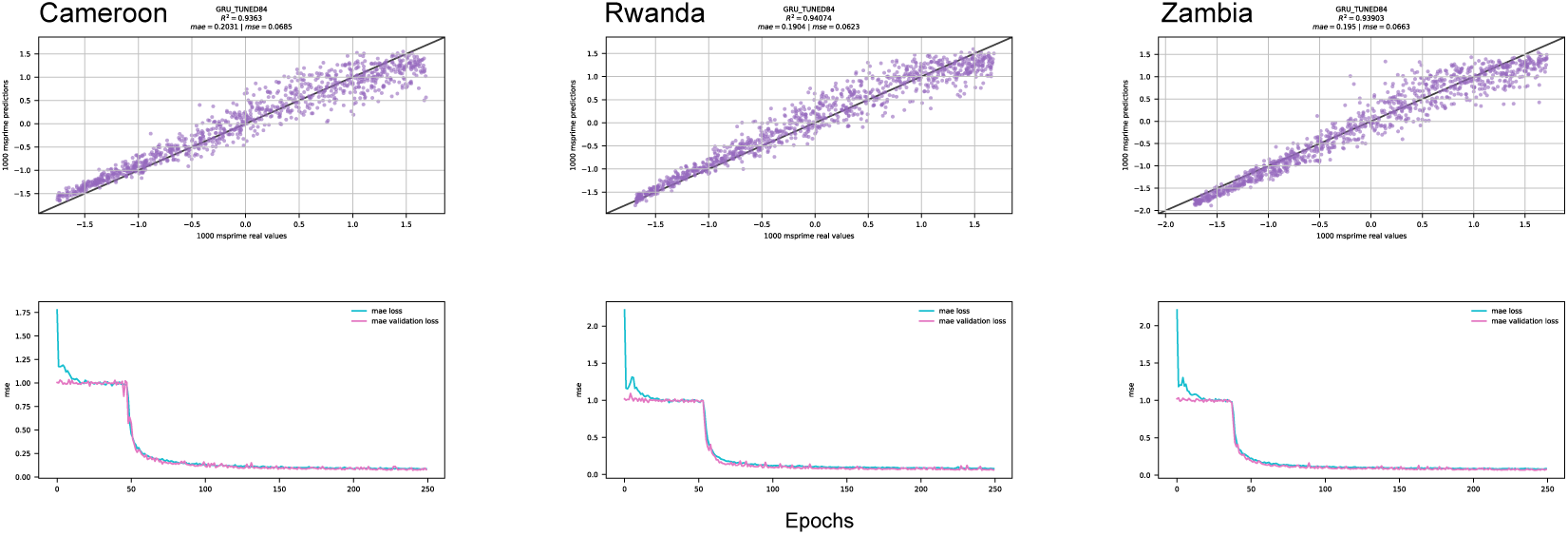
ReLERNN test results for Cameroon, Rwanda, and Zambia when trained under assumptions of mutation-drift equilibrium. **(Top)** Scatter plot of raw (unnormalized) predictions for 1000 test examples using ReLERNN with the same parameters used in ***Figure 2***. Mean absolute error and mean squared error are shown for each population. **(Bottom)** Line graph showing the convergence of loss (measured by mean squared error) over time (epochs) during training on the same data as above, for both the training set (blue lines) and the validation set (purple lines).

**Figure S24.**
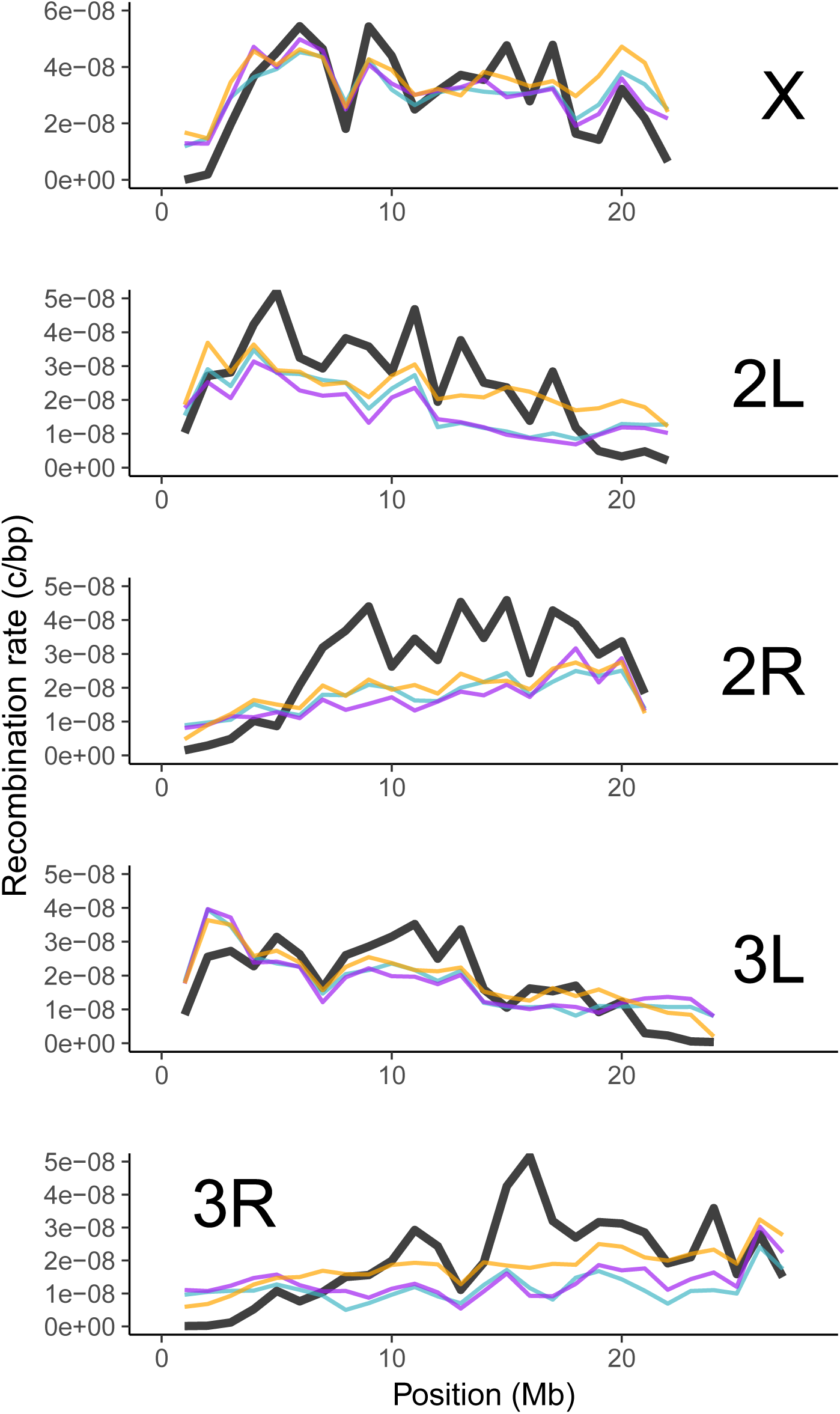
Genome-wide recombination landscapes for *D. melanogaster* populations from Cameroon (teal lines), Rwanda (purple lines), and Zambia (orange lines). Rates are compared to those experimentally derived by ***Comeron et al***. (***2012***) (black lines). All rates have been scales to 1 Mb windows by using a weighted average (see Materials and Methods). Sample sizes (*n* = 10) are the same for all populations.

**Figure S25.**
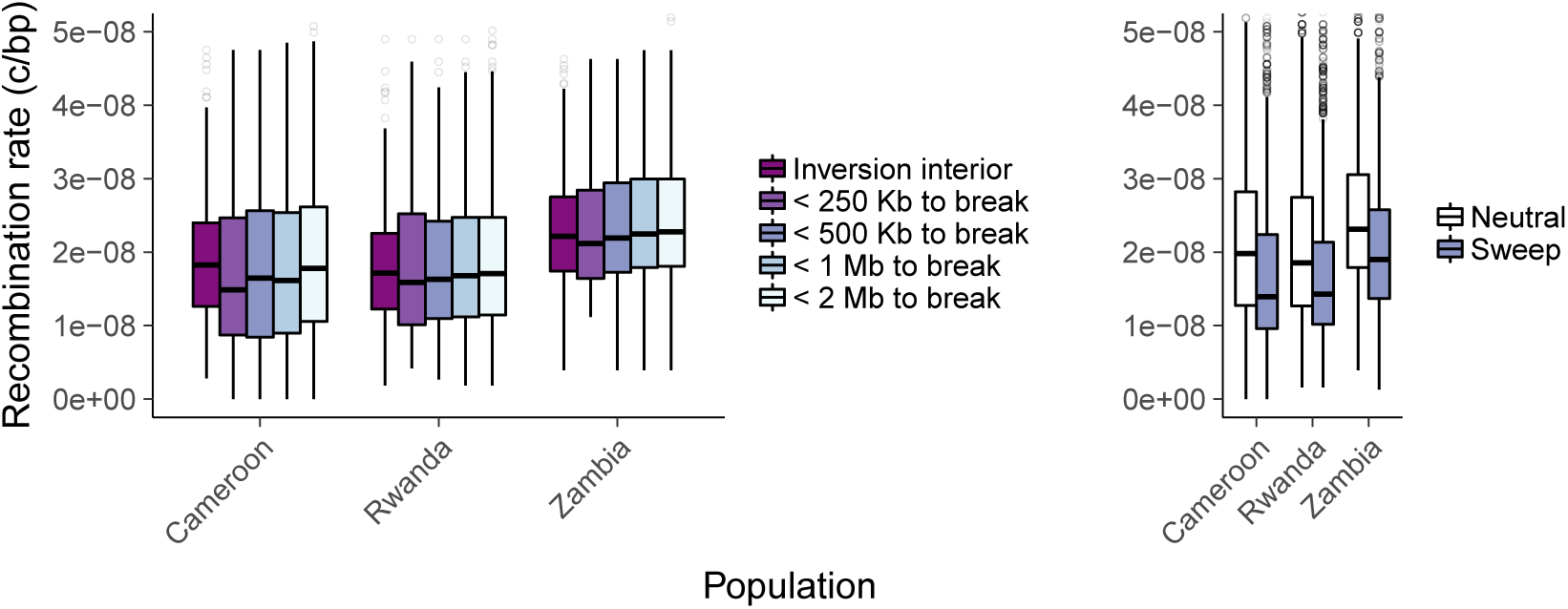
**(Left)** Recombination rate estimates for genomic windows > 2 Mb inside, < 250 kb surrounding, < 500 kb surrounding, < 1 Mb surrounding, and <2 Mb surrounding all inversion breakpoints. **(Right)** Recombination rate estimates for all genomic windows overlapping windows predicted as either hard/soft sweeps (purple) or as neutral (white) by diploS/HIC (***Kern and Schrider, 2018***).

**Figure S26.**
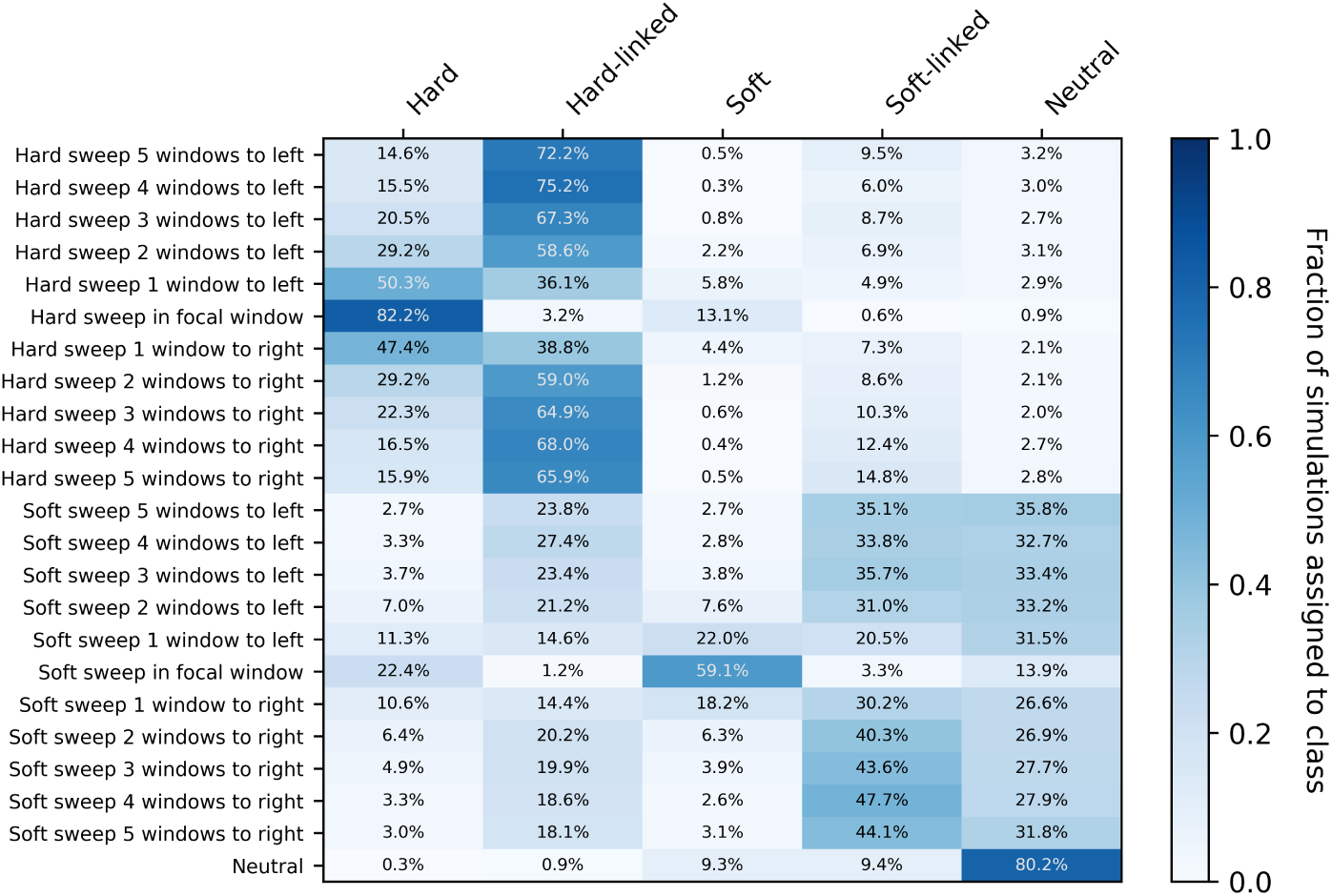
Confusion matrix showing the fraction of test simulation windows assigned to each of five prediction categories by diploS/HIC (***Kern and Schrider, 2018***): hard, hard-linked, soft, soft-linked, and neutral. The y-axis shows the location of the window being classified relative to the selected window.

**Figure S27.**
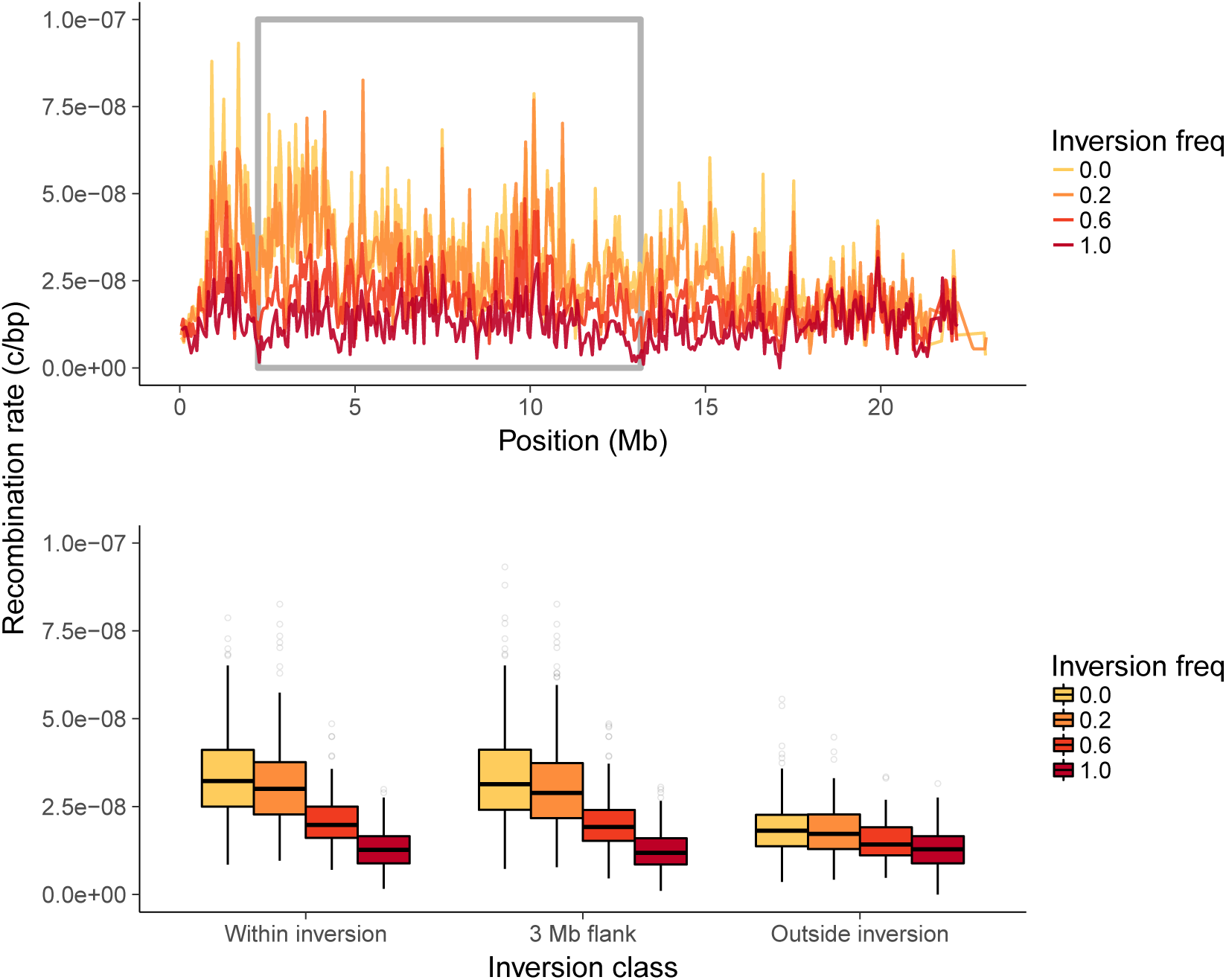
**(Top)** Recombination landscapes for Zambian *D. melanogaster* surrounding *In(2L)t*, sampled at different inversion frequencies. The grey box denotes the inversion boundaries of *In(2L)t* in *Drosophila* (***Corbett-Detig and Hartl, 2012***). **(Bottom)** Recombination rate estimates from genomic windows within the inversion, within a 3 Mb region flanking the inversion, and 3 Mb outside the inversion, sampled at different inversion frequencies.

**Figure S28.**
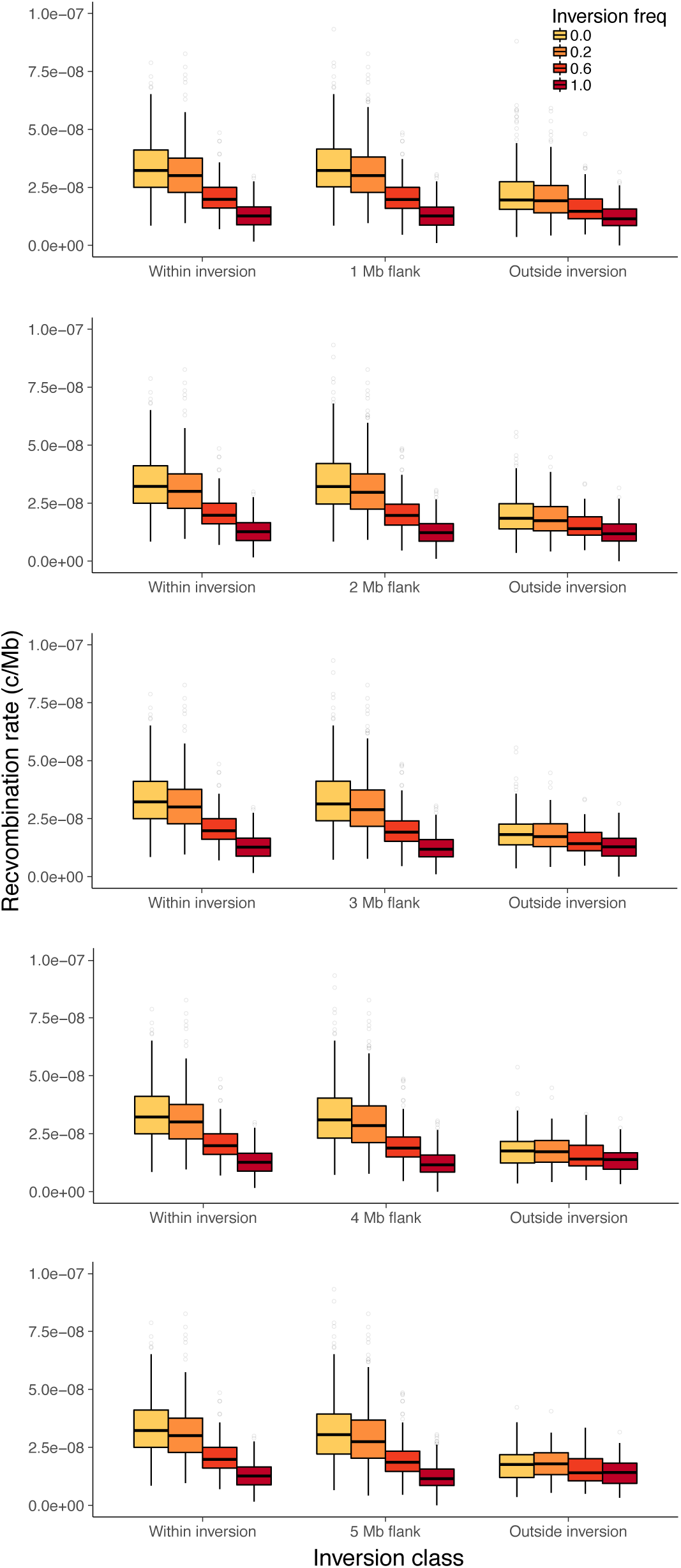
Recombination rate estimates using flanking window sizes from 1-5 Mb. Rates are shown for genomic windows within the inversion, within regions flanking the inversion, and for regions outside both the inversion and flanking regions. All estimates are from chromosome 2L with *In(2L)t* sampled at different inversion frequencies

## References

Abadi M, Agarwal A, Barham P, Brevdo E, Chen Z, Citro C, Corrado GS, Davis A, Dean J, Devin M, Ghemawat S, Goodfellow I, Harp A, Irving G, Isard M, Jia Y, Jozefowicz R, Kaiser L, Kudlur M, Levenberg J, et al., TensorFlow: Large-Scale Machine Learning on Heterogeneous Systems; 2015. https://www.tensorflow.org/, software availablefrom tensorflow.org.

Aulard S, David JR, Lemeunier F. Chromosomal inversion polymorphism in Afrotropical populations of Drosophila melanogaster. Genetics Research. 2002; 79(1):49–63.

Ayala D, Guerrero RF, Kirkpatrick M. Reproductive isolation and local adaptation quantified for a chromosome inversion in a malaria mosquito. Evolution: International Journal of Organic Evolution. 2013; 67(4):946–958.

Barton N. A general model for the evolution of recombination. Genetics Research. 1995; 65(2):123–144.

Brandvain Y, Kenney AM, Flagel L, Coop G, Sweigart AL. Speciation and introgression between Mimulus nasutus and Mimulus guttatus. PLoS genetics. 2014; 10(6):e1004410.

Bulik-Sullivan BK, Loh PR, Finucane HK, Ripke S, Yang J, Patterson N, Daly MJ, Price AL, Neale BM, of the Psychiatric Genomics Consortium SWG, et al. LD Score regression distinguishes confounding from polygenicity in genome-wide association studies. Nature genetics. 2015; 47(3):291.

Chan AH, Jenkins PA, Song YS. Genome-wide fine-scale recombination rate variation in Drosophila melanogaster. PLoS genetics. 2012; 8(12):e1003090.

Chan J, Perrone V, Spence JP, Jenkins PA, Mathieson S, Song YS. A Likelihood-Free Inference Framework for Population Genetic Data using Exchangeable Neural Networks. bioRxiv. 2018; https://www.biorxiv.org/content/early/2018/11/05/267211, doi: 10.1101/267211.

Charlesworth B. Recombination modification in a fluctuating environment. Genetics. 1976; 83(1):181–195.

Cho K, Van Merriënboer B, Bahdanau D, Bengio Y. On the properties of neural machine translation: Encoder-decoder approaches. arXiv preprint 14091259. 2014;.

Chollet F, et al., Keras. GitHub; 2015. https://github.com/fchollet/keras.

Chung J, Gulcehre C, Cho K, Bengio Y. Empirical evaluation of gated recurrent neural networks on sequence modeling. arXiv preprint 14123555. 2014;.

Comeron JM, Ratnappan R, Bailin S. The Many Landscapes of Recombination in Drosophila melanogaster. PLOS Genetics. 2012 10; 8(10):1–21. https://doi.org/10.1371/journal.pgen.1002905, doi: 10.1371/journal.pgen.1002905.

Consortium GP, et al. A global reference for human genetic variation. Nature. 2015; 526(7571):68.

Corbett-Detig RB, Hartl DL. Population Genomics of Inversion Polymorphisms in Drosophila melanogaster. PLOS Genetics. 2012 12; 8(12):1–15. https://doi.org/10.1371/journal.pgen.1003056, doi: 10.1371/journal.pgen.1003056.

Do AT, Brooks JT, Le Neveu MK, LaRocque JR. Double-strand break repair assays determine pathway choice and structure of gene conversion events in Drosophila melanogaster. G3: Genes, Genomes, Genetics. 2014; 4(3):425–432.

Dobzhansky T. Genetics and the origin of species. Genetics and the origin of species. 1937;.

Dobzhansky T, Epling C. The suppression of crossing over in inversion heterozygotes of Drosophila pseudoobscura. Proceedings of the National Academy of Sciences of the United States of America. 1948; 34(4):137.

Elyashiv E, Sattath S, Hu TT, Strutsovsky A, McVicker G, Andolfatto P, Coop G, Sella G. A genomic map of the effects of linked selection in Drosophila. PLoS genetics. 2016; 12(8):e1006130.

Feder AF, Petrov DA, Bergland AO. LDx: estimation of linkage disequilibrium from high-throughput pooled resequencing data. PloS one. 2012; 7(11):e48588.

Fisher R. The genetical theory of natural selection. 1930;.

Flagel L, Brandvain Y, Schrider DR. The Unreasonable Effectiveness of Convolutional Neural Networks in Population Genetic Inference. Molecular Biology and Evolution. 2018 12; 36(2):220–238. https://dx.doi.org/10.1093/molbev/msy224, doi: 10.1093/molbev/msy224.

Fuller ZL, Koury SA, Leonard CJ, Young RE, Ikegami K, Westlake J, Richards S, Schaeffer SW, Phadnis N. Extensive recombination suppression and chromosome-wide differentiation of a segregation distorter in Drosophila. bioRxiv. 2018; https://www.biorxiv.org/content/early/2018/12/21/504126, doi: 10.1101/504126.

Gao F, Ming C, Hu W, Li H. New software for the fast estimation of population recombination rates (FastEPRR) in the genomic era. G3: Genes, Genomes, Genetics. 2016; 6(6):1563–1571.

Gay J, Myers S, McVean G. Estimating meiotic gene conversion rates from population genetic data. Genetics. 2007; 177(2):881–894.

Gravel S, Henn BM, Gutenkunst RN, Indap AR, Marth GT, Clark AG, Yu F, Gibbs RA, Bustamante CD. Demographic history and rare allele sharing among human populations. Proceedings of the National Academy of Sciences. 2011; 108(29):11983–11988. https://www.pnas.org/content/108/29/11983, doi: 10.1073/pnas.1019276108.

Graves A, Jaitly N, Mohamed A. Hybrid speech recognition with Deep Bidirectional LSTM. In: 2013 IEEE Workshop on Automatic Speech Recognition and Understanding, Olomouc, Czech Republic, December 8-12, 2013; 2013. p. 273–278. https://doi.org/10.1109/ASRU.2013.6707742, doi: 10.1109/ASRU.2013.6707742.

Hahn MW. Molecular population genetics. Sinauer Associates; 2018.

Hill WG, Robertson A. The effect of linkage on limits to artificial selection. Genetics Research. 1966; 8(3):269–294.

Hilliker A J, Harauz G, Reaume AG, Gray M, Clark SH, Chovnick A. Meiotic gene conversion tract length distribution within the rosy locus of Drosophila melanogaster. Genetics. 1994; 137(4):1019–1026.

Hinch AG, Tandon A, Patterson N, Song Y, Rohland N, Palmer CD, Chen GK, Wang K, Buxbaum SG, Akylbekova EL, et al. The landscape of recombination in African Americans. Nature. 2011; 476(7359):170.

Hinton G, Deng L, Yu D, Dahl G, rahman Mohamed A, Jaitly N, Senior A, Vanhoucke V, Nguyen P, Sainath T, Kingsbury B. Deep Neural Networks for Acoustic Modeling in Speech Recognition. Signal Processing Magazine. 2012;.

Hudson RR. Estimation the recombination parameter of a finite population model without selection. Genetical Research. 1987; 50:245–250.

Hudson RR. Generating samples under a Wright-Fisher neutral model of genetic variation. Bioinformatics. 2002 Feb; 18(2):337–338.

Hudson RR, Kaplan NL. Statistical properties of the number of recombination events in the history of a sample of DNA sequences. Genetics. 1985; 111(1):147–164.

Jaenike J. Sex chromosome meiotic drive. Annual Review of Ecology and Systematics. 2001; 32(1):25–49.

Jeffreys AJ, Kauppi L, Neumann R. Intensely punctate meiotic recombination in the class II region of the major histocompatibility complex. Nature genetics. 2001; 29(2):217.

Jeffreys AJ, May CA. Intense and highly localized gene conversion activity in human meiotic crossover hot spots. Nature genetics. 2004; 36(2):151.

Jozefowicz R, Zaremba W, Sutskever I. An empirical exploration of recurrent network architectures. In: International Conference on Machine Learning; 2015. p. 2342–2350.

Kelleher J, Etheridge AM, McVean G. Efficient Coalescent Simulation and Genealogical Analysis for Large Sample Sizes. PLOS Computational Biology. 2016 May; 12(5):e1004842. https://doi.org/10.1371/journal.pcbi.1004842, doi: 10.1371/journal.pcbi.1004842.

Kern AD, Schrider DR. Discoal: flexible coalescent simulations with selection. Bioinformatics. 2016; 32(24):3839–3841.

Kern AD, Schrider DR. diploS/HIC: An Updated Approach to Classifying Selective Sweeps. G3: Genes, Genomes, Genetics. 2018; 8(6):1959–1970. http://www.g3journal.org/content/8/6/1959, doi: 10.1534/g3.118.200262.

Kim Y, Nielsen R. Linkage disequilibrium as a signature of selective sweeps. Genetics. 2004; 167(3):1513–1524.

Kingma DP, Ba J. Adam: A method for stochastic optimization. arXiv preprint 14126980. 2014;.

Kirkpatrick M, Barton N. Chromosome inversions, local adaptation and speciation. Genetics. 2006; 173(1):419–434.

Kong A, Thorleifsson G, Gudbjartsson DF, Masson G, Sigurdsson A, Jonasdottir A, Walters GB, Jonasdottir A, Gylfason A, Kristinsson KT, et al. Fine-scale recombination rate differences between sexes, populations and individuals. Nature. 2010; 467(7319):1099.

Krizhevsky A, Sutskever I, Hinton GE. ImageNet Classification with Deep Convolutional Neural Networks. In: Pereira F, Burges CJC, Bottou L, Weinberger KQ, editors. Advances in Neural Information Processing Systems 25 Curran Associates, Inc.; 2012. p. 1097–1105. http://papers.nips.cc/paper/4824-imagenet-classification-with-deep-convolutional-neural-networks.pdf.

Kulathinal RJ, Stevison LS, Noor MA. The genomics of speciation in Drosophila: diversity, divergence, and introgression estimated using low-coverage genome sequencing. PLoS genetics. 2009; 5(7):e1000550.

Lack JB, Cardeno CM, Crepeau MW, Taylor W, Corbett-Detig RB, Stevens KA, Langley CH, Pool JE. The Drosophila Genome Nexus: A Population Genomic Resource of 623 Drosophila melanogaster Genomes, Including 197 from a Single Ancestral Range Population. Genetics. 2015; 199(4):1229–1241. http://www.genetics.org/content/199/4/1229, doi: 10.1534/genetics.115.174664.

Langley CH, Stevens K, Cardeno C, Lee YCG, Schrider DR, Pool JE, Langley SA, Suarez C, Corbett-Detig RB, Kolaczkowski B, et al. Genomic variation in natural populations of Drosophila melanogaster. Genetics. 2012; 192(2):533–598.

Lecun Y, Bottou L, Bengio Y, Haffner P. Gradient-based learning applied to document recognition. In: Proceedings of the IEEE; 1998. p. 2278–2324.

Lemeunier F, Aulard S. Inversion polymorphism in Drosophila melanogaster. Drosophila inversion polymorphism. Boca Raton (FL): CRC Press; 1992.

Lewontin R, Kojima Ki. The evolutionary dynamics of complex polymorphisms. Evolution. 1960; 14(4):458–472.

Li N, Stephens M. Modeling linkage disequilibrium and identifying recombination hotspots using singlenucleotide polymorphism data. Genetics. 2003; 165(4):2213–2233.

Lichten M. Meiotic recombination: breaking the genome to save it. Current Biology. 2001; 11(7):R253–R256.

Lin K, Futschik A, Li H. A fast estimate for the population recombination rate based on regression. Genetics. 2013; p. genetics–113.

Liu X, Fu YX. Exploring population size changes using SNP frequency spectra. Nature Genetics. 2015 04; 47:555 EP –. https://doi.org/10.1038/ng.3254.

McVean G, Awadalla P, Fearnhead P. A coalescent-based method for detecting and estimating recombination from gene sequences. Genetics. 2002; 160(3):1231–1241.

Miller DE, Cook KR, Arvanitakis AV, Hawley RS. Third Chromosome Balancer Inversions Disrupt Protein-Coding Genes and Influence Distal Recombination Events in Drosophila melanogaster. G3: Genes, Genomes, Genetics. 2016; 6(7):1959–1967. https://www.g3journal.org/content/6/7/1959, doi: 10.1534/g3.116.029330.

Muller HJ. Some genetic aspects of sex. The American Naturalist. 1932; 66(703):118–138.

Myers S, Bottolo L, Freeman C, McVean G, Donnelly P. A fine-scale map of recombination rates and hotspots across the human genome. Science. 2005; 310(5746):321–324.

Myers SR, Griffiths RC. Bounds on the minimum number of recombination events in a sample history. Genetics. 2003; 163(1):375–394.

Nicklas RB. Chromosome segregation mechanisms. Genetics. 1974; 78(1):205–213.

Noor MA, Grams KL, Bertucci LA, Reiland J. Chromosomal inversions and the reproductive isolation of species. Proceedings of the National Academy of Sciences. 2001; 98(21):12084–12088.

Novitski E, Braver G. An analysis of crossing over within a heterozygous inversion in Drosophila melanogaster. Genetics. 1954; 39(2):197.

Ohta T, Kimura M. Linkage disequilibrium due to random genetic drift. Genetics Research. 1969; 13(1):47–55.

Ohta T, Kimura M. Development of associative overdominance through linkage disequilibrium in finite populations. Genetics Research. 1970; 16(2):165–177.

Otto SP, Barton NH. The evolution of recombination: removing the limits to natural selection. Genetics. 1997; 147(2):879–906.

O’Reilly PF, Birney E, Balding DJ. Confounding between recombination and selection, and the Ped/Pop method for detecting selection. Genome research. 2008; 18(8):1304–1313.

Parsch J, Meiklejohn CD, Hartl DL. Patterns of DNA sequence variation suggest the recent action of positive selection in the janus-ocnus region of Drosophila simulans. Genetics. 2001; 159(2):647–657.

Pascanu R, Mikolov T, Bengio Y. On the difficulty of training recurrent neural networks. In: International conference on machine learning; 2013. p. 1310–1318.

Pool JE, Corbett-Detig RB, Sugino RP, Stevens KA, Cardeno CM, Crepeau MW, Duchen P, Emerson JJ, Saelao P, Begun DJ, Langley CH. Population Genomics of Sub-Saharan Drosophila melanogaster: African Diversity and Non-African Admixture. PLOS Genetics. 2012 12; 8(12):1–24. https://doi.org/10.1371/journal.pgen.1003080, doi: 10.1371/journal.pgen.1003080.

Price AL, Tandon A, Patterson N, Barnes KC, Rafaels N, Ruczinski I, Beaty TH, Mathias R, Reich D, Myers S. Sensitive detection of chromosomal segments of distinct ancestry in admixed populations. PLoS genetics. 2009; 5(6):e1000519.

Przeworski M, Wall JD. Why is there so little intragenic linkage disequilibrium in humans? Genetics Research. 2001; 77(2):143–151.

R Core Team. R: A Language and Environment for Statistical Computing. R Foundation for Statistical Computing, Vienna, Austria; 2018, https://www.R-project.org.

Rieseberg LH. Chromosomal rearrangements and speciation. Trends in ecology & evolution. 2001; 16(7):351–358.

Ritz KR, Noor MA, Singh ND. Variation in recombination rate: adaptive or not? Trends in Genetics. 2017; 33(5):364–374.

Rogers AR. How population growth affects linkage disequilibrium. Genetics. 2014; 197(4):1329–1341.

Russakovsky O, Deng J, Su H, Krause J, Satheesh S, Ma S, Huang Z, Karpathy A, Khosla A, Bernstein M, Berg AC, Fei-Fei L. ImageNet Large Scale Visual Recognition Challenge. Int J Comput Vision. 2015 Dec; 115(3):211–252. http://dx.doi.org/10.1007/s11263-015-0816-y, doi: 10.1007/s11263-015-0816-y.

Schiffels S, Durbin R. Inferring human population size and separation history from multiple genome sequences. Nature Genetics. 2014 06; 46:919 EP –. https://doi.org/10.1038/ng.3015.

Schrider DR, Ayroles J, Matute DR, Kern AD. Supervised machine learning reveals introgressed loci in the genomes of Drosophila simulans and D. sechellia. PLoS genetics. 2018; 14(4):e1007341.

Schrider DR, Kern AD. Supervised Machine Learning for Population Genetics: A New Paradigm. Trends in Genetics. 2018 Apr; 34(4):301–312. https://doi.org/10.1016/j.tig.2017.12.005, doi: 10.1016/j.tig.2017.12.005.

Schrider DR, Mendes FK, Hahn MW, Kern AD. Soft shoulders ahead: spurious signatures of soft and partial selective sweeps result from linked hard sweeps. Genetics. 2015; 200(1):267–284.

Schultz J, Redfield H. Interchromosomal effects on crossing over in Drosophila. In: Cold Spring Harbor symposia on quantitative biology, vol. 16 Cold Spring Harbor Laboratory Press; 1951. p. 175–197.

Schumer M, Xu C, Powell DL, Durvasula A, Skov L, Holland C, Blazier JC, Sankararaman S, Andolfatto P, Rosenthal GG, Przeworski M. Natural selection interacts with recombination to shape the evolution of hybrid genomes. Science. 2018; 360(6389):656–660. https://science.sciencemag.org/content/360/6389/656, doi: 10.1126/science.aar3684.

Singh ND, Stone EA, Aquadro CF, Clark AG. Fine-scale heterogeneity in crossover rate in the garnet-scalloped region of the Drosophila melanogaster X chromosome. Genetics. 2013; 194(2):375–387.

Slatkin M. Linkage disequilibrium in growing and stable populations. Genetics. 1994; 137(1):331–336.

Smith JM, Haigh J. The hitch-hiking effect of a favourable gene. Genetics Research. 1974; 23(1):23–35.

Srivastava N, Hinton G, Krizhevsky A, Sutskever I, Salakhutdinov R. Dropout: a simple way to prevent neural networks from overfitting. The journal of machine learning research. 2014; 15(1):1929–1958.

Sturtevant A. A case of rearrangement of genes in Drosophila. Proceedings of the National Academy of Sciences of the United States of America. 1921; 7(8):235.

Sutskever I, Vinyals O, Le QV. Sequence to Sequence Learning with Neural Networks. In: Proceedings of the 27th International Conference on Neural Information Processing Systems - Volume 2 NIPS’14, Cambridge, MA, USA: MIT Press; 2014. p. 3104–3112. http://dl.acm.org/citation.cfm?id=2969033.2969173.

Szegedy C, Liu W, Jia Y, Sermanet P, Reed SE, Anguelov D, Erhan D, Vanhoucke V, Rabinovich A. Going deeper with convolutions. In: IEEE Conference on Computer Vision and Pattern Recognition, CVPR 2015, Boston, MA, USA, June 7-12, 2015; 2015. p. 1–9. https://doi.org/10.1109/CVPR.2015.7298594, doi: 10.1109/CVPR.2015.7298594.

Tennessen JA, Bigham AW, O’Connor TD, Fu W, Kenny EE, Gravel S, McGee S, Do R, Liu X, Jun G, Kang HM, Jordan D, Leal SM, Gabriel S, Rieder MJ, Abecasis G, Altshuler D, Nickerson DA, Boerwinkle E, Sunyaev S, et al. Evolution and Functional Impact of Rare Coding Variation from Deep Sequencing of Human Exomes. Science. 2012; 337(6090):64–69. https://science.sciencemag.org/content/337/6090/64, doi: 10.1126/science.1219240.

Terhorst J, Kamm JA, Song YS. Robust and scalable inference of population history from hundreds of unphased whole genomes. Nature Genetics. 2016 12; 49:303 EP –. https://doi.org/10.1038/ng.3748.

Vincent P, Larochelle H, Bengio Y, Manzagol PA. Extracting and Composing Robust Features with Denoising Autoencoders. In: Proceedings of the 25th International Conference on Machine Learning ICML ‘08, New York, NY, USA: ACM; 2008. p. 1096–1103. http://doi.acm.org/10.1145/1390156.1390294, doi: 10.1145/1390156.1390294.

Wakeley J. Using the variance of pairwise differences to estimate the recombination rate. Genetics Research. 1997; 69(1):45–48.

Wall JD. A comparison of estimators of the population recombination rate. Molecular Biology and Evolution. 2000; 17(1):156–163.

Wang RJ, Gray MM, Parmenter MD, Broman KW, Payseur BA. Recombination rate variation in mice from an isolated island. Molecular ecology. 2017; 26(2):457–470.

White MJD. Animal cytology and evolution. CUP Archive; 1977.

Winckler W, Myers SR, Richter DJ, Onofrio RC, McDonald GJ, Bontrop RE, McVean GA, Gabriel SB, Reich D, Donnelly P, et al. Comparison of fine-scale recombination rates in humans and chimpanzees. Science. 2005; 308(5718):107–111.

Wiuf C. On the minimum number of topologies explaining a sample of DNA sequences. Theoretical population biology. 2002; 62(4):357–363.

Yi X, Liang Y, Huerta-Sanchez E, Jin X, Cuo ZXP, Pool JE, Xu X, Jiang H, Vinckenbosch N, Korneliussen TS, Zheng H, Liu T, He W, Li K, Luo R, Nie X, Wu H, Zhao M, Cao H, Zou J, et al. Sequencing of 50 human exomes reveals adaptation to high altitude. Science (New York, NY). 2010 07; 329(5987):75–78. https://www.ncbi.nlm.nih.gov/pubmed/20595611, doi: 10.1126/science.1190371.

Zickler D, Kleckner N. Recombination, pairing, and synapsis of homologs during meiosis. Cold Spring Harbor perspectives in biology. 2015; 7(6):a016626.

